# Signal Antagonists Suppress *Pseudomonas syringae* Pathogenicity

**DOI:** 10.1101/2024.06.10.598192

**Authors:** Josep Mas-Roselló, Anugraha Mathew, Veronika Avramenko, Jiajun Ren, Till Steiner, Simon Sieber, Leo Eberl, Karl Gademann

## Abstract

The bacterial plant pathogen *Pseudomonas syringae* causes significant damage to economically important crops worldwide. These bacteria coordinate their behavior and virulence through specific signaling compounds, such as the diazeniumdiolate leudiazen. Conventional antibacterial treatments enable the development of resistant strains. A more attractive treatment strategy would involve antagonists that suppress the expression of virulence factors without killing the pathogen, potentially reducing the risk of resistance development. Herein, we present the design and synthesis of analogs of leudiazen, which positively regulates the production of mangotoxin in *P. syringae* pv. *syringae* (Pss). Several compounds display inhibitory activity towards mangotoxin production, and a lead compound abolishes necrosis in infected tomato leaves, without significantly affecting bacterial growth. Thus, this study represents a promising advance towards developing effective and sustainable methods for bacterial disease control.

## Introduction

*Pseudomonas syringae* is a major plant pathogen that poses a significant threat to agriculture by infecting important crop species.^1,2^ Disease outbreaks caused by *P. syringae* can lead to disastrous economic consequences, such as the bacterial canker of kiwifruit initiated by *P. syringae* pv. *actinidiae* in New Zealand. This outbreak, which started in 2010, resulted in a loss of exports exceeding $500 million in less than five years.^3^ Another important pathovar is *P. syringae* pv. *syringae* (Pss), which is responsible for bacterial apical necrosis (BAN) in mango trees,^4,5^ as well as several other fruit tree diseases.^6^ Treating BAN remains challenging due to the limited efficacy of current control methods, such as traditional copper-based bactericides like the Bordeaux mixture (BM),^7,8^ which are associated with the development of copper resistant strains^9,10^ and the risk of copper bioaccumulation.^11^

Pss employs diverse virulence strategies for host colonization and disease progression,^1,12^ including a type III secretion system that delivers diverse effector proteins as well as the production of phytotoxins, exopolysaccharides, cell wall-degrading enzymes, and plant hormones. Among them, the antimetabolite toxin, mangotoxin, is the major virulence factor of Pss isolated from mango trees (Figure 1A).^4^ Despite its significance, the chemical structure of mangotoxin remains unknown, with studies being mostly limited to the genomic level.^13,14^ The virulence of Pss is linked to the *mgo* operon, which was shown to encode an unknown signaling molecule.^15–17^ Recently, we identified **leudiazen** as the signaling molecule regulating mangotoxin production (Figure 1B).^18^ Leudiazen belongs to a unique class of signaling molecules containing a diazeniumdiolate functional group like the prototypical quorum-sensing signal valdiazen.^19–21^ We previously demonstrated that the strategic degradation of leudiazen using potassium permanganate (KMnO_4_), compatible with organic farming, significantly decreases the virulence of Pss in infected tomato leaf models without negatively impacting bacterial growth.^18^

**Figure 1.**
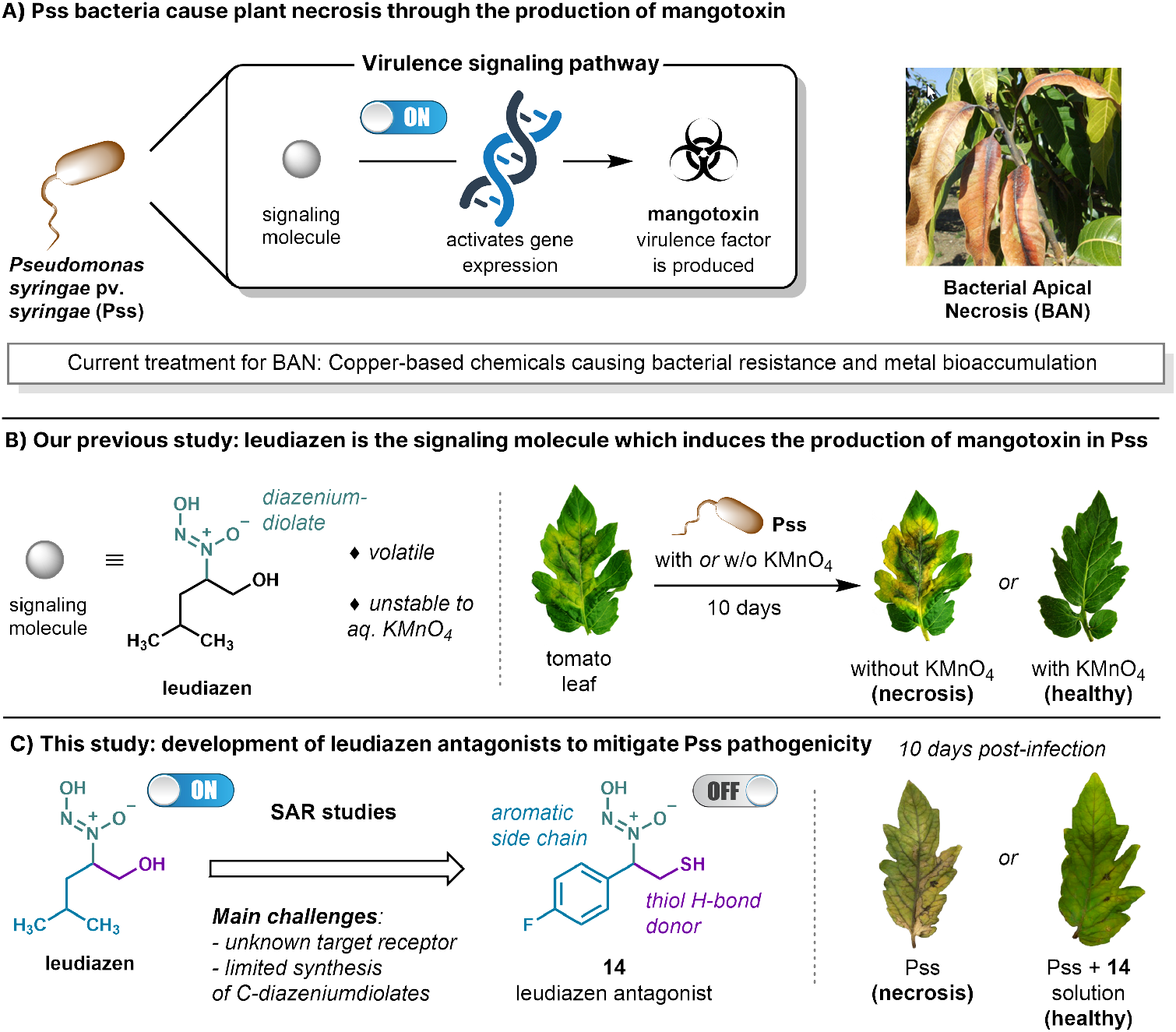
A) Virulence regulation in *Pseudomonas syringae* pv. *syringae* (Pss) mediated by a signaling molecule (image cropped from original in ref. 5 (license CC BY)); B) Our prior work showing that the signaling molecule is leudiazen, and that its chemical degradation with KMnO_4_ attenuates Pss virulence (leaf images taken from ref. 18 (license CC BY-NC)); C) Novel approach to mitigate Pss virulence with a synthetic inhibitor (**this study**).

Interfering with the virulence signaling pathway has emerged as an attractive strategy, aiming to attenuate virulence rather than killing the pathogen, thereby potentially reducing the risks of bacterial resistance.^22–27^ In this context, a potentially effective and sustainable approach involves using small-molecule synthetic inhibitors or antagonists to disrupt the signalling cascade.^28–36^ In principle, antagonists could be designed to specifically target the signaling pathways involved in virulence, providing a more targeted approach compared to the chemical degradation of signaling molecules with broad-spectrum treatments, such as KMnO_4_ in our previous study.^18^ However, while several inhibitors of *N*-acylated L-homoserine lactones (AHLs)^37^ − the most common signal class in gram-negative bacteria − have been reported,^38–41^ inhibitors of less common signaling molecule classes remain understudied. Therefore, research in this direction is essential to control the virulence of pathogens with diverse signaling pathways.^42^

We speculated that synthetic analogs of leudiazen could mitigate Pss virulence by disrupting its signaling pathway, thereby reducing the production of mangotoxin (Figure 1C). To achieve this goal, we anticipated two main challenges: i) the target receptor of leudiazen is unknown, limiting the rational design of potential antagonist derivatives; ii) the chemistry of diazeniumdiolates is relatively underdeveloped,^43,44^ potentially limiting the diversity of chemically tractable derivatives. Herein, we present an efficient synthesis of *C*-diazeniumdiolates, enabling access to a broad range of analogs. Systematic phenotypic screening^45^ revealed key structural features responsible for their biological activity. Several compounds exhibited inhibitory activity towards mangotoxin production, and a lead compound effectively abolished necrosis in infected tomato leaves, without significantly affecting bacterial growth.

## Results and Discussion

### Design and synthesis of *N*-alkyl diazeniumdiolates

To develop antagonists for leudiazen, identifying the structural components responsible for its bioactivity as an inducer of mangotoxin expression in Pss is crucial. Previous studies on structurally related diazeniumdiolates, such as the antifungal fragin,^20^ revealed three key components playing an important role in bioactivity: the side chain (colored blue in Figure 1C), the diazeniumdiolate functional group (colored green), and the methylenehydroxy part (coloured purple). Thus, in a first step we performed a phenotypic screening of leudiazen analogs by sequentially modifying these three parts.

Regarding the side chain, leudiazen contains a leucine-derived *iso*-butyl group with lipophilic character. Interestingly, leudiazen is slightly volatile due to its low molecular weight, a feature that may facilitate long-distance communication for bacteria on plant surfaces.^18^ To explore modifications, we aimed to replace the *iso*-butyl group with smaller and bigger alkyl groups, potentially modulating volatility, and also to test heavier, more lipophilic aromatic substituents. However, general efficient synthetic routes to access diazeniumdiolate analogs were missing in the literature. While the biosynthesis of some natural diazeniumdiolates has been investigated,^19,20,47,50–52,54–60^ only a limited number of synthetic derivatives have been described.^18–20,61^ In the case of leudiazen, the previous synthesis in 4 steps from DL-**leucinol** yielded only a 15% overall yield (Scheme 1A),^12^ being far from efficient. Additionally, 1,2-amino alcohol starting materials have restricted availability. Mainly those derived from α-amino acids are commercial, limiting the choice of side chain substituents.

**Scheme 1.**
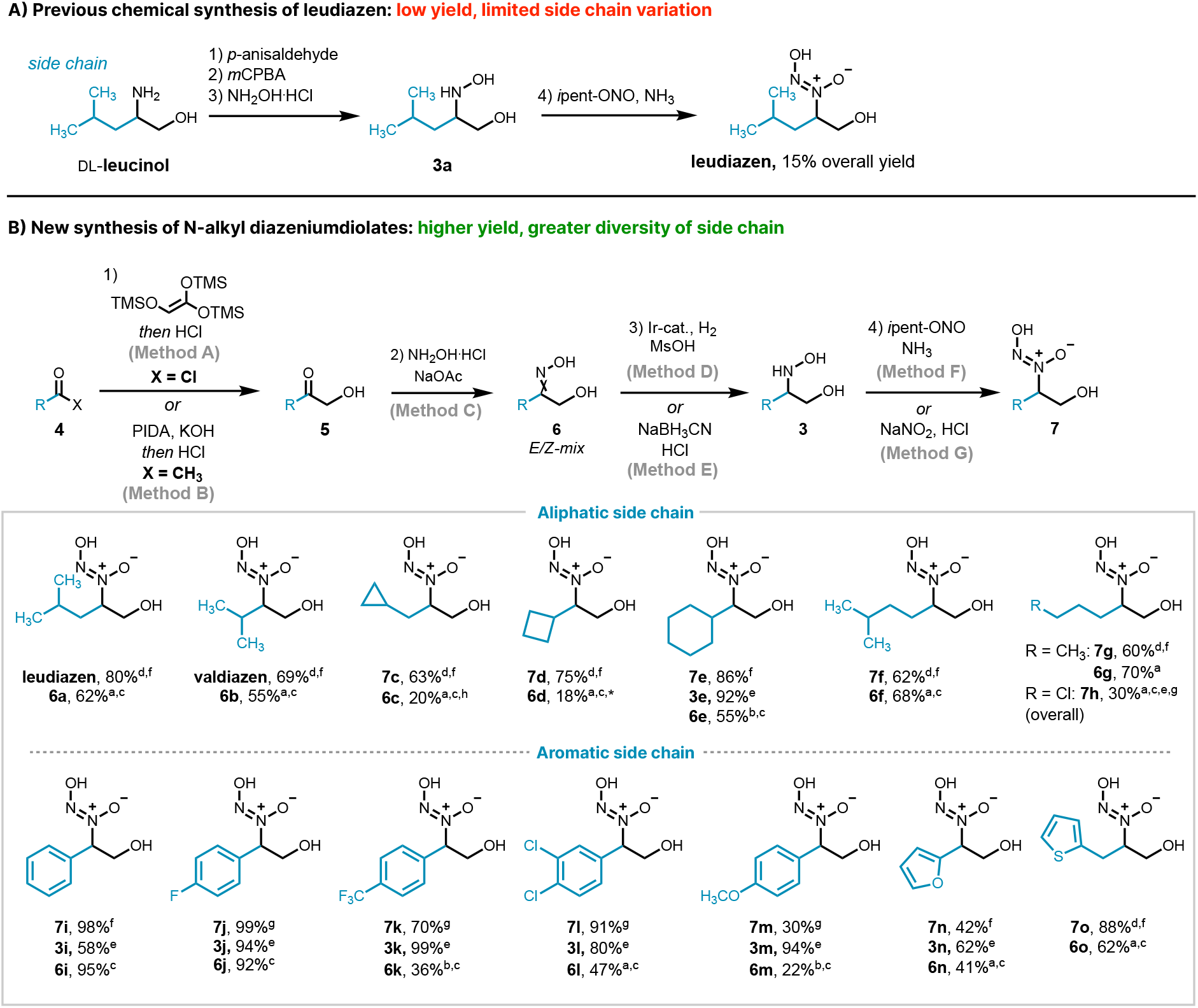
Syntheses of *N*-alkyl diazeniumdiolates. A) Our previous synthesis of leudiazen from DL-leucinol, resulting in low overall yield; B) A more efficient synthesis of leudiazen and other diverse diazeniumdiolate compounds developed in this study. For full experimental details, see the supporting information document. *Diminished yield due to volatility of the α-hydroxyketone intermediate **5c**.

The newly developed synthetic route involves 4 steps starting from readily available and inexpensive acyl chlorides or methyl ketones **4** (Scheme 1B). From these, α-hydroxy ketones **5** were prepared *via* the Wissner reaction (X = Cl)^62,63^ or *via* α-hydroxylation of methyl ketones **4** (X = CH_3_).^64^ Subsequently, oxime condensation between crude intermediates **5** and hydroxylamine yielded oxime substrates **6** - typically as mixtures of C=N double bond *E/Z* diastereoisomers - in good overall yields. Reduction of the oximes **6** gave hydroxylamines **3**. The use of an acid-mediated Ir-catalyzed hydrogenation protocol^65,66^ to prepare *volatile* aliphatic hydroxylamine derivatives (e.g. **3a-d**) facilitated their isolation as the crude methanesulfonate salts, without need for work-up or extra purification steps. For heavier hydroxylamines (e.g. **3i-n**), NaBH_3_CN was used as stoichiometric reductant.^67^ Finally, established *N-*nitrosation of hydroxylamines **3** under basic^68^ or acidic conditions^69^ provided the target *C*-diazeniumdiolate compounds **7** as racemates in high yields.

Through this route, a wide range of diazeniumdiolate analogs bearing both aliphatic and aromatic side chain (R) substituents were accessed. Within the first class, linear (**7g**,**h**), branched (**leudiazen, valdiazen, 7f**), and cyclic derivatives having different ring sizes (**7c-e**) could be prepared. Notably, a 50% overall yield of leudiazen was achieved, significantly improving the 15% overall yield obtained previously.^18^ The mild nature of the current synthesis was showcased by obtaining analog **7h**, bearing a reactive primary alkyl chloride. Concerning aromatic derivatives, electron-rich (**7m**), neutral (**7i**), and electron-deficient (**7j-l**) phenyl rings were compatible. Furthermore, high yields were also obtained for derivatives **7n** and **7o** containing furane and thiophene electron-rich heterocycles, respectively.

### Structure-activity-relationship (SAR) studies

With a series of leudiazen derivatives in hand, β-galactosidase assays were performed to assess their ability to competitively inhibit the promoter activity of the mangotoxin gene cluster (*mbo*), by employing a respective promoter-reporter transcriptional fusion (Figure 2).

**Figure 2.**
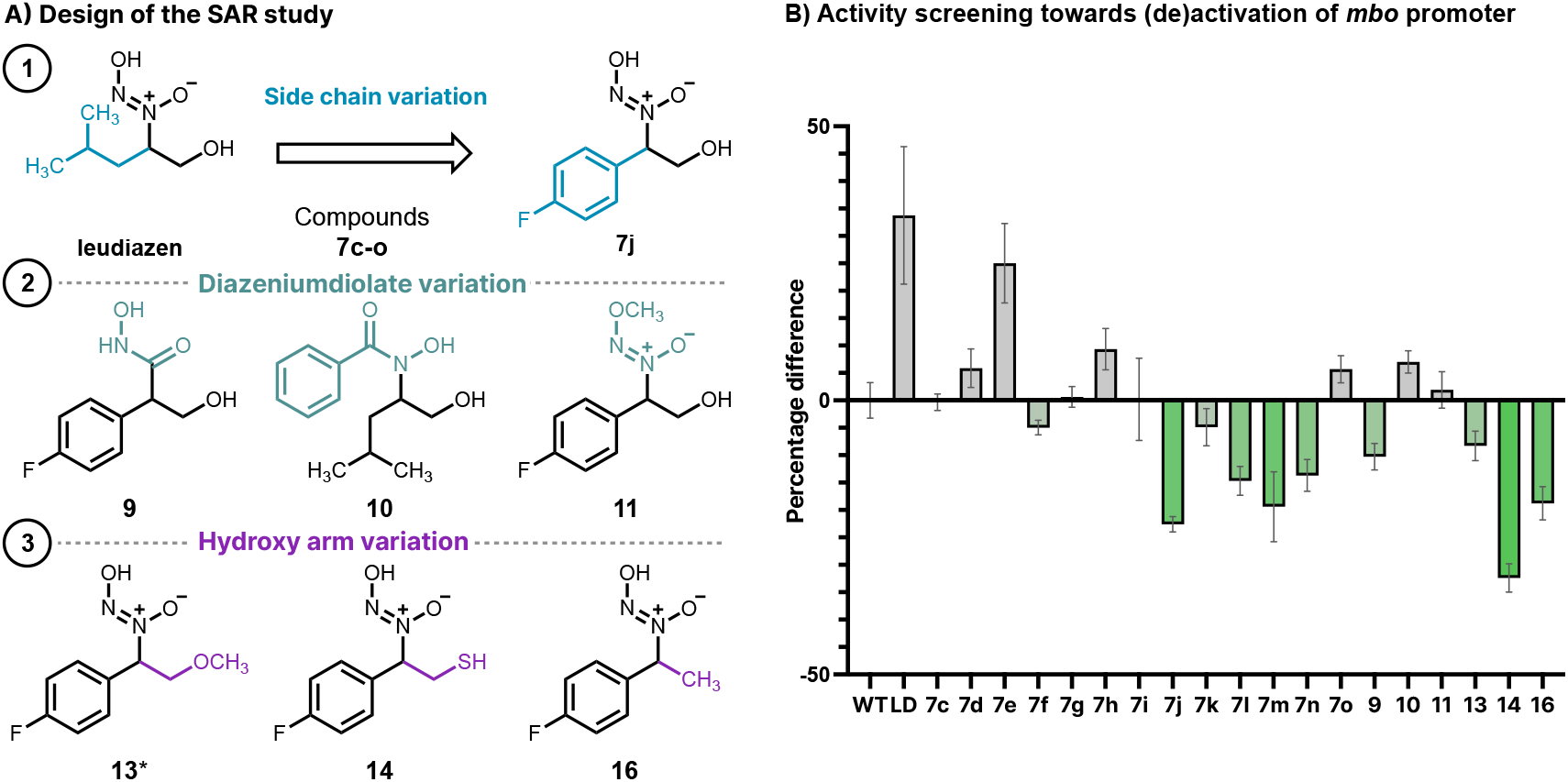
**Competitive inhibition of the mangotoxin promoter by diazeniumdiolate derivatives**, including **A**) a workflow of the SAR studies, and **B**) the activities of the *mboA* promoter in the presence of the synthesized compounds (20 μM) relative to the wild-type Pss. Experimental and synthesis details on compounds **9-11**,**13-16** included in the supporting information document. *Isolated as the potassium salt. The abbreviations in the figure are WT for Pss and LD for leudiazen.

None of the derivatives bearing an aliphatic side chain showed significant inhibitory activity (**7c-h**). By contrast, the cyclohexyl derivative **7e** and the chloropropyl compound **7h** exhibited leudiazen-like induction activity, possibly for distinct reasons: for **7e**, a steric match may be responsible, as its cyclohexyl side chain is similar in size to leudiazen’s isobutyl substituent. On the other hand, the promoting activity of **7h** could be attributed to covalent binding to an unknown target receptor, facilitated by its primary chloride electrophilic handle. Supporting this hypothesis, analog **7g** lacking the chlorine atom showed no significant increase of activity. In stark contrast, some analogs bearing an aromatic side chain displayed promising inhibitory activity (i.e. **7j, 7m, 7n, 14, 16**). The aromatic vs non-aromatic nature of the side chain clearly has a major impact, as seen in the opposite activities of analogs **7h** (R = cyclohexyl) and **7i** (R = phenyl). However, there is no general trend in terms of electronic character, as both electron-rich (**7m, 7n**) and electron-deficient rings (**7j**) exhibited comparable levels of inhibitory activity. From this screening, fluorophenyl derivative **7j** emerged as the most promising candidate. Noteworthy, no significant difference in the activities between the (*R*) or (*S*) enantiomer of **7j** was observed at 20 μM concentration (Figure S1 in the supporting information), in line with the previous findings with leudiazen.^18^

Subsequently, in addition to the side chain, the other structural features of the compounds that *a priori* may be important for bioactivity were tested, such as aforementioned the diazeniumdiolate group. To date, only few natural products containing this motif have been isolated,^46–52^ and their biological activity is commonly attributed to the strong metal-binding ability conferred by the diazeniumdiolate group.^19,49,53,56^ Compounds **9** and **10**, containing a hydroxamate group also known to have metallophore ability,^70,71^ instead of the diazeniumdiolate, showed little to no inhibitory activity. Compound **11**, a version of the antagonist **7j** with a masked diazeniumdiolate group methylated at the diazeniumdiolate O^2^-position,^43^ exhibited no inhibitory activity and even had a slightly promoting effect, demonstrating the importance of the diazeniumdiolate group for antagonistic activity.

Lastly, the hydroxy arm was varied keeping the diazeniumdiolate group unmasked (**13-16**). For instance, compound **16**, a close analog of the hit compound **7j** lacking the hydroxy substituent, showed only a minor reduction in the antagonistic activity, while methylation of the hydroxy group (as in **13**), eliminating its hydrogen-bond donating ability, largely suppressed the activity. Remarkably, replacement by thiol, a better hydrogen-bond donor, increased the inhibitory activity, rendering fluoroaryl-thio lead compound **14** as the most promising leudiazen antagonist for subsequent biological studies.

### Suppressing virulence through chemical antagonism

The lead compound **14** was further tested to evaluate its efficacy in suppressing Pss virulence through inhibition of mangotoxin production. In plate bioassays using an *Escherichia coli* (E. coli) indicator strain (Figure 3A), the wild type Pss exhibited antimicrobial activity through the production of mangotoxin. However, in the presence of 20 μM of **14**, the antimicrobial activity was abolished, indicating a substantial decrease in mangotoxin production. As a control, a Pss Δ*mgoA* mutant background (ΔmgoA) – which does not produce mangotoxin – was used, both with and without **14**; in none of the two assays antimicrobial activity was observed, indicating that compound **14** cannot functionally complement **leudiazen**.

**Figure 3.**
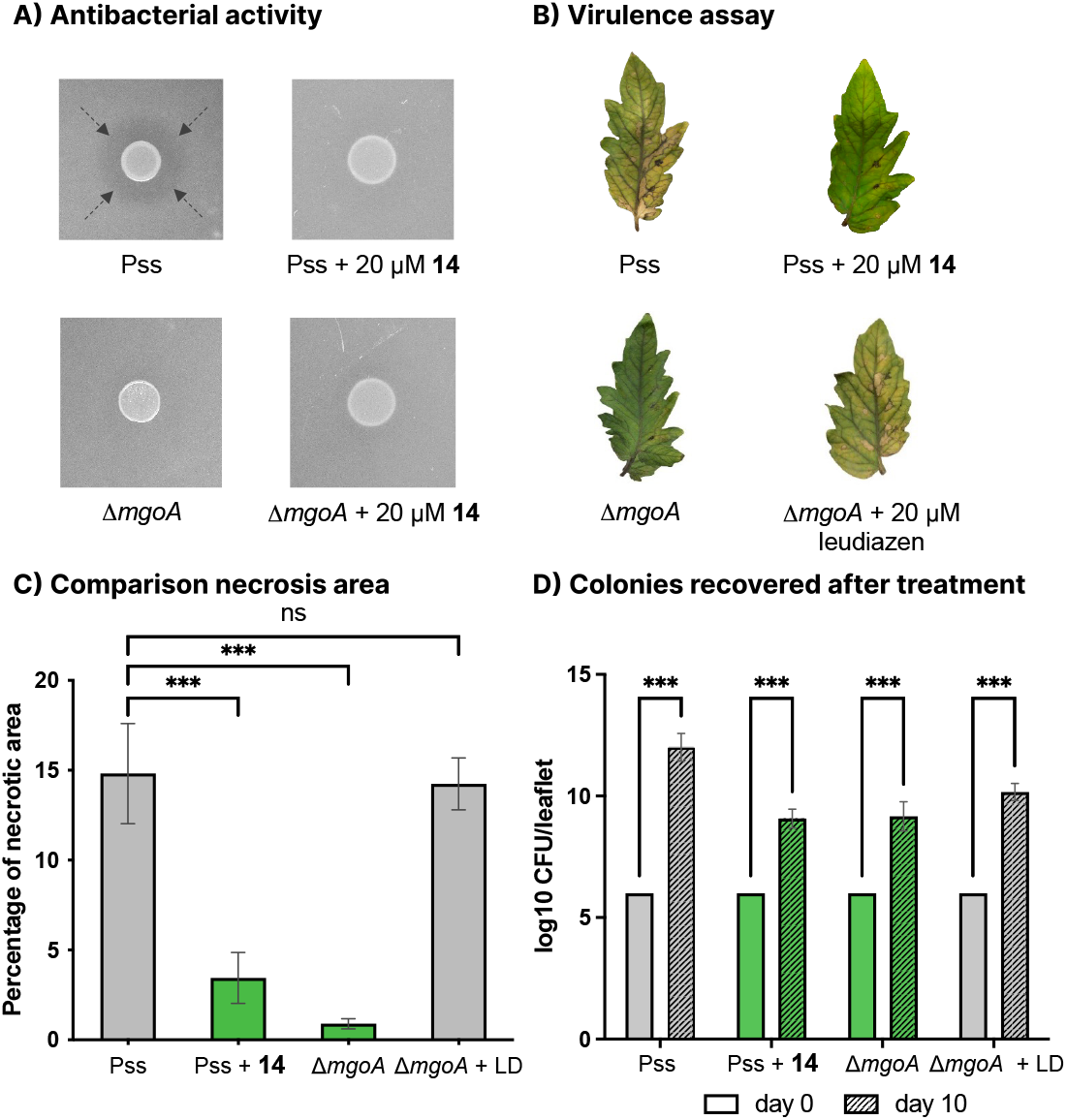
Competitive inhibition of mangotoxin production by the novel diazeniumdiolates. (A) Mangotoxin production assay using *E. coli* as an indicator. The halos, indicating mangotoxin production, are labelled with dashed arrows. (B) Virulence assay including: tomato leaflets infected with Pss and treated with compound **14** (Pss + **14**); leaflets infected with Pss without any treatment (Pss), as well as leaflets infected with ΔmgoA mutant (controls). Six leaflets per strain/condition, and three independent experiments were performed. All sets of leaflets were maintained for a period of 10 days and the development of necrotic symptoms was recorded. The concentration of compound **14** and leudiazen employed was 20 μM. Pictures were taken after 10 days postinfection (dpi). Photo Credit: Dr. Anugraha Mathew, University of Zurich. (C) Relative virulence of Pss, ΔmgoA, Pss + **14** and ΔmgoA + leudiazen in tomato leaflets measured by the % area of necrosis. Statistical analysis was performed using one-way ANOVA and Tukey’s multiple comparisons was employed as the posttest. Error bars represent the SEM (standard error of the mean) and ***P < 0.001. (D) Amount of *P. syringae* cells recovered from infected leaves at 0 and 10 dpi; the initial bacteria colonies per leaf were selected as 10^6^; error bars represent the SEM; the statistical analysis was performed using two-way ANOVA with the Bonferroni-Dunn method, and the p-values of the statistical analysis are reported according to the American Psychological Association style with *** ≤ 0.001. The abbreviation in the figure is LD for leudiazen.

Moreover, experiments were conducted to assess the impact of compound **14** in controlling infection on model tomato leaves (Figures 3B and 3C). An aqueous solution of compound **14** (20 μM) was applied to infected leaves for 20 seconds, followed by rinsing with water. After 10 days post-infection (dpi), untreated leaves (Pss) exhibited clear signs of necrosis, with approximately 15% of the leaf area affected, whereas treated leaves (Pss + **14**) showed less than 5% necrotic area, maintaining their healthy green coloration. As controls, the Δ*mgoA* mutant stayed healthy but developed necrosis in the presence of leudiazen. Importantly, mangotoxin production was inhibited by the chemical treatment without affecting bacterial growth, as evidenced by the increase in Pss cell recovery from both treated and untreated leaves after 10 dpi (Figure 3D).

Taken together, these findings clearly demonstrate that chemical inhibition of the mgo signaling system can effectively control Pss pathogenicity. Remarkably, the *mbo* gene cluster encoding mangotoxin biosynthesis genes, initially described in Pss,^15^ has been identified in many other P. *syringae* pathovars,^72,73^ as well as in strains of the genus *Serratia*.^74^ This suggests that chemical interference with diazeniumdiolate inhibitors holds the potential to become a general strategy to mitigate the virulence of a broader range of genetically-related bacterial pathogens.

## Conclusion

In summary, our study introduces a promising strategy for controlling *Pseudomonas syringae* pathogenicity by treatment with diazeniumdiolate analogs of leudiazen. Through systematic design and structure-activity-relationship studies, we identified lead compound **14** as an effective antagonist, significantly inhibiting mangotoxin production without impeding bacterial growth. Biological assays further demonstrated the efficacy of **14** in suppressing mangotoxin-induced phenotypes both *in vitro* and *in planta* models. This chemical inhibition of the mgo signaling system presents a compelling approach to mitigate Pss pathogenicity without killing the bacteria, thus potentially minimizing the risk of resistance development. Moreover, diazeniumdiolate inhibitors may be applicable to a broader spectrum of genetically-related bacterial pathogens. Thus, our findings may serve as a blueprint for future research in sustainable and effective disease control methods.

## Acknowledgments

We thank the Swiss National Science Foundation for funding parts of this work (186410).

## Authors contributions

K.G. and L.E. conceived and supervised the project. J.M.R. designed the compounds and synthesized most of them. A.M. designed and performed all biological assays. J.M.R., A.M. and S.S. analyzed the data. V.A. synthesized some of the compounds. J.J. resynthesized the lead compound **14**. T.S. conducted preliminary synthesis experiments. S.S. co-designed the biological assays and coordinated the synthesis and biological experiments. J.M.R. wrote the manuscript with substantial contributions from A.M., S.S. and K.G. All authors revised the manuscript.

