## Supporting information for "Signal Antagonists Suppress *Pseudomonas syringae* Pathogenicity"

**This PDF file includes:**

Materials and Methods
NMR Spectra
References

### Materials and Methods

General information

All chemicals were purchased from Acros, Fluka, or Sigma-Aldrich and were used without further purification. All reactions were carried out in flame-dried glassware (unless aqueous reagents were used) and reactions involving air sensitive compounds were performed under an argon or nitrogen atmosphere. Solvents applied for chemical transformations were either puriss quality or HPLC grade solvents, which had been dried by filtration through activated aluminium oxide under nitrogen ($H_2O$ content <10 ppm, *Karl-Fischer* titration). For work-up and purification, solvents were distilled from technical grade. All synthetic transformations were monitored either by thin layer chromatography (TLC) or $^1H$ NMR spectroscopy. Yields refer to purified, dried and spectroscopically pure compounds. TLC was performed on Merck silica gel 60 F254 plates (0.25 mm thickness) pre-coated with a fluorescent indicator, and visualized by staining with a potassium permanganate solution. Concentration under reduced pressure was performed by rotary evaporation at 40 °C. Flash chromatography was performed using silica gel 60 (230-400 mesh) from Sigma-Aldrich with a forced flow eluent at 0.1-0.3 bar pressure. All $^1H$ and $^{13}C$ NMR spectra were recorded using a Bruker Avance 400 MHz, 500 MHz, 600 MHz ($^1H$) & 101 MHz or 126 MHz ($^{13}C$) spectrometer at RT (unless otherwise stated). Chemical shifts (δ-values) are reported in ppm, spectra were calibrated relative to the residual proton chemical shifts (MeOD, δ = 3.31; $D_2O$, δ = 4.79) and the residual carbon chemical shifts (MeOD, δ = 49.0) of the solvents, multiplicity is reported as follows: s = singlet, d = doublet, t = triplet, q = quartet, m = multiplet or unresolved and coupling constant *J* in Hz. IR spectra were recorded on a Perkin Elmer SpectrumTwo ATR-FTIR. The absorptions are reported in $cm^{-1}$. All mass spectra (HRMS) were recorded by the Mass Spectrometric Service of the University of Zurich on a QExactive instrument (Thermo Fisher Scientific, Bremen, Germany) equipped with a heated electrospray (ESI) ionization source and connected to a Dionex Ultimate 3000 UHPLC system. Melting points (m.p.) were determined using a Büchi B-545 apparatus in open capillaries and are uncorrected. Optical rotations °[$\alpha$] were measured at the sodium D line using a 1 mL cell with a 1 dm path length on a Jasco P-2000 digital polarimeter and the concentration c is given in g/100 mL MeOH.

Syntheses of oxime intermediates (**6**):

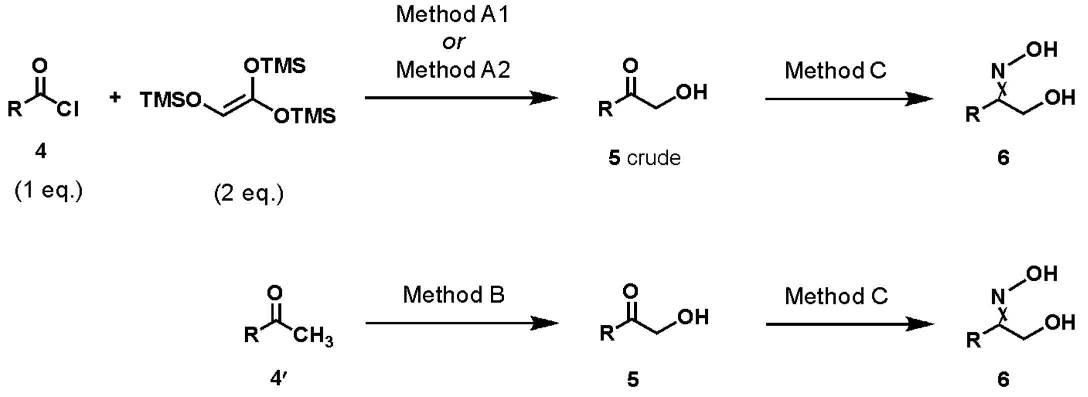


Method A1: neat, 90 °C, 16 h; *then* 1N HCl (aq.), 80 °C, 2 h
Method A2: $Et_3N$, THF, 100 °C, MW, 5 min; *then* 1N HCl (aq.), rt, 30 min
Method B: $PhI(OAc)_2$, KOH, MeOH, rt, 12 h; *then* 1N HCl (aq.), 80 °C, 2 h
Method C: $HONH_2 \cdot HCl$, NaOAc, EtOH, 60 °C, 6 h

Synthesis of diazeniumdiolates (**7**) from oxime intermediates (**6**):

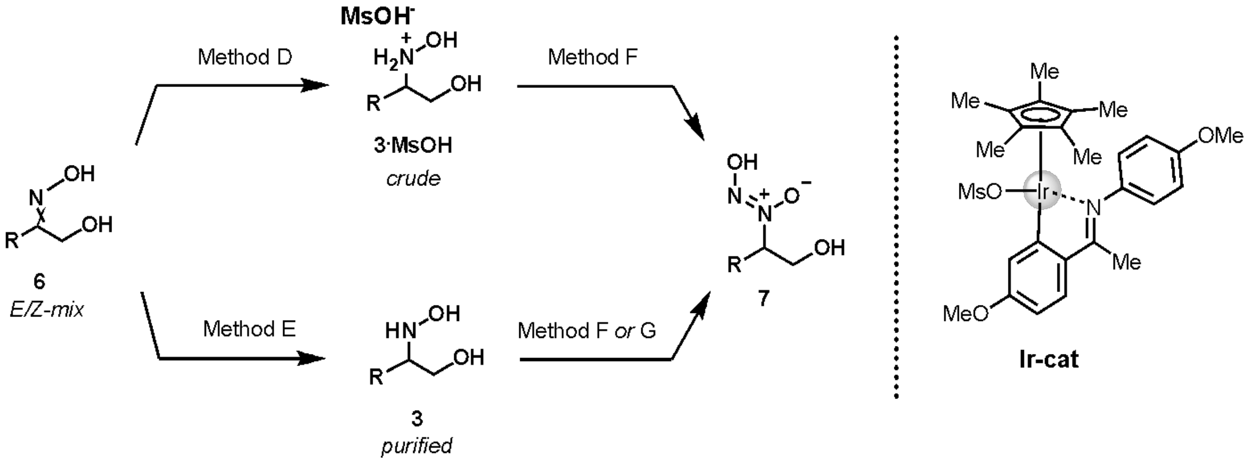


Method D: 1 mol% **Ir-cat**, 1.5 equiv MsOH, 50 bar $H_2$, *i*PrOH, rt, 16 h
Method E: $NaBH_3CN$, $HCl_{(aq.)}$, MeOH, rt, 30 min
Method F: *i*pentONO, $NH_3$, MeOH, 0 °C, 45 min
Method G: $NaNO_2$, $HCl_{(aq.)}$, EtOH, 0 °C, 30 min

General methods **A-G**:

**Method A1**: A Schlenk tube, equipped with a magnetic stirring bar, was charged with the acid chloride **4** (1 eq.) under an atmosphere of nitrogen. 1,1,2-Tris(trimethylsilyloxy)ethane (2 eq.) was added, the tube was sealed, and the reaction mixture was heated at 90 °C for 12 h. The reaction mixture was cooled to 50 °C. 1,4-Dioxane was added (0.8 mL/mmol of **4**) followed by a dropwise addition of 1 N aqueous HCl (0.2 mL/mmol of **4**) (CAUTION: vigorous bubbling generated). The resulting mixture was stirred at 80 °C for 2 h. After cooling to room temperature, the mixture was diluted with brine and extracted three times in EtOAc. The combined organic layers were washed with a saturated aqueous solution of $NaHCO_3$, dried over $Na_2SO_4$, filtered, and concentrated under reduced pressure to afford the crude hydroxyl ketone **5**, which was used in the next step without further purification.

**Method A2**: A flame-dried microwave reactor vessel, equipped with a magnetic stirring bar and cooled to -10 °C with a NaCl/ice/water bath, was charged successively with anhydrous THF (1 M), acid chloride **4** (1.0 eq.), tris(trimethylsiloxy)ethylene (1.1 eq.), and triethylamine (1.0 eq., dropwise addition) under an atmosphere of argon (white precipitate formed). The reaction mixture was stirred at -10 °C for additional 5 min. The mixture was heated to 100 °C for 5 min under microwave irradiation. After cooling to room temperature, a 1 M HCl aqueous solution (15 mL/mmol of **4**) was added (bubbling observed), and the resulting solution was stirred at ambient temperature for 30 min. The aqueous phase was saturated with NaCl and extracted three times in EtOAc. The combined organic layers were washed with a saturated aqueous solution of $NaHCO_3$, dried over $Na_2SO_4$, filtered, and concentrated under reduced pressure to afford the crude α-hydroxyl ketone **5**, which was used in the next step without further purification.

**Method B**: A flame-dried round-bottom flask, equipped with a magnetic stirring bar, was charged with the methyl ketone **4'** (1 eq.), anhydrous methanol (0.3 M) and KOH (4.5 eq.) at 0 °C under an atmosphere of nitrogen. $PhI(OAc)_2$ (1.2 eq.) was added portion-wise and the reaction mixture was stirred for 10 min before allowing it to warm to room temperature. After stirring for 3 h, the reaction mixture was concentrated under reduced pressure, diluted in $H_2O$, and the product extracted three times in EtOAc. The combined organic layers were washed with brine, dried over $Na_2SO_4$, filtered, and concentrated under reduced pressure. The residue obtained was dissolved in methanol (1 M) and a 3 M aqueous HCl solution (1 mL/mmol of **4'**) was added dropwise. The mixture was stirred overnight at room temperature. The MeOH solvent was removed under reduced pressure and the aqueous phase was extracted three times in EtOAc. The combined organic extracts were washed with a saturated $NaHCO_3$ aqueous solution, dried over $Na_2SO_4$, filtered, and concentrated under reduced pressure. The crude α-hydroxyl ketone **5** was purified by silica gel flash column chromatography.

**Method C**: A teflon-capped tube, equipped with a magnetic stirring bar, was charged with the crude ketone **5** (1.0 eq.), EtOH (0.5 M), hydroxylamine hydrochloride (2.0 eq.) and sodium acetate (3.0 eq.). The reaction mixture was stirred at 60 °C for 6 h. After cooling to room temperature, the solvent was removed under reduced pressure. The resulting solid residue was diluted with brine and extracted three times in EtOAc. The combined organic layers were washed with brine, dried over $Na_2SO_4$, filtered, and concentrated under reduced pressure. The crude oxime product **6** was purified by silica gel flash column chromatography and isolated as mixtures of *E/Z* diastereoisomers (unless otherwise stated).

**Method D**: Without protective precaution from air and moisture, a test tube equipped with a magnetic stirring bar was charged with the iridium catalyst **Ir-cat**[1] (1 mol%) and the oxime substrate **6** (1 eq.). The tube was sealed with a septum cap and an atmosphere of nitrogen was created. Anhydrous 2-propanol (1.0 M) and methanesulfonic acid (1.5 eq.) were added at room temperature. The septum was pierced with a needle and the test tube was placed in a high-pressure reactor. The reactor was purged with argon (3 x 5 bar) then hydrogen (3 x 5 bar), pressurized to 50 bar of hydrogen and the reaction mixture was stirred at room temperature for 16 h. Hydrogen was released and the solvent removed under reduced pressure to afford the corresponding hydroxylammonium methanesulfonate product salt **3·MsOH** without need for further purification (typically >95% purity by NMR).

**Method E**: A round-bottom flask, equipped with a magnetic stirring bar, was charged with the oxime substrate **6** (1.0 eq.), and methanol (1.0 M). A minimum amount of methyl orange indicator was added, followed by sodium cyanoborohydride (1.5 eq.). The reaction vessel was capped with a rubber septum and flushed with nitrogen. A 3 M solution of hydrochloric acid in MeOH was added dropwise (colour changes from orange to violet upon sufficient acid addition). After 30 min of the reaction remaining violet, the solvent was removed under reduced pressure, The residue was diluted with a saturated aqueous solution of $Na_2CO_3$ and extracted three times in EtOAc. The combined organic layers were dried over $Na_2SO_4$, filtered and the solvent was removed under reduced pressure. The crude hydroxylamine product **3** was purified by flash column chromatography, eluting in EtOAc:MeOH 95:5 unless otherwise stated.

**Method F**: A round-bottom flask, equipped with a magnetic stirring bar, was charged with the pure hydroxylamine substrate **3** (1 eq.). An atmosphere of argon was created and a 7 N solution of ammonia in MeOH (2.5 mL/mmol of **3**) was added at 0 °C. Isoamyl nitrite (3 eq.) was added dropwise and the reaction mixture was stirred at 0 °C for 45 min. The reaction mixture was concentrated under reduced pressure. The residue was dissolved in $H_2O$ and the solution was passed through a Discovery® DSC-18 SPE tube (Supelco), eluting with more $H_2O$ (e.g. 3 tube volumes). The solvent was concentrated under reduced pressure, and the residual $H_2O$ removed by two cycles of dissolving in MeOH followed by concentration under reduced pressure, to afford the pure diazeniumdiolate **7**. [**Note**: when starting from the hydroxylammonium methanesulfonate product salt **3·MsOH**, an extra purification step consisting of trituration with *n*-pentane:$Et_2O$ 1:1 is needed (after DSC-18 SPE filtration and concentration) to separate the diazeniumdiolate product **7** from the $MsONH_4$ precipitate].

**Method G**: A round-bottom flask, equipped with a magnetic stirring bar, was charged with the hydroxylamine substrate **3** (1 eq.). EtOH and $H_2O$ were added (1:1 v/v, 0.25 M) and the solution was cooled to 0 °C. 1 N aqueous HCl (1.05 eq.) was added, and the system was degassed by bubbling argon under vigorous stirring for 5 min. Separately, a solution of $NaNO_2$ (1.05 eq) in $H_2O$ (1 M) was prepared and degassed likewise. The $NaNO_2$ aqueous solution was added dropwise over 5 min at 0 °C. The reaction was stirred for 30 min at 0 °C. The resulting yellow solution was diluted with $H_2O$, brine, and extracted three times in EtOAc. [Note: the pH of the aqueous phase should be approximately 5; below pH 3 the product slowly decomposes]. The combined organic layers were dried over $Na_2SO_4$, filtered and the solvent was removed under reduced pressure to afford the diazeniumdiolate **7** without need for further purification (unless otherwise stated).

*(E/Z)-1-hydroxy-4-methylpentan-2-one oxime* **(6a)**

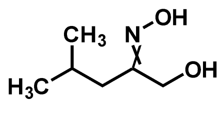


*(E/Z)*-**6a**

Following methods A1 and C, **6a** was prepared from commercial isovaleryl chloride (0.91 mL, 7.50 mmol). *(E/Z)*-**6a**: colourless solid (610 mg, 62% overall yield); $\mathbf{R}_f$ ($SiO_2$; *n*-pent:EtOAc 40:60) = 0.40, 0.25 (two C=N double bond isomers) (non-UV active, stained with $KMnO_4$); **IR** (neat, $cm^{-1}$): $\nu_{max}$ = 3233.4, 2958.0, 2928.0, 1463.4, 1369.0, 1293.7, 1166.6, 1092.2, 987.9, 944.8; **$^1$H NMR** (400 MHz, $CD_3CN$) (mixture of two C=N double bond isomers in a 93:7 ratio) δ 4.30 (s, 2H, *minor*), 4.01 (s, 2H, *major*), 2.24 (d, *J* = 7.5 Hz, 4H, *major+minor*), 2.12 – 1.96 (m, 2H, *major+minor*), 0.90 (d, *J* = 6.6 Hz, 12H, *major+minor*); **$^{13}$C NMR** (101 MHz, $CD_3CN$) δ 164.87 (*minor*), 160.14 (*major*), 63.76 (*major*), 57.58 (*minor*), 40.43 (*minor*), 34.53 (*major*), 26.62 (*minor*), 26.43 (*major*), 23.09 (*major*), 22.79 (*minor*); **HRMS** (ESI+): m/z calc. for $C_6H_{14}NO_2$ $[M+H]^+$ 132.10191, found 118.10176.

*(E/Z)-1-hydroxy-3-methylbutan-2-one oxime* **(6b)**

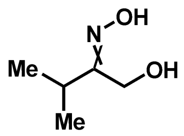


*(E/Z)*-**6b**

Following methods A1 and C, **6b** was prepared from commercial acid chloride (712 mg, 6.70 mmol). *(E/Z)*-**6b**: colourless solid (430 mg, 55% overall yield); $\mathbf{R}_f$ ($SiO_2$; *n*-pent:EtOAc 65:35) = 0.45, 0.3 (two C=N double bond isomers) (non-UV active, stained with $KMnO_4$); **IR** (neat, $cm^{-1}$): $\nu_{max}$ = 3201, 2976, 1461, 1042, 989, 945, 752, 685; **$^1$H NMR** (400 MHz, $CD_3CN$) (mixture of two C=N double bond isomers in a 90:10 ratio) δ 4.30 (s, 2H, *minor*), 4.09 (s, 2H, *major*), 3.22 (hept, *J* = 7.1 Hz, 1H, *major*), 2.77 – 2.63 (m, 1H, *minor*), 1.09 (d, *J* = 7.1 Hz, 6H, *major*), 1.07 (d, *J* = 6.9 Hz, 6H, *minor*); **$^{13}$C NMR** (101 MHz, $CD_3CN$) δ 165.68 (*minor*), 163.79 (*major)*, 61.57 *(major)*, 57.07 (*minor*), 31.04 (*minor)*, 26.70 (*major)*, 20.33 (*minor*), 19.03 (*major)*; **HRMS** (ESI+): m/z calc. for $C_5H_{12}NO_3$ $[M+H]^+$ 118.0863, found 118.0861.

*(E/Z)-1-cyclopropyl-3-hydroxypropan-2-one oxime* **(6c)**

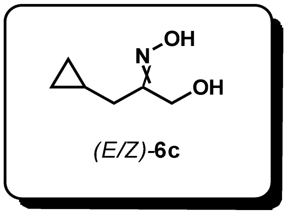


Following methods A2 and C, **6c** was prepared from commercial acid chloride (1.19 g, 10.00 mmol). *(E/Z)*-**6c**: Yellow solid (0.26 g, 20% overall yield); **R*f*** ($SiO_2$; *n*-pent:EtOAc 20:80) = 0.55, 0.6 (two C=N double bond isomers) (non-UV active, stained with $KMnO_4$); **IR** (neat, $cm^{-1}$): 3254, 2917, 1426, 1219, 1048, 1019, 960, 828; **$^1$H NMR** (400 MHz, $CDCl_3$) (mixture of two C=N double bond isomers in a 75:25 ratio) δ 4.45 (s, 2H, *minor*), 4.32 (s, 2H, *major*), 2.31 (d, J = 7.2 Hz, 2H, *major*), 2.20 (d, J = 6.9 Hz, 2H, *minor*), 0.99 – 0.83 (m, 2H, *major+minor*), 0.59 – 0.44 (m, 2H (*major*) + 2H (*minor*)), 0.25 – 0.09 (m, 2H (*major*) + 2H (*minor*)); **$^{13}$C NMR** (101 MHz, $CDCl_3$) δ 161.24 (*minor*), 159.89 (*major*), 62.72 (*major*), 58.67 (*minor*), 36.69 (*minor*), 30.27 (*major*), 7.72 (*major*), 6.90 (*minor*), 4.56 (*major*), 4.44 (*minor*); **HRMS** (ESI+): m/z calc. for $C_6H_{12}NO_2$ $[M+H]^+$ 130.08626 found 130.08637.

*(E/Z)-1-cyclobutyl-2-hydroxyethan-1-one oxime* **(6d)**

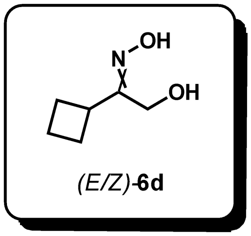


Following methods A1 and C, **6d** was prepared from acyl chloride (889 mg, 7.50 mmol). *(E/Z)*-**6d**: White solid (194 mg, 18% overall yield); **R*f*** ($SiO_2$; *n*-pent:EtOAc 20:70) = 0.4, 0.35 (two C=N double bond isomers) (non-UV active, stained with $KMnO_4$); **IR** (neat, $cm^{-1}$): $\nu_{max}$ = 2970, 1740, 1369, 1229, 1217, 527; **$^1$H NMR** (400 MHz, $CDCl_3$) (mixture of two C=N double bond isomers in a 55:45 ratio) δ 4.81 – 4.00 (br. s, 2H, *major+minor*), 4.35 (s, 2H, *major*), 4.34 (d, J = 1.3 Hz, 2H, *minor*), 3.68 (t, J = 9.2 Hz, 1H, *minor*), 3.24 (p, J = 8.5 Hz, 1H, *major*), 2.31 – 1.78 (m, 6H (*major*) + 6H (*minor*)); **$^{13}$C NMR** (101 MHz, CDCl3) δ 163.19 (*major*), 163.22 (*minor*), 61.50 (*minor*), 58.46 (*major*), 37.33 (*major*), 32.98 (*minor*), 26.34 (*minor*), 25.88 (*major*), 19.74 (*minor*), 18.60 (*major*); **HRMS** (ESI+): m/z calc. for $C_6H_{12}NO_2$ $[M+H]^+$ 130.08626, found 130.08611.

*(E/Z)-1-cyclohexyl-2-hydroxyethan-1-one oxime* **(6e)**

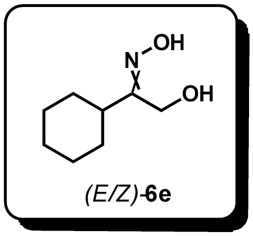


Prepared in two-steps from commercial cyclohexyl methyl ketone:

Step 1) Cyclohexyl methyl ketone (1.89 g, 15.0 mmol, 1 eq.) was weighed into a 250 mL round-bottom flask equipped with a magnetic stirring bar. A solution of trifluoroacetic acid (2.23 mL, 30.0 mmol, 2 eq.) in acetonitrile (75 mL) and water (15 mL) was added to the reaction vessel. Bis(trifluoroacetoxy)iodobenzene (12.9 g, 30.0 mmol, 2 eq.) was added in one portion. The flask was equipped with a reflux condenser, and the reaction mixture was heated to reflux (95 °C) for 3 h. The reaction mixture was cooled to room temperature and acetonitrile was removed under reduced pressure. The residue was diluted with brine (100 mL) and the aqueous phase was extracted in $Et_2O$ (3x 80 mL). The combined organic extracts were washed twice with a saturated aqueous solution of $NaHCO_3$ (2x 80 mL), dried over $Na_2SO_4$, filtered and concentrated under reduced pressure to a yellow liquid. The residue was purified by silica gel flash column chromatography (eluting with 80:20 to 50:50 *n*-pent:diethyl ether) to afford pure (cyclohexyl)-2-hydroxyethan-1-one **5e** as a light yellow liquid (1.70 g, 60% yield). The spectral data is in accordance with the literature.[2] **$^1$H NMR** (400 MHz, $CDCl_3$) δ 4.30 (s, 2H), 3.14 (br. s, 1H), 2.38 (tt, *J* = 11.6, 3.4 Hz, 1H), 1.88 – 1.76 (m, 4H), 1.72 – 1.64 (m, 1H), 1.47 – 1.36 (m, 2H), 1.35 – 1.22 (m, 3H).

Step 2) According to the general method C, oxime **6e** was prepared from pure (cyclohexyl)-2-hydroxyethan-1-one **5e** (1.00 g, 7.00 mmol). **6e**: Colourless solid (1.01 g, 92%); **$R_f$** ($SiO_2$; *n*-pent:EtOAc 60:40) = 0.45, 0.15 (two C=N double bond isomers) (non-UV active, stained with $KMnO_4$); **IR** (neat, $cm^{-1}$): $\nu_{max}$ = 3216.3, 2926.0, 2852.4, 1449.7, 948.8; **$^1$H NMR** (400 MHz, $CD_3OD$) (mixture of two C=N double bond isomers in a 58:42 ratio): δ 4.38 (s, 2H, *minor*), 4.10 (s, 2H, *major*), 3.05 (tt, *J* = 12.2, 3.2 Hz, 1H, *major*), 2.44 (ddq, *J* = 11.6, 8.6, 3.2 Hz, 1H, *minor*), 1.89 – 1.60 (m, 5H, *minor+major*), 1.57 – 1.43 (m, 1H, *minor+major*), 1.41 – 1.14 (m, 4H, *minor+major*); **$^{13}$C NMR** (101 MHz, $CD_3OD$) δ 165.25 (*minor*), 163.08 (*major*), 62.19 (*major*), 56.96 (*minor*), 41.00 (*minor*), 37.69 (*major*), 31.67 (*minor*), 29.83 (*major*), 27.49 (*minor*), 27.46 (*major*), 27.33 (*minor*), 27.23 (*major*); **HRMS** (ESI+): m/z calc. for $C_8H_{16}NO_2$ [M+H$^+$] 158.11756, found 158.11744.

*(E/Z)-1-hydroxy-5-methylhexan-2-one oxime* **(6f)**

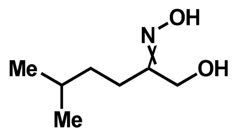


Prepared from commercial 4-methylvaleryl chloride (673 mg, 5.00 mmol) following the general methods A2 and C. *(E/Z)*-**6f**: colourless solid (420 mg, 58% overall yield); $\mathbf{R}_f$ ($SiO_2$; *n*-pent:EtOAc 60:40) = 0.35, 0.15 (two C=N double bond isomers) (non-UV active, stained with $KMnO_4$); **IR** (neat, $cm^{-1}$): $\nu_{max}$ = 3222.3, 3111.6, 2956.6, 1464.0, 989.1, 938.1; **$^1$H NMR** (400 MHz, $CDCl_3$) (mixture of two C=N double bond isomers in a 82:18 ratio): δ 5.50 (br. s, 2H, *minor+major*), 4.38 (s, 2H, *minor*), 4.19 (s, 2H, *major*), 2.38 – 2.31 (m, 2H, *major*), 2.31 – 2.25 (m, 2H, *minor*), 1.63 – 1.50 (m, 1H, *minor+major*), 1.47 – 1.34 (m, 2H, *minor+major*), 0.94 – 0.89 (m, 6H, *minor+major*); **$^{13}$C NMR** (101 MHz, $CDCl_3$) δ 162.06 (*minor*), 160.82 (*major*), 63.04 (*major*), 59.10 (*minor*), 35.20 (*minor*), 34.37 (*major*), 30.32 (*minor*), 28.50 (*major*), 27.94 (*minor*), 24.21 (*major*), 22.45 (*minor*), 22.40 (*major*); **HRMS** (ESI+): m/z calc. for $C_7H_{16}NO_2$ $[M+H]^+$ 146.11756, found 146.11756.

*(E/Z)-1-hydroxyhexan-2-one oxime* **(6g)**

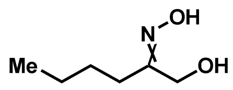


Prepared from commercial valeroyl chloride (603 mg, 5.00 mmol) following the general methods A2 and C. *(E/Z)*-**6g**: colourless oil (461 mg, 70% overall yield); $\mathbf{R}_f$ ($SiO_2$; *n*-pent:EtOAc 60:40) = 0.25, 0.15 (two C=N double bond isomers) (non-UV active, stained with $KMnO_4$); **IR** (neat, $cm^{-1}$): $\nu_{max}$ = 3224.0, 3110.6, 2961.3, 1465.1, 990.8, 938.2; **$^1$H NMR** (400 MHz, $CDCl_3$) (mixture of two C=N double bond isomers in a 70:30 ratio): δ 5.58 (br. s, 2H, *minor+major*), 4.38 (s, 2H, *minor*), 4.19 (s, 2H, *major*), 2.39 – 2.31 (m, 2H, *major*), 2.31 – 2.24 (m, 2H, *minor*), 1.57 – 1.45 (m, 2H, *minor+major*), 1.43 – 1.29 (m, 2H, *minor+major*), 0.97 – 0.87 (m, 3H, *minor+major*); **$^{13}$C NMR** (101 MHz, $CDCl_3$) δ 161.85 (*minor*), 160.66 (*major*), 63.08 (*major*), 59.06 (*minor*), 32.00 (*minor*), 28.39 (*minor*), 27.70 (*major*), 25.92 (*major*), 23.06 (*major*), 22.49 (*minor*), 13.90 (*major*), 13.89 (*minor*); **HRMS** (ESI+): m/z calc. for $C_6H_{14}NO_2$ $[M+H]^+$ 132.10191, found 132.10185.

*(E/Z)-5-chloro-1-hydroxypentan-2-one oxime* **(6h)**

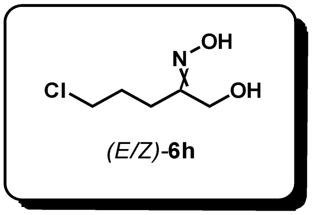


Following methods A2 and C, **6h** was prepared from commercial acid chloride (1.09 g, 7.72 mmol). The crude product was used in the next steps without further purification (attempted purification by silica gel column chromatography led to decomposition). **$^{1}$H NMR** (400 MHz, $CDCl_3$) (mixture of two C=N double bond isomers in a 70:30 ratio) δ 4.36 (d, *J* = 1.5 Hz, 2H *minor*), 4.15 (d, *J* = 2.1 Hz, 2H, *major*), 3.56 – 3.44 (m, 4H, *minor+major*), 2.49 – 2.37 (m, 4H, *minor+major*), 2.05 – 1.90 (m, 4H, *minor+major*).

*(E/Z)-2-hydroxy-1-phenylethan-1-one oxime* **(6i)**

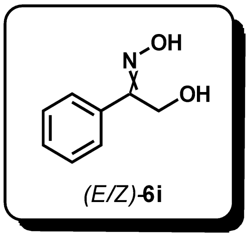


Prepared from commercial 2-hydroxyacetophenone (681 mg, 5.00 mmol) following the general method C. *(E/Z)*-**6i**: White solid (687 mg, 91%); **R*f*** ($SiO_2$; *n*-pent:EtOAc 60:40) = 0.45, 0.25 (two C=N double bond isomers); **IR** (neat, $cm^{-1}$): $\nu_{max}$ = 3282.5, 2904.5, 1637.6, 1496.7, 1442.8, 1300.6, 1042.3, 1018.9, 983.2, 916.6, 759.9, 695.4; **$^{1}$H NMR** (400 MHz, $CDCl_3$) (mixture of two C=N double bond isomers in a 75:25 ratio) δ 7.64 – 7.36 (m, 5H *(minor)* + 5H *(major)*), 4.77 (s, 2H, *major*), 4.54 (s, 2H, *minor*) **$^{13}$C NMR** (101 MHz, $CDCl_3$) δ 159.68, 134.34, 130.22, 130.19, 129.18, 128.95, 128.42, 127.20, 64.33 (*minor*), 58.17 (*major*). The mass spectroscopic data was in accordance the one reported in the literature.[3]

*(E/Z)-1-(4-fluorophenyl)-2-hydroxyethan-1-one oxime* **(6j)**

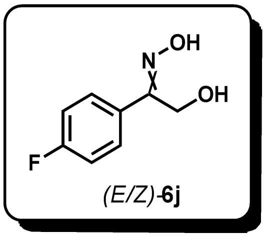


Prepared from commercial 4'-fluoro-2-hydroxyacetophenone (771 mg, 5.00 mmol) following the general method C.

***Minor isomer:*** Colourless solid (157 mg, 19%); **R*f*** ($SiO_2$; *n*-pent:EtOAc 60:40) = 0.35; **m.p.** = 83-84 °C; **IR** (neat, $cm^{-1}$): $\nu_{max}$ = 3249.8, 1603., 1514.9, 1236.0, 839.2; **$^1$H NMR** (400 MHz, Acetone-$d_6$) (contains 5% of *Z*-isomer) δ 10.49 (s, 1H), 7.83 – 7.67 (m, 2H), 7.23 – 7.05 (m, 2H), 4.78 (d, J = 5.0 Hz, 2H), 4.11 (t, J = 6.1 Hz, 1H); **$^{19}$F NMR** (377 MHz, Acetone-$d_6$) δ -115.04; **$^{13}$C NMR** (101 MHz, Acetone-$d_6$) δ 163.86 (d, *J* = 245.7 Hz), 157.45, 132.70 (d, *J* = 3.4 Hz), 129.87 (d, *J* = 8.3 Hz), 115.61 (d, *J* = 21.7 Hz), 54.97; **HRMS** (ESI+): m/z calc. for $C_8H_9NO_2F$ $[M+H]^+$ 170.06118, found 170.06122.

***Major isomer:*** Colourless solid (620 mg, 73%); **R*f*** ($SiO_2$; *n*-pent:EtOAc 60:40) = 0.10; **m.p.** = 152-153 °C; **IR** (neat, $cm^{-1}$): $\nu_{max}$ = 3208.7, 1607.4, 1515.4, 1244.9, 989.7; **$^1$H NMR** (400 MHz, Acetone-$d_6$) δ 10.17 (s, 1H), 7.85 – 7.69 (m, 2H), 7.25 – 7.10 (m, 2H), 4.42 (d, *J* = 5.9 Hz, 2H), 4.07 (t, *J* = 5.9 Hz, 1H); **$^{19}$F NMR** (377 MHz, Acetone-$d_6$) δ -113.99; **$^{13}$C NMR** (101 MHz, Acetone-$d_6$) δ 163.37 (d, *J* = 246.5 Hz), 154.92, 132.09 (d, *J* = 8.3 Hz), 129.68 (d, *J* = 3.5 Hz), 115.43 (d, *J* = 21.5 Hz), 64.57; **HRMS** (ESI+): m/z calc. for $C_8H_9NO_2F$ $[M+H]^+$ 170.06118, found 170.06119.

*(E/Z)-2-hydroxy-1-(4-(trifluoromethyl)phenyl)ethan-1-one oxime* **(6k)**

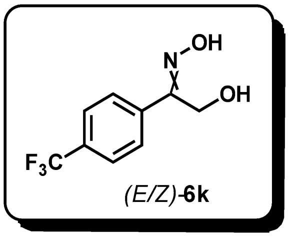


Prepared from commercial 4′-(trifluoromethyl)acetophenone (1.00 g, 5.30 mmol) following the general methods B and C. **6k**: White solid (423 mg, 36% overall yield); **R*f*** ($SiO_2$; *n*-pent:EtOAc 70:30) = 0.65, 0.45 (two C=N double bond isomers); **IR** (neat, $cm^{-1}$): $\nu_{max}$ = 3281.9, 1407.1, 1328.0, 1168.3, 1123.5, 1065.2, 846.5; **$^1$H NMR** (400 MHz, $CDCl_3$) (mixture of two C=N double bond isomers in a 75:25 ratio) δ 7.66 – 7.51 (m, 4H (*minor*) + 4H (*major*)), 4.68 (s, 2H, *major*), 4.43 (s, 2H, *minor*); **$^{13}$C NMR** (101 MHz, $CDCl_3$) δ 157.99 (*major*), 155.26 (*minor*), 137.25 (*major*), 134.26 (*minor*), 131.64 (q, J = 32.5 Hz) (*major*+*minor*), 128.64 (*minor*), 127.21 (*major*), 125.65 (q, J = 3.7 Hz) (*major*), 125.42 (q, J = 3.6 Hz) (*minor*), 123.90 (q, J = 272.3 Hz) (*major*), 123.77 (q, J = 272.3 Hz) (*minor*), 63.45 (*minor*), 56.75 (*major*); **$^{19}$F NMR** (377 MHz, $CDCl_3$) δ -62.90, -63.06.

*(E/Z)-1-(3,4-dichlorophenyl)-2-hydroxyethan-1-one oxime* **(6l)**

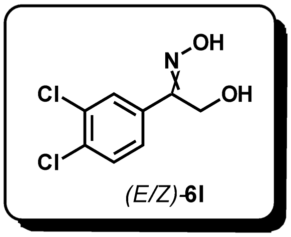

Prepared from commercial 3,4-dichlorobenzoyl chloride (1.00 g, 4.77 mmol) following the general methods A1 and C. **6l**: Yellow solid (501 mg, 47% overall yield); $\mathbf{R}_f$ ($SiO_2$; *n*-pent:EtOAc 60:40) = 0.35, 0.2 (two C=N double bond isomers); **IR** (neat, $cm^{-1}$): $\nu_{max}$ = 3252.6, 2785.0, 1695.0, 1473.5, 1381.5, 1138.4, 1030.4, 820.6, 763.5; **$^1$H NMR** (400 MHz, $CD_3OD$) (mixture of two C=N double bond isomers in a 75:25 ratio) δ 7.85 – 7.84 (m, 2H, *minor+major*), 7.62 – 7.50 (m, 2H (*minor*) + 2H (*major*)), 4.73 (s, 2H, *major*), 4.40 (s, 2H, *minor*); **$^{13}$C NMR** (101 MHz, $CD_3OD$) δ 156.54, 155.73, 136.96, 133.70, 133.41, 133.15, 132.10, 131.27, 131.16, 129.87, 129.78, 127.77, 64.35 (*minor*), 54.64 (*major*); **HRMS** (ESI+): m/z calc. for $C_8H_8NO_2Cl_2$ $[M+H]^+$ 219.99266, found 219.99252.

*(E/Z)-2-hydroxy-1-(4-methoxyphenyl)ethan-1-one oxime* **(6m)**

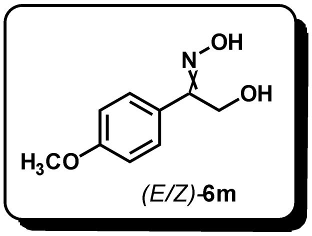

Prepared from commercial 4'-methoxyacetophenone (1.50 g, 10.00 mmol) following the general methods B and C. **6m**: White solid (400 mg, 22% overall yield); $\mathbf{R}_f$ ($SiO_2$; *n*-pent:EtOAc 70:30) = 0.45, 0.25 (two C=N double bond isomers); **IR** (neat, $cm^{-1}$): $\nu_{max}$ = 3288.2, 2924.3, 1606.5, 1515.2, 1301.1, 1252.5, 1180.5, 978.9, 917.7; **$^1$H NMR** (400 MHz, $CD_3OD$) (mixture of two C=N double bond isomers in a 80:20 ratio) δ 7.71 – 7.64 (m, 2H, *minor*), 7.63 – 7.55 (m, 2H, *major*), 6.96 – 6.93 (m, 2H, *minor*), 6.92 – 6.89 (m, 2H, *major*), 4.71 (s, 2H, *major*), 4.40 (s, 2H, *minor*), 3.81 (s, 3H, *minor*), 3.80 (s, 3H, *major*); **$^{13}$C NMR** (101 MHz, $CD_3OD$) δ 161.34 (*major*), 161.02 (*minor*), 158.15 (*major*), 155.43 (*minor*), 131.22, 128.94, 128.25, 124.94, 114.19, 113.88, 64.08, 55.28, 55.25, 54.95; **HRMS** (ESI+): m/z calc. for $C_9H_{12}NO_3$ $[M+H]^+$ 182.08117, found 182.08139.

*1-(furan-2-yl)-2-hydroxyethan-1-one oxime* **(6n)**

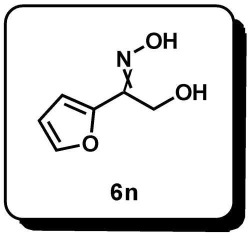

Prepared from commercial 2-furoyl chloride (979 mg, 7.50 mmol) following the general methods A1 and C. **6n**: Brown solid (430 mg, 41% overall yield); $\mathbf{R}_f$ ($SiO_2$; *n*-pent:EtOAc 30:70) = 0.3; **IR** (neat, $cm^{-1}$): $\nu_{max}$ = 3148.4, 2923.7, 2854.3, 1725.0, 1638.0, 1565.8, 1460.1, 1395.5, 1269.1, 1117.1, 1082.2, 1018.6, 979.9, 937.2, 884.2, 835.2, 759.7, 592.8; **$^1$H NMR** (400 MHz, $CD_3OD$) (single C=N double bond isomer) δ 6.03 (d, J = 2.1 Hz, 1H), 5.85 (d, J = 3.5 Hz, 1H), 5.03 (dd, J = 3.5, 1.8 Hz, 1H), 2.97 (s, 2H). **$^{13}$C NMR** (101 MHz, $CD_3OD$) δ 146.45, 146.38, 143.47, 118.59, 112.57, 61.70; **HRMS** (ESI+): m/z calc. for $C_6H_8NO_3$ $[M+H]^+$ 142.04987, found 142.05000.

*1-hydroxy-3-(thiophen-2-yl)propan-2-one oxime* **(6o)**

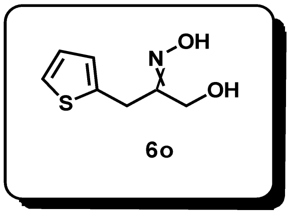

Prepared from commercial 2-thienylacetyl chloride (0.68 mL, 5.50 mmol) following the general methods A2 and C. **6o:** off-white solid (584 mg, 62% overall yield); $\mathbf{R}_f$ ($SiO_2$; *n*-pent:EtOAc 50:50) = 0.3; **m.p.** = 95-96 °C; **IR** (neat, $cm^{-1}$): $\nu_{max}$ = 3195.7, 3111.4, 2920.1, 1430.8, 993.8, 956.5, 686.7; **$^1$H NMR** (400 MHz, acetone-$d_6$) (single C=N double bond isomer) δ 10.06 (s, 1H), 7.24 (dd, *J* = 5.0, 1.5 Hz, 1H), 6.96 – 6.89 (m, 2H), 4.09 – 4.00 (m, 3H), 3.96 (d, *J* = 0.8 Hz, 2H); **$^{13}$C NMR** (101 MHz, acetone-$d_6$) δ 157.44, 139.48, 127.43, 127.07, 124.96, 62.42, 25.22; **HRMS** (ESI+): m/z calc. for $C_7H_{10}NO_2S$ $[M+H]^+$ 172.04268, found 172.04259.

*(Z)-2-hydroxy-1-(1-hydroxy-4-methylpentan-2-yl)diazene 1-oxide* **(7a = leudiazen)**

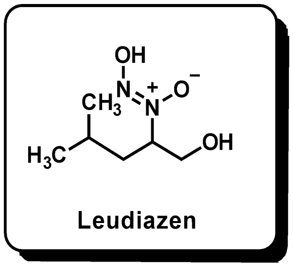


Prepared from the corresponding oxime precursor **6a** (1.38 g, 10.50 mmol) following the general methods D and F. **Leudiazen**: off-white solid (1.36 g, 80%). The spectral data matched the described in the literature.[5]

*(Z)-2-hydroxy-1-(1-hydroxy-3-methylbutan-2-yl)diazene 1-oxide* **(7b = valdiazen)**

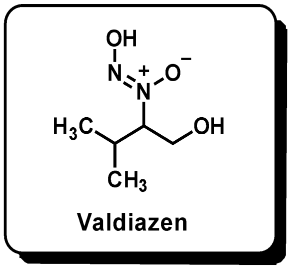


Prepared from the corresponding oxime precursor **6b** (269 mg, 2.30 mmol) following the general methods D and F. **Valdiazen**: off-white solid (236 mg, 69%). The spectral data matched the described in the literature.[6]

*(Z)-1-(1-cyclopropyl-3-hydroxypropan-2-yl)-2-hydroxydiazene 1-oxide* **(7c)**

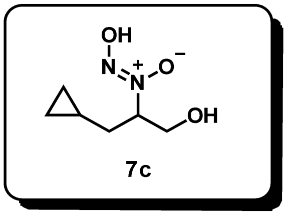


Prepared from the corresponding oxime precursor **6c** (110 mg, 0.85 mmol) following the general methods D and F. **7c**: off-white solid (86 mg, 63%); **m.p.** = 63-64 °C; **IR** (neat, $cm^{-1}$): $\nu_{max}$ = 3080.9, 2934.1, 1455.6, 1335.5, 1047.0, 939.0; **$^1$H NMR** (400 MHz, $CD_3OD$) δ 4.46 – 4.36 (m, 1H), 3.92 (dd, $J$ = 11.8, 9.4 Hz, 1H), 3.68 (dd, $J$ = 11.8, 3.9 Hz, 1H), 1.73 (ddd, $J$ = 14.4, 9.9, 7.1 Hz, 1H), 1.53 (ddd, $J$ = 14.4, 7.0, 4.3 Hz, 1H), 0.64 (dddt, $J$ = 15.0, 8.1, 7.1, 4.8 Hz, 1H), 0.51 – 0.36 (m, 2H), 0.13 – 0.03 (m, 2H); **$^{13}$C NMR** (101 MHz, $CD_3OD$) δ 77.06, 62.59, 34.20, 8.31, 4.56, 4.13; **HRMS** (ESI-): m/z calc. for $C_6H_{11}N_2O_3$ [M-$H^+$] 159.07752, found 159.07742.

*(Z)-1-(1-cyclobutyl-2-hydroxyethyl)-2-hydroxydiazene 1-oxide* **(7d)**

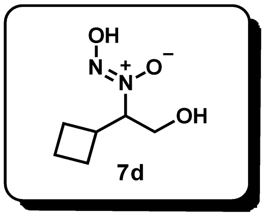


Prepared from the corresponding oxime precursor **6d** (136 mg, 1.05 mmol) following the general methods D and F. **7d**: colourless solid (126 mg, 75%); **m.p.** = 67-68 °C; **IR** (neat, $cm^{-1}$): $\nu_{max}$ = 3368.2, 2949.9, 1453.3, 1057.2; **$^1$H NMR** (400 MHz, $CD_3OD$) δ 4.27 (ddd, *J* = 10.3, 9.3, 3.2 Hz, 1H), 3.88 (dd, *J* = 12.0, 9.3 Hz, 1H), 3.63 (dd, *J* = 12.0, 3.2 Hz, 1H), 2.83 – 2.68 (m, 1H), 2.11 (dddd, *J* = 13.8, 5.8, 4.0, 2.4 Hz, 1H), 2.03 – 1.78 (m, 5H); **$^{13}$C NMR** (101 MHz, $CD_3OD$) δ 81.30, 60.35, 35.39, 26.19, 26.02, 18.79; **HRMS** (ESI-): m/z calc. for $C_6H_{11}N_2O_3$ [M-$H^+$] 159.07752, found 159.07752.

*(Z)-1-(1-cyclohexyl-2-hydroxyethyl)-2-hydroxydiazene 1-oxide* **(7e)**

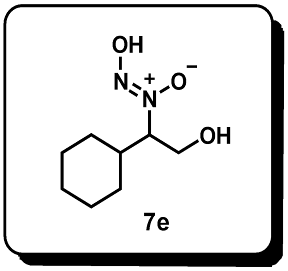


Prepared from the corresponding oxime precursor **6e** (472 mg, 3.00 mmol) according to a modified version of the general method E [crude hydroxylamine **3e** was purified by trituration using 1:1 *n*-pent:$Et_2O$ (2x 10 mL), discarding the liquid], followed by the general method F.
**3e**: Off-white solid (440 mg, 92%); **$R_f$** ($SiO_2$; EtOAc:MeOH 95:5) = 0.35 (non-UV active, stained with $KMnO_4$); **m.p.** = 74-75 °C; **IR** (neat, $cm^{-1}$): $\nu_{max}$ = 3271.0, 2921.6, 2850.7, 1448.1, 1039.8; **$^1$H NMR** (400 MHz, $CD_3OD$) δ 3.69 (dd, *J* = 11.1, 4.0 Hz, 1H), 3.54 (dd, *J* = 11.2, 7.6 Hz, 1H), 2.59 (ddd, *J* = 7.6, 6.3, 4.0 Hz, 1H), 1.89 – 1.53 (m, 6H), 1.35 – 0.96 (m, 5H); **$^{13}$C NMR** (101 MHz, $CD_3OD$) δ 69.24, 60.51, 38.17, 30.98, 30.33, 27.71, 27.67, 27.65; **HRMS** (ESI+): m/z calc. for $C_8H_{18}NO_2$ [M+$H^+$] 160.13321, found 160.13305.

**7e**: colourless solid (71 mg, 86%; starting from 70 mg of **3e**); **m.p.** = 116-117 °C; **IR** (neat, $cm^{-1}$): $\nu_{max}$ = 3321.7, 2929.8, 2854.3, 1450.7, 1056.5, 935.0; **$^1$H NMR** (400 MHz, $CD_3OD$) δ 4.06 (td, *J* = 9.1, 3.1 Hz, 1H), 3.98 (dd, *J* = 11.5, 9.4 Hz, 1H), 3.83 (dd, *J* = 11.5, 3.1 Hz, 1H), 1.90 – 1.60 (m, 5H), 1.50 (dtd, *J* = 13.3, 3.9, 2.2 Hz, 1H), 1.38 – 1.16 (m, 3H), 1.16 – 0.98 (m, 2H); **$^{13}$C NMR** (101 MHz, $CD_3OD$) δ 80.61, 61.00, 38.24, 30.65, 30.31, 27.14, 26.86, 26.70; **HRMS** (ESI-): m/z calc. for $C_8H_{15}N_2O_3$ [M-$H^+$] 187.10882, found 187.10873.

*(Z)-2-hydroxy-1-(1-hydroxy-5-methylhexan-2-yl)diazene 1-oxide* **(7f)**

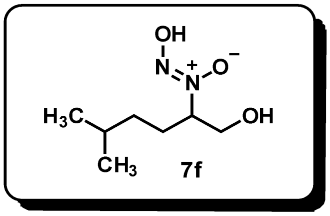


Prepared from the corresponding oxime precursor **6f** (145 mg, 1.00 mmol) following the general methods D and F. **7f**: colourless solid (109 mg, 62%); **m.p.** = 42-43 °C; **IR** (neat, $cm^{-1}$): $\nu_{max}$ = 3366.7, 2957.1, 2872.4, 1457.6, 1062.6, 1037.8, 937.6; **$^{1}$H NMR** (500 MHz, $CD_3OD$) δ 4.27 (dddd, *J* = 10.2, 8.9, 4.4, 2.4 Hz, 1H), 3.96 – 3.86 (m, 1H), 3.67 (ddd, *J* = 11.8, 3.9, 1.1 Hz, 1H), 1.87 – 1.76 (m, 1H), 1.67 – 1.49 (m, 2H), 1.20 (ddt, *J* = 13.2, 10.6, 6.0 Hz, 1H), 1.13 – 1.03 (m, 1H), 0.93 – 0.86 (m, 6H); **$^{13}$C NMR** (126 MHz, $CD_3OD$) δ 76.99, 62.91, 35.70, 28.79, 27.17, 22.91, 22.53; **HRMS** (ESI-): m/z calc. for $C_7H_{15}N_2O_3$ [M-$H^+$] 175.10882, found 175.10884.

*(Z)-2-hydroxy-1-(1-hydroxyhexan-2-yl)diazene 1-oxide* **(7g)**

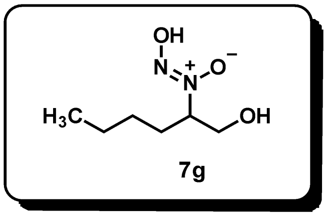


Prepared from the corresponding oxime precursor **6g** (98 mg, 0.75 mmol) following the general methods D and F. **7g**: off-white solid (73 mg, 60%); **m.p.** = 82-83 °C; **IR** (neat, $cm^{-1}$): $\nu_{max}$ = 3364.8, 2958.9, 2932.6, 1457.0, 1336.9, 1059.4, 937.6; **$^{1}$H NMR** (400 MHz, $CD_3OD$) δ 4.30 (ddt, *J* = 10.4, 9.3, 4.0 Hz, 1H), 3.91 (dd, *J* = 11.8, 9.3 Hz, 1H), 3.66 (dd, *J* = 11.8, 3.8 Hz, 1H), 1.89 – 1.76 (m, 1H), 1.60 (dddd, *J* = 13.9, 9.8, 5.9, 4.1 Hz, 1H), 1.46 – 1.14 (m, 4H), 0.91 (t, *J* = 7.1 Hz, 3H); **$^{13}$C NMR** (101 MHz, $CD_3OD$) δ 76.67, 62.88, 28.92, 28.77, 23.13, 14.12; **HRMS** (ESI-): m/z calc. for $C_6H_{13}N_2O_3$ [M-$H^+$] 161.09317, found 161.09329.

*(Z)-1-(5-chloro-1-hydroxypentan-2-yl)-2-hydroxydiazene 1-oxide* **(7h)**

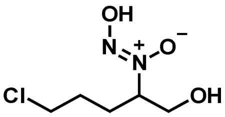


**7h**

Prepared from the corresponding crude oxime precursor **6h** (7.72 mmol theoretical) following the general methods E and G. **7h**: yellow solid (423 mg, 30% overall); **m.p.** = 101-104 °C; **IR** (neat, $cm^{-1}$): $\nu_{max}$ = 3368.8, 2937.5, 1450.0, 1335.8, 1279.6, 1050.7, 932.8, 651.0; **$^1$H NMR** (400 MHz, $CD_3OD$) δ 4.41 – 4.30 (m, 1H), 3.93 (dd, J = 11.8, 9.2 Hz, 1H), 3.68 (dd, J = 11.8, 3.8 Hz, 1H), 3.57 (t, J = 6.3 Hz, 2H), 2.03 – 1.89 (m, 1H), 1.86 – 1.62 (m, 3H); **$^{13}$C NMR** (101 MHz, $CD_3OD$) δ 76.03, 62.76, 44.80, 29.74, 26.70; **HRMS** (ESI+): m/z calc. for $C_5H_{10}N_2O_3Cl$ [M+H]+181.03854 , found 181.03848.

*(Z)-2-hydroxy-1-(2-hydroxy-1-phenylethyl)diazene 1-oxide* **(7i)**

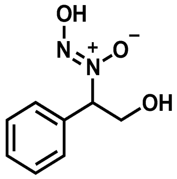


**7i**

Prepared from the corresponding oxime precursor **6i** (529 mg, 3.50 mmol) according to the general methods E and F.

**3i**: colourless solid (310 mg, 58%); **R*f*** ($SiO_2$; EtOAc:MeOH 95:5) = 0.5 (non-UV active, stained with $KMnO_4$); The spectral data matched the one described in the literature.[7]

**7i**: colourless solid (340 mg, 98%; starting from 291 mg of **3i**); **m.p.** = 106-107 °C; **IR** (neat, $cm^{-1}$): $\nu_{max}$ = 3221.4, 3064.1, 2940.6, 1454.4, 1064.2, 702.3; **$^1$H NMR** (400 MHz, $CD_3OD$) δ 7.48 (dd, *J* = 7.0, 2.6 Hz, 2H), 7.41 – 7.30 (m, 3H), 5.38 (dd, *J* = 9.7, 4.4 Hz, 1H), 4.43 (dd, *J* = 11.8, 9.7 Hz, 1H), 3.86 (dd, *J* = 11.8, 4.4 Hz, 1H); **$^{13}$C NMR** (101 MHz, $CD_3OD$) δ 136.03, 129.83, 129.62, 128.80, 77.48, 63.21; **HRMS** (ESI-): m/z calc. for $C_8H_9N_2O_3$ $[M-H]^-$ 181.06187, found 181.06188.

*(Z)-1-(1-(4-fluorophenyl)-2-hydroxyethyl)-2-hydroxydiazene 1-oxide* **(7j)**

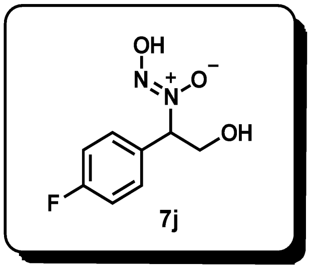

Prepared from the corresponding oxime precursor **6j** (5.24 g, 31.00 mmol) according to a modified version of the general method E (crude hydroxylamine **3j** was purified by trituration in $Et_2O$, discarding the liquid), followed by the general method G (product **7j** was purified by filtration through a short silica plug, eluting in pure EtOAc).

**3j**: off-white solid (5.00 g, 94%); $\mathbf{R}_f$ ($SiO_2$; EtOAc:MeOH 95:5) = 0.45 (poorly-UV active, stained with $KMnO_4$); **m.p.** = 126-127 °C; **IR** (neat, $cm^{-1}$): $\nu_{max}$ = 3254.3, 1509.3, 1225.2, 1024.9, 838.6; **$^1$H NMR** (400 MHz, $CD_3OD$) δ 7.47 – 7.32 (m, 2H), 7.10 – 6.98 (m, 2H), 4.05 (t, *J* = 6.5 Hz, 1H), 3.67 (d, *J* = 6.5 Hz, 2H); **$^{13}$C NMR** (101 MHz, $CD_3OD$) δ 163.68 (d, *J* = 243.6 Hz), 136.80 (d, *J* = 3.2 Hz), 130.82 (d, *J* = 8.1 Hz), 115.90 (d, *J* = 21.4 Hz), 68.98, 64.63; **$^{19}$F NMR** (376 MHz, $CD_3OD$) δ -117.63; **HRMS** (ESI+): m/z calc. for $C_8H_{11}FNO_2$ $[M+H]^+$ 172.07683, found 172.07705.

**7j**: yellow oil (5.26 g, 99%; starting from 4.50 g of **3j**); **IR** (neat, $cm^{-1}$): $\nu_{max}$ = 3351.7, 3076.8, 2941.0, 1511.8, 1453.9, 1231.0, 1061.7, 837.9; **$^1$H NMR** (400 MHz, $CD_3OD$) δ 7.61 – 7.51 (m, 2H), 7.19 – 7.10 (m, 2H), 5.45 (dd, *J* = 10.1, 4.1 Hz, 1H), 4.45 (dd, *J* = 11.8, 10.1 Hz, 1H), 3.84 (dd, *J* = 11.8, 4.1 Hz, 1H); **$^{13}$C NMR** (126 MHz, $CD_3OD$) δ 164.69 (d, *J* = 247.1 Hz), 131.18 (d, *J* = 8.6 Hz), 130.71 (d, *J* = 3.4 Hz), 116.56 (d, *J* = 21.8 Hz), 78.53, 62.87; **$^{19}$F NMR** (377 MHz, $CD_3OD$) δ -114.35; **HRMS** (ESI-): m/z calc. for $C_8H_8FN_2O_3$ $[M-H]^-$ 199.05244, found 199.05245.

*(S,Z)-1-(1-(4-fluorophenyl)-2-hydroxyethyl)-2-hydroxydiazene 1-oxide* (***(S)*-7j**)

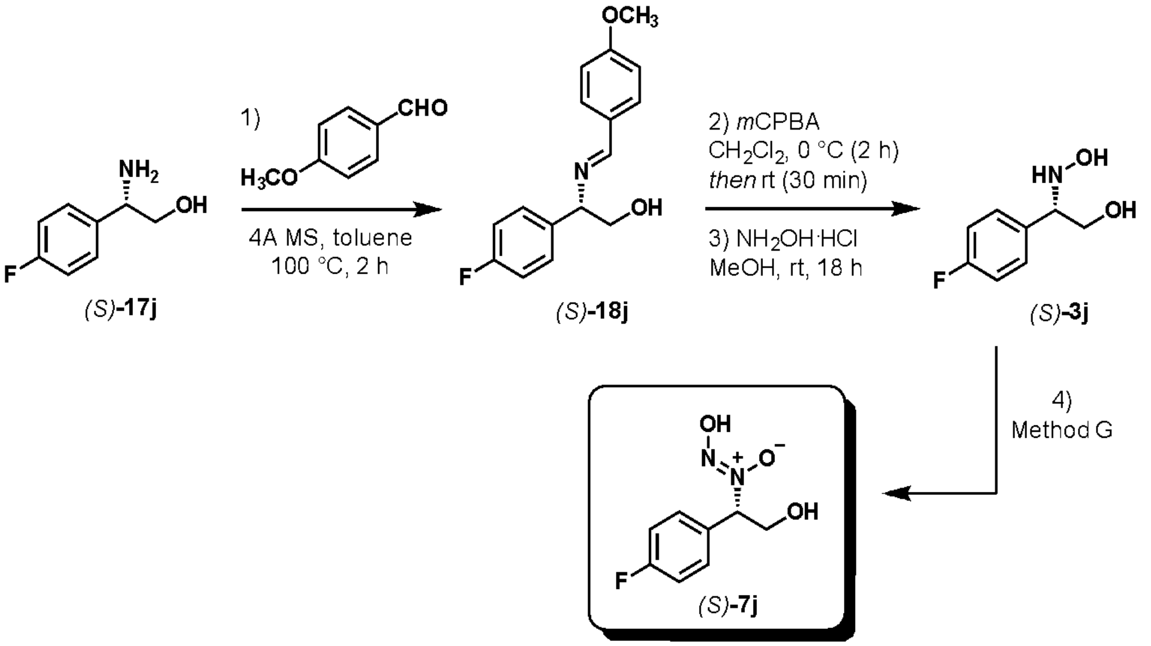


Prepared from commercially available (*S*)-2-amino-2-(4-fluorophenyl)ethanol **17j** *via* a modified reported protocol to prepare chiral *N*-hydroxylamines (steps 1-3),[8] followed by the general method G (step 4):

Step 1) A flame-dried round-bottom flask, equipped with a magnetic stirring bar and 4Å molecular sieves, was charged with (*S*)-2-amino-2-(4-fluorophenyl)ethanol **17j** (492 mg, 3.17 mmol, 1 eq.) and dry toluene (6 mL) at room temperature under a nitrogen atmosphere. *p*-Anisaldehyde (0.39 mL, 3.17 mmol, 1 eq.) was added, a reflux condenser was attached to the flask and the reaction was heated to 100 °C stirring for 2 h. The reaction mixture was cooled to room temperature and filtered through a short plug of Celite, washing with additional toluene. The filtrate was concentrated under reduced pressure to afford the crude imine product *(S)*-**18j** (866 mg) which was used in the next step without further purification.

Step 2) A solution of the crude imine in dry $CH_2Cl_2$ (4 mL) was cooled to 0 °C. A solution of *m*-chloroperbenzoic acid (*m*-CPBA, 710 mg, 3.17 mmol, 1 eq., ≤77% purity) in dry $CH_2Cl_2$ (8 mL) was added dropwise causing the formation of a white precipitate. The suspension was stirred at 0 °C for 2 h then 30 min at room temperature. The white solid (*m*-CPBA) was filtered off, and the filtrate was washed with an aqueous saturated $NaHCO_3$ solution. The organic phase was dried over $MgSO_4$, filtered, and concentrated under reduced pressure to yield a yellow oil.

Step 3) The oil residue was dissolved in dry MeOH (7 mL) and hydroxylamine hydrochloride (286 mg, 4.12 mmol, 1.3 eq) was added. After stirring at room temperature for 18 h, the reaction mixture was concentrated under reduced pressure. 1 N aqueous HCl was added (ensure pH = 1) and the aqueous layer was washed multiple times with $Et_2O$ (e.g. 3 times, until no non-polar materials were observed by TLC), basified to a pH >8 by adding excess of $Na_2CO_3$ and extracted three times in EtOAc. The combined organic layers were dried over $Na_2SO_4$, filtered, and concentrated under reduced pressure yielding the analytically pure product *(S)*-**3j** (510 mg, 94% over 3 steps). The experimental data matched the one of *rac*-**3j** described above.

Step 4) *(S)*-**7j** was prepared from *(S)*-**3j** (200 mg, 1.17 mmol) following the general method G. *(S)*-**7j**: yellow oil (203 mg, 85%); $[\alpha]_D^{20}$= +17.75 (*c* = 1.0, MeOH); the rest of the experimental data matched the one of *rac*-**7j** described above.

*(R,Z)-1-(1-(4-fluorophenyl)-2-hydroxyethyl)-2-hydroxydiazene 1-oxide* (***(R)*-7j**)

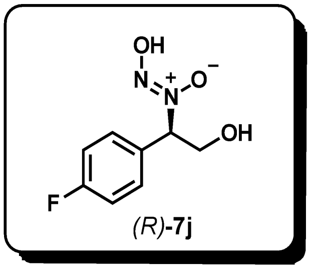

*(R)*-**7j** was prepared as described above for *(S)*-**7j**. *(R)*-**7j**: Yellow oil (278 mg, 30%); $[\alpha]_D^{20}$= -11.52 (*c* = 1.0, MeOH); the rest of the experimental data matched the one of *rac*-**7j** described above.

*(Z)-2-hydroxy-1-(2-hydroxy-1-(4-(trifluoromethyl)phenyl)ethyl)diazene 1-oxide* (**7k**)

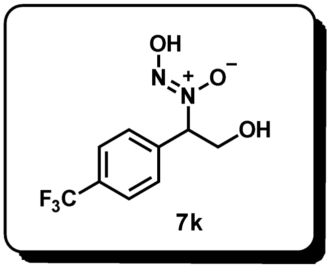

Prepared from the corresponding oxime precursor **6k** (120 mg, 0.55 mmol) according to a modified version of the general method E (crude hydroxylamine **3k** was purified by trituration in $Et_2O$, discarding the liquid), followed by the general method G (product **7k** was purified by trituration in *n*-pent/$Et_2O$ 60:40, discarding the liquid).

**3k**: off-white solid (121 mg, 99%); $\mathbf{R_f}$ ($SiO_2$; EtOAc:MeOH 95:5) = 0.4 (poorly-UV active, stained with $KMnO_4$); **m.p.** = 115-119 °C; **IR** (neat, $cm^{-1}$): $\nu_{max}$ = 3250.6, 2886.0, 1621.3, 1418.2, 1327.0, 1165.4, 1125.4, 1066.9, 1017.2, 961.7, 880.0, 841.5, 763.9, 607.6, 544.3; **$^1$H NMR** (400 MHz, $CD_3OD$) δ 7.64 – 7.58 (m, 4H), 4.14 (dd, J = 7.8, 5.2 Hz, 1H), 3.75 – 3.62 (m, 2H); **$^{13}$C NMR** (101 MHz, $CD_3OD$) δ 157.99, 155.26, δ 139.31, 132.78 (q, J = 32.3 Hz), 130.13, 127.10 (q, J = 4.0 Hz), 125.82 (q, J = 271.5 Hz), 79.03, 63.23; **$^{19}$F NMR** (377 MHz, $CD_3OD$) δ -63.98; **HRMS** (ESI+): m/z calc. for $C_9H_{11}F_3NO_2$ $[M+H]^+$ 222.0736, found 222.0739.

**7k**: white solid (96 mg, 70%; starting from 120 mg of **3k**); **m.p.** = 73-76 °C; **IR** (neat, $cm^{-1}$): $\nu_{max}$ = 3352.3, 1422.6, 1326.9, 1168.4, 1122.7, 1068.1, 1020.2, 842.3, 698.1; **$^1$H NMR** (400 MHz, $CD_3OD$) δ 7.73 – 7.68 (m, 4H), 5.53 (dd, J = 9.9, 4.2 Hz, 1H), 4.45 (dd, J = 11.8, 9.8 Hz, 1H), 3.89

(dd, J = 11.8, 4.2 Hz, 1H); **$^{13}$C NMR** (101 MHz, $CD_3OD$) δ 139.31, 132.78 (q, J = 32.3 Hz), 130.13, 127.10 (q, J = 4.0 Hz), 125.82 (q, J = 271.5 Hz), 79.03, 63.23; **$^{19}$F NMR** (377 MHz, $CD_3OD$) δ -64.32; **HRMS** (ESI-): m/z calc. for $C_9H_8F_3N_2O_3$ $[M-H]^-$ 249.04925, found 249.04925.

*(Z)-1-(1-(3,4-dichlorophenyl)-2-hydroxyethyl)-2-hydroxydiazene 1-oxide* **(7l)**

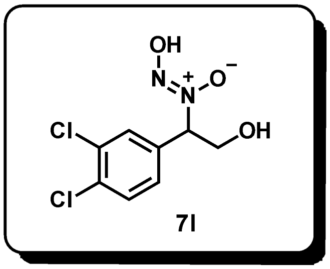


Prepared from the corresponding oxime precursor **6l** (400 mg, 1.81 mmol) according to a modified version of the general method E (crude hydroxylamine **3l** was purified by trituration in $Et_2O$, discarding the liquid), followed by the general method G (product **7l** was purified by trituration in $Et_2O$, discarding the liquid).

**3l**: off-white solid (320 mg, 80%); **$R_f$** ($SiO_2$; EtOAc:MeOH 95:5) = 0.5 (poorly-UV active, stained with $KMnO_4$); **m.p.** = 73-75 °C; **IR** (neat, $cm^{-1}$): $\nu_{max}$ = 3286.8, 2927.4, 1562.4, 1469.9, 1396.4, 1319.94, 1197.1, 1134, 1030.5, 882.5, 822.3, 733, 707.2, 675, 601, 528.2, 467.8; **$^1$H NMR** (400 MHz, $CD_3OD$) δ 7.77 (d, J = 2.2 Hz, 1H), 7.63 (d, J = 8.4 Hz, 1H), 7.49 (dd, J = 8.4, 2.1 Hz, 1H), 4.54 (dd, J = 6.4, 4.3 Hz, 1H), 4.03 (qd, J = 11.9, 5.4 Hz, 2H). **$^{13}$C NMR** (101 MHz, $CD_3OD$) δ 135.06, 134.04 (d, J = 2.7 Hz), 132.57, 132.30, 130.34, 66.99, 60.47; **HRMS** (ESI+): m/z calc. for $C_9H_{12}NO_2Cl_2$ $[M+H]^+$ 236.02396, found 236.02359.

**7l**: yellow solid (310 mg, 91%; starting from 320 mg of **3l**); **m.p.** = 92-93 °C: **IR** (neat, $cm^{-1}$): $\nu_{max}$ = 3324.8, 2973.6, 2886, 1380.11, 1087.3, 1045.34, 879.7, 618.9; **$^1$H NMR** (400 MHz, $CD_3OD$) δ 7.60 – 7.51 (m, 2H), 7.18 – 7.08 (m, 2H), 5.44 (dd, J = 10.1, 4.1 Hz, 1H), 4.44 (dd, J = 11.8, 10.1 Hz, 1H), 3.83 (dd, J = 11.8, 4.1 Hz, 1H); **$^{13}$C NMR** (101 MHz, $CD_3OD$) δ 166.31, 163.86, 131.65, 131.56, 117.08, 116.86, 78.85, 63.30; **HRMS** (ESI-): m/z calc. for $C_8H_8N_2O_3Cl_2$ $[M-H]^-$ 248.98392, found 248.98406.

*(Z)-2-hydroxy-1-(2-hydroxy-1-(4-methoxyphenyl)ethyl)diazene 1-oxide* **(7m)**

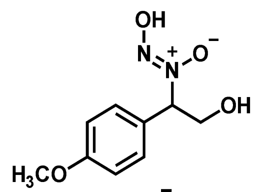

**7m**

Prepared from the corresponding oxime precursor **6m** (400 mg, 2.21 mmol) according to a modified version of the general method E (crude hydroxylamine **3m** was purified by trituration in $Et_2O$, discarding the liquid), followed by the general method G (product **7m** was purified by filtration through a short silica plug, eluting in pure EtOAc).

**3m**: off-white solid (380 mg, 94%); **R*f*** ($SiO_2$; EtOAc:MeOH 95:50) = 0.55 (poorly-UV active, stained with $KMnO_4$); **m.p.** = 118-121°C **IR** (neat, $cm^{-1}$): $\nu_{max}$ = 3260.2, 2934.5, 2837.1, 1611.7, 1584.3, 1513.9, 1462.8, 1442.1, 1303.2, 1246.7, 1179.0, 1113.6, 1029.9, 830.7, 759.1, 553.2; **$^1$H NMR** (400 MHz, $CD_3OD$) δ 7.50 – 7.40 (m, 2H), 7.03 – 6.95 (m, 2H), 4.42 (dd, J = 7.1, 4.7 Hz, 1H), 4.02 (qd, J = 11.8, 6.0 Hz, 2H), 3.81 (s, 3H); **$^{13}$C NMR** (101 MHz, $CD_3OD$) δ 160.90, 132.74, 130.35, 114.97, 69.32, 64.93, 55.87; **HRMS** (ESI+): m/z calc. for $C_9H_{11}O_2$ [M-$OHNH_2$+H]$^+$ 151.07536, found 151.07550.

**7m**: pale yellow oil (100 mg, 30%; starting from 260 mg of **3m**); **IR** (neat, $cm^{-1}$): $\nu_{max}$ = 3357.9, 2932.9, 1611.5, 1515.5, 1462.1, 1306.2, 1252.5, 1179.2, 1061.7, 1031.5, 870.4, 833.3, 535.3; **$^1$H NMR** (400 MHz, $CD_3OD$) δ 7.45 – 7.38 (m, 2H), 6.96 – 6.89 (m, 2H), 5.36 (dd, *J* = 10.2, 4.1 Hz, 1H), 4.44 (dd, *J* = 11.8, 10.3 Hz, 1H), 3.85 – 3.73 (m, 4H); **$^{13}$C NMR** (101 MHz, $CD_3OD$) δ 162.41, 130.75, 126.94, 115.52, 79.41, 63.33, 56.18; **HRMS** (ESI-): m/z calc. for $C_9H_{11}N_2O_4$ [M-H]$^-$ 211.07243, found 211.07241.

*(Z)-1-(1-(furan-2-yl)-2-hydroxyethyl)-2-hydroxydiazene 1-oxide* **(7n)**

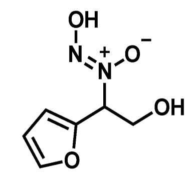

**7n**

A round-bottom flask, equipped with a magnetic stirring bar, was charged with the oxime substrate **6n** (353 mg, 2.50 mmol, 1 eq.) and glacial acetic acid (6.0 mL). Sodium cyanoborohydride (496 mg, 7.50 mmol, 3 eq.) was added portion-wise keeping the temperature at 10 °C. The reaction mixture was allowed to warm to room temperature and stirred for 16 h. The mixture was added to a saturated aqueous solution of $Na_2CO_3$ and extracted three times in *n*-butanol. The combined organic layers were dried over $Na_2SO_4$, filtered and the solvent was removed under reduced pressure. The crude hydroxylamine product **3n** was purified by silica gel flash column chromatography, eluting in $CH_2Cl_2$:MeOH 95:5 to 90:10. **3n**: pale-yellow oil (220 mg, 62%); **R*f*** ($SiO_2$; $CH_2Cl_2$:MeOH 95:5) = 0.35 (non-UV active, stained with $KMnO_4$); **IR** (neat, $cm^{-1}$): $\nu_{max}$ = 3256.4, 2883.3, 1503.5, 1031.8, 1008.9, 738.4; **$^1$H NMR** (400 MHz, $CD_3OD$) δ 7.44 (dt, *J* = 1.7, 0.8 Hz, 1H), 6.39 – 6.29 (m, 2H), 4.08 (t, *J* = 6.4 Hz, 1H), 3.89 – 3.75 (m, 2H); **$^{13}$C NMR** (101 MHz, $CD_3OD$) δ 154.01, 143.07, 111.21, 108.62, 63.40, 61.83; **HRMS** (ESI+): m/z calc. for $C_6H_{10}NO_3$ [M+H$^+$] 144.06552, found 144.06552.

Subsequently, **7n** was prepared following the general method F plus an extra purification step by reversed-phase prep-HPLC (Synergi Hydro-RP column - Phenomenex®, elution gradient from pure $H_2O$ to $H_2O$:$CH_3CN$ 50:50 over 60 min, flow = 12 mL/min, room temperature). Off-white solid (108 mg, 42%; starting from 215 mg of **3n**); **IR** (neat, $cm^{-1}$): $\nu_{max}$ = 3226.8, 3027.3, 2900.7, 2793.8, 1427.1, 1265.4, 1057.4, 1010.6, 746.6; **$^1$H NMR** (400 MHz, $CD_3OD$) δ 7.55 (dd, *J* = 1.9, 0.7 Hz, 1H), 6.58 (dt, *J* = 3.4, 0.7 Hz, 1H), 6.45 (dd, *J* = 3.3, 1.9 Hz, 1H), 5.54 (dd, *J* = 9.8, 4.3 Hz, 1H), 4.43 (dd, *J* = 11.8, 9.9 Hz, 1H), 4.01 (dd, *J* = 11.8, 4.3 Hz, 1H); **$^{13}$C NMR** (101 MHz, $CD_3OD$) δ 147.55, 144.72, 111.75, 111.37, 72.91, 60.75; **HRMS** (ESI-): m/z calc. for $C_6H_7N_2O_4$ [M-H]$^-$ 171.04113, found 171.04121.

*(Z)-2-hydroxy-1-(1-hydroxy-3-(thiophen-2-yl)propan-2-yl)diazene 1-oxide* **(7o)**

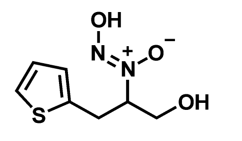


7o

Prepared from the corresponding oxime precursor **6o** (205 mg, 1.20 mmol) following the general methods D and F. **7o**: colourless solid (205 mg, 88%); **m.p.** = 78-79 °C; **IR** (neat, $cm^{-1}$): $\nu_{max}$ = 3362.4, 2936.3, 1443.4, 1070.0, 1037.2, 700.4; **$^1$H NMR** (400 MHz, $CD_3OD$) δ 7.23 (dd, *J* = 5.2, 1.2 Hz, 1H), 6.91 (dd, *J* = 5.2, 3.5 Hz, 1H), 6.85 (dd, *J* = 3.5, 1.0 Hz, 1H), 4.50 (ddt, *J* = 10.0, 8.7, 4.3 Hz, 1H), 3.98 (dd, *J* = 11.8, 8.9 Hz, 1H), 3.78 (dd, *J* = 11.8, 4.1 Hz, 1H), 3.37 (ddd, *J* = 15.2, 10.0, 0.8 Hz, 1H), 3.20 (ddd, *J* = 15.2, 4.5, 0.9 Hz, 1H); **$^{13}$C NMR** (101 MHz, $CD_3OD$) δ 139.07, 127.94, 127.56, 125.58, 78.07, 62.42, 29.55; **HRMS** (ESI-): m/z calc. for $C_7H_9N_2O_3S$ [M-H]$^-$ 201.03394, found 201.03389.

*2-(4-fluorophenyl)-N,3-dihydroxypropanamide* **(9)**

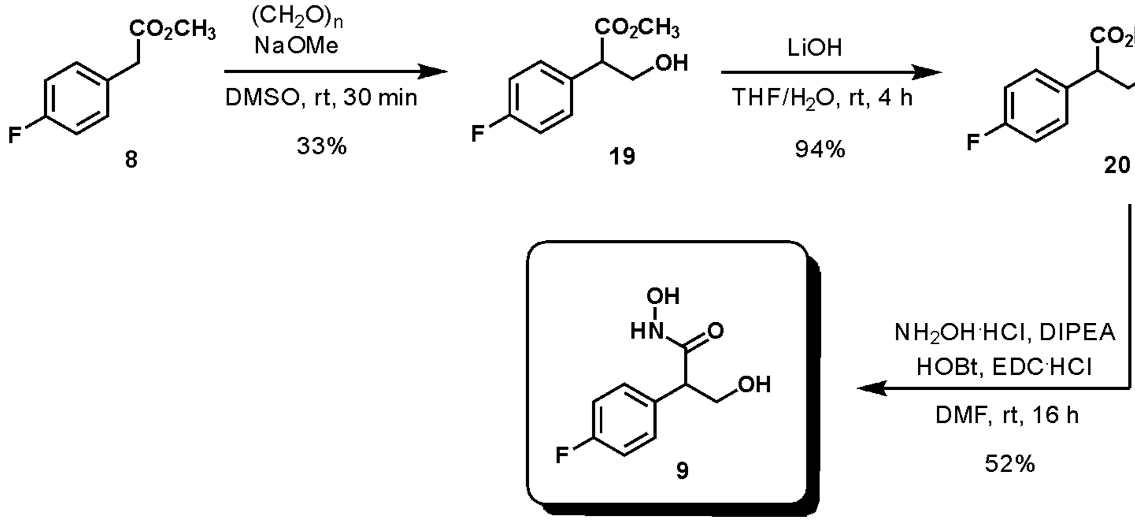


Step 1) Experimental procedure adapted from the literature.[9] A flame-dried round-bottom flask, equipped with a magnetic stirring bar, was charged with commercial methyl 4-fluorophenylacetate **8** (2.02 g, 12.00 mmol) and anhydrous DMSO (24 mL) under an atmosphere of argon. The resulting solution was cooled to 0 °C. Paraformaldehyde (324 mg, 10.8 mmol) and sodium methoxide (130 mg, 2.40 mmol) were added. The reaction mixture was allowed to warm to room temperature and stirred for further 30 min. The reaction was quenched with an aqueous saturated solution of $NH_4Cl$ and then diluted with water. The aqueous layer was extracted three times in EtOAc. The combined organic layers were washed twice with brine, dried over $Na_2SO_4$, filtered, and concentrated under reduced pressure. The crude product **19** was purified by flash column chromatography ($SiO_2$; *n*-pent:EtOAc 90:10 to 60:40). **19**: colourless oil (700 mg, 33%); $\mathbf{R}_f$ ($SiO_2$; *n*-pent:EtOAc 80:20) = 0.15 (poorly-UV active, stained in $KMnO_4$); The spectral data matched the one described in the literature.[10]

Step 2) A round-bottom flask, equipped with a magnetic stirring bar, was charged with compound **19** (396 mg, 2.00 mmol) and THF (6 mL). The resulting solution was cooled to 0 °C. A solution of lithium hydroxide monohydrate (120 mg, 5 mmol) in water (2 mL) was added dropwise. The reaction mixture was allowed to warm to room temperature and stirred for further 4 h. A 2 N HCl aqueous solution was added until pH 3. The aqueous layer was extracted three times in EtOAc. The combined organic layers were washed with brine, dried over $Na_2SO_4$, filtered, and concentrated under reduced pressure. The crude product **20** was used in the next step without further purification. **20**: colourless solid (345 mg, 94%); **m.p.** = 82-84 °C; **IR** (neat, $cm^{-1}$): $\nu_{max}$ = 3700-2200 (broad peak), 1710.7, 1509.6, 1226.6; **$^1$H NMR** (400 MHz, $CD_3OD$): δ 7.41 – 7.29 (m, 2H), 7.12 – 6.99 (m, 2H), 4.13 – 3.99 (m, 1H), 3.79 – 3.65 (m, 2H); **$^{13}$C NMR** (101 MHz, $CD_3OD$) δ 175.89, 163.62 (d, $J$ = 244.4 Hz), 134.02 (d, $J$ = 3.4 Hz), 131.10 (d, $J$ = 7.9 Hz), 116.26 (d, $J$ = 21.7 Hz), 65.04, 54.98; **$^{19}$F NMR** (377 MHz, $CD_3OD$) δ -117.51; **HRMS** (ESI-): m/z calc. for $C_9H_8O_3F$ $[M-H]^-$ 183.04630, found 183.04621.

Step 3) A round-bottom flask, equipped with a magnetic stirring bar, was charged with compound **20** (147 mg, 0.80 mmol). A nitrogen atmosphere was created and anhydrous DMF (5 mL) was added. 1-Hydroxybenzotriazole hydrate (130 mg, 0.96 mmol) and 1-(3-dimethylaminopropyl)-3-ethyl-carbodiimide hydrochloride (184 mg, 0.96 mmol) were added. After 1 h of stirring at room temperature, hydroxylamine hydrochloride (67 mg, 0.96 mmol) and DIPEA (0.35 mL, 2.00 mmol) were added, and the stirring was continued at room temperature for 16 h. The reaction was diluted with brine and extracted twice in *n*-BuOH. The combined organic layers were dried over $Na_2SO_4$, filtered, and concentrated under reduced pressure. The crude product **9** was purified by flash column chromatography ($SiO_2$; $CH_2Cl_2$:MeOH 95:5 to 90:10). **9**: colourless solid (82 mg, 52%); **R*f*** ($SiO_2$; $CH_2Cl_2$:MeOH 90:10) = 0.5; **m.p.** = 165-166 °C; **IR** (neat, $cm^{-1}$): $\nu_{max}$ = 3338.3, 3172.3, 3007.1, 2891.3, 1630.8, 1509.3, 1230.6, 1055.4, 1017.5, 835.2; **$^1$H NMR** (400 MHz, $CD_3OD$) δ 7.44 – 7.31 (m, 2H), 7.09 – 6.98 (m, 2H), 4.10 (dd, *J* = 10.6, 9.0 Hz, 1H), 3.70 (dd, *J* = 10.6, 5.7 Hz, 1H), 3.51 (dd, *J* = 9.1, 5.7 Hz, 1H); **$^{13}$C NMR** (101 MHz, $CD_3OD$) δ 171.58, 163.60 (d, *J* = 244.2 Hz), 134.37 (d, *J* = 3.2 Hz), 130.93 (d, *J* = 8.1 Hz), 116.16 (d, *J* = 21.6 Hz), 64.62, 52.48; **$^{19}$F NMR** (377 MHz, $CD_3OD$) δ -117.57; **HRMS** (ESI+): m/z calc. for $C_9H_{11}O_3NF$ $[M+H]^+$ 200.07175, found 200.07185.

*N-hydroxy-N-(1-hydroxy-4-methylpentan-2-yl)benzamide* **(10)**

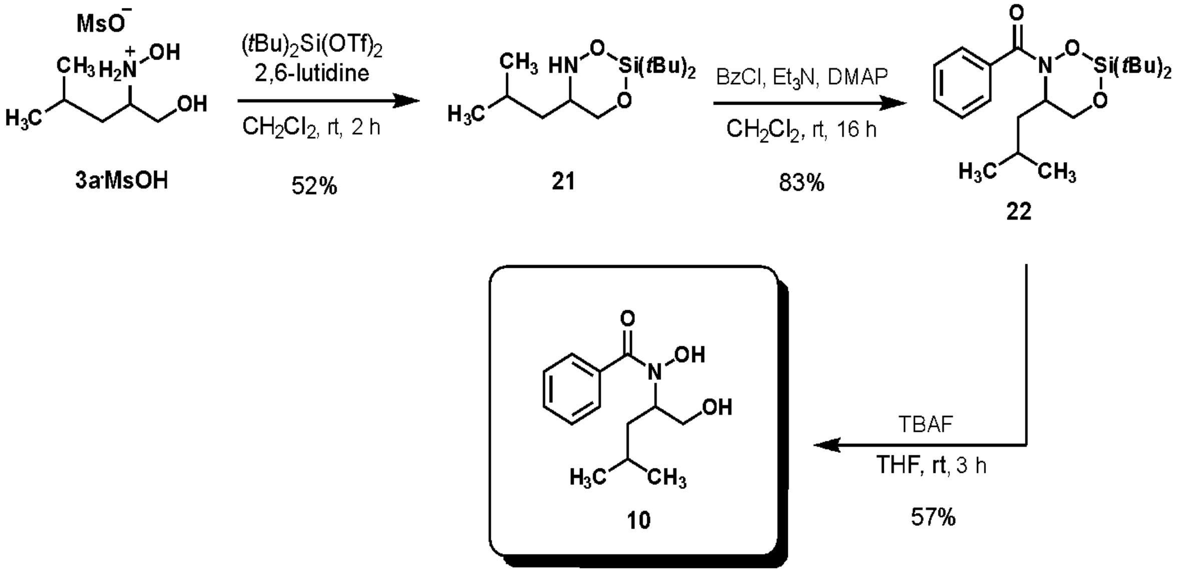


Step 1) A flame-dried round-bottom flask, equipped with a magnetic stirring bar, was charged with hydroxylammonium methanesulfonate **3a·MsOH** (434 mg, 1.80 mmol). An atmosphere of nitrogen was created. Anhydrous $CH_2Cl_2$ (18 mL) was added and the resulting solution was cooled to –5 °C (salt/ice/water bath). 2,6-lutidine (0.73 mL, 6.30 mmol) and di-*tert*-butyldichlorosilane (0.65 mL, 1.98 mmol) were added. The reaction mixture was allowed to warm to room temperature and further stirred for 2 h. The reaction was quenched with an saturated aqueous solution of $NaHCO_3$. The aqueous layer was extracted twice in $CH_2Cl_2$. The combined organic layers were washed with brine. The organic layer was dried over $Na_2SO_4$, filtered, and concentrated under reduced pressure. The crude product **21** was purified by flash column chromatography ($SiO_2$; *n*-pent:EtOAc 99:1 to 95:5). **21**: colourless oil (256 mg, 52%); **R*f*** ($SiO_2$; *n*-pent:EtOAc 95:5) = 0.4 (non-UV active, stained in $KMnO_4$); **$^1$H NMR** (400 MHz, $CDCl_3$): δ 5.48 (br. s, 1H), 3.93 (dd, *J* = 10.4, 3.6 Hz, 1H), 3.77 (t, *J* = 10.4 Hz, 1H), 3.32 (tdd, *J* = 10.8, 5.2, 3.6 Hz, 1H), 1.68 – 1.50

(m, 1H), 1.14 – 0.96 (m, 20H), 0.92 (d, $J$ = 6.5 Hz, 6H); **$^{13}$C NMR** (101 MHz, $CDCl_3$) δ 70.88, 59.43, 37.58, 28.30, 27.33, 25.00, 23.22, 22.69, 22.46, 20.02.

Step 2) A flame-dried round-bottom flask, equipped with a magnetic stirring bar, was charged with compound **21** (109 mg, 0.40 mmol). Anhydrous $CH_2Cl_2$ (2 mL) was added and the resulting solution was cooled to 0 °C. Triethylamine (0.11 mL, 0.80 mmol), 4-dimethylaminopyridine (5 mg, 0.04 mmol), and benzoyl chloride (57 µL, 0.48 mmol) were added. The reaction mixture was allowed to warm to room temperature and stirred for further 16 h. The reaction was diluted with a saturated aqueous solution of $NaHCO_3$. The aqueous layer was extracted twice in $CH_2Cl_2$. The combined organic layers were dried over $Na_2SO_4$, filtered, and concentrated under reduced pressure. The crude product **22** was purified by flash column chromatography ($SiO_2$; *n*-pent:$Et_2O$ 90:10 to 70:30). **22**: colourless oil (125 mg, 83%); **R*f*** ($SiO_2$; *n*-pent:$Et_2O$ 80:20) = 0.2; **IR** (neat, cm$^{-1}$): $\nu_{max}$ = 2948.9, 2862.8, 1636.4, 1474.7, 1407.7, 1125.7, 1052.2, 826.4; **$^{1}$H NMR** (400 MHz, $CDCl_3$): δ 7.86 – 7.67 (br. m, 2H), 7.47 – 7.32 (m, 3H), 4.84 (br. s, 1H), 4.51 (dd, $J$ = 11.3, 4.0 Hz, 1H), 3.92 (dd, $J$ = 11.3, 1.4 Hz, 1H), 2.35 (ddd, $J$ = 14.4, 9.8, 5.0 Hz, 1H), 1.69 (br. s, 1H), 1.54 – 1.38 (m, 1H), 1.08 (s, 9H), 1.02 – 0.68 (br. m, 15H); **$^{13}$C NMR** (101 MHz, $CDCl_3$) δ 167.80, 134.25, 130.74, 129.02, 127.87, 67.96, 38.86, 27.82, 27.46, 25.12, 23.28, 22.70, 22.31, 20.71.

Step 3) A flame-dried round-bottom flask, equipped with a magnetic stirring bar, was charged with compound **22** (76 mg, 0.20 mmol). A nitrogen atmosphere was created and anhydrous THF (1 mL) was added. The resulting solution was cooled to 0 °C and TBAF (0.5 mL, 0.5 mmol, 1 M solution in THF) was added dropwise. The reaction mixture was allowed to warm to room temperature and stirred for further 3 h. The reaction was diluted with EtOAc and water. The aqueous layer was extracted three times in EtOAc. The combined organic layers were dried over $Na_2SO_4$, filtered, and concentrated under reduced pressure. The crude product **10** was purified by flash column chromatography ($SiO_2$; *n*-pent:EtOAc 60:40 to 40:60). **10**: orange oil (27 mg, 57%); **R*f*** ($SiO_2$; *n*-*n*-pent:EtOAc 50:50) = 0.2; **IR** (neat, cm$^{-1}$): $\nu_{max}$ = 3207.4, 2955.4, 1609.6, 1597.1, 1572.7, 1447.5, 703.0; **$^{1}$H NMR** (500 MHz, $CD_2Cl_2$) δ 7.63 – 7.55 (br. m, 2H), 7.54 – 7.40 (br. m, 3H), 4.20 – 4.04 (br. m, 1H), 3.98 – 3.82 (br. m, 1H), 3.55 – 3.35 (br. m, 1H), 1.80 – 1.70 (br. m, 1H), 1.61 – 1.47 (br. m, 1H), 1.09 – 0.78 (br. m, 5H), 0.60 (br. d, $J$ = 6.3 Hz, 3H); **$^{13}$C NMR** (126 MHz, $CD_2Cl_2$) δ 170.48, 133.79, 130.84, 128.82, 128.43, 62.22, 62.08, 37.01, 24.72, 23.23, 22.03; **HRMS** (ESI+): m/z calc. for $C_{13}H_{20}O_3N$ $[M+H]^+$ 238.14377, found 238.14375.

*(Z)-1-(1-(4-fluorophenyl)-2-hydroxyethyl)-2-methoxydiazene 1-oxide* **(11)**

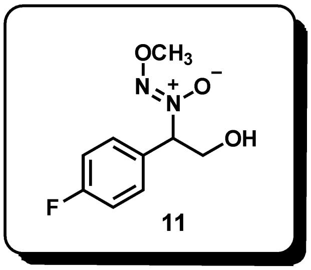


A flame-dried round-bottom flask, equipped with a magnetic stirring bar, was charged with the diazeniumdiolate **7j** (360 mg, 1.80 mmol). An atmosphere of nitrogen was created. Anhydrous MeOH (8.0 mL) and $NaHCO_3$ (227 mg, 2.70 mmol) were added. The mixture was cooled to 0 °C and dimethyl sulfate (0.20 mL, 2.16 mmol) was added dropwise. The reaction mixture was stirred at 0 °C for 15 min and then allowed to warm to room temperature. After 16 h, the mixture was diluted with water and extracted three times in EtOAc. The combined organic layers were dried over $Na_2SO_4$, filtered, and concentrated under reduced pressure. The crude product was purified by flash column chromatography ($SiO_2$; *n*-pent:EtOAc 60:40 to 30:70). **11**: Colourless solid (270 mg, 70%); **R*f*** ($SiO_2$; *n*-pent:EtOAc 50:50) = 0.15; **m.p.** = 82-83 °C; **IR** (neat, $cm^{-1}$): $\nu_{max}$ = 3416.8, 2947.6, 1605.8, 1511.2, 1228.1, 1058.1, 1012.6; **$^1$H NMR** (400 MHz, $CDCl_3$): δ = 7.49 – 7.41 (m, 2H), 7.11 – 7.01 (m, 2H), 5.38 (dd, *J* = 9.7, 3.8 Hz, 1H), 4.50 (dd, *J* = 12.2, 9.7 Hz, 1H), 4.06 (s, 3H), 3.87 (dd, *J* = 12.2, 3.9 Hz, 1H), 2.66 (s, 1H); **$^{13}$C NMR** (101 MHz, $CDCl_3$) δ = 163.45 (d, *J* = 249.1 Hz), 129.72 (d, *J* = 8.4 Hz), 128.83 (d, *J* = 3.3 Hz), 116.02 (d, *J* = 21.8 Hz), 77.75, 62.97, 61.62; **$^{19}$F NMR** (377 MHz, $CDCl_3$) δ -111.59; **HRMS** (ESI+): m/z calc. for $C_9H_{11}FN_2O_3Na$ $[M+Na]^+$ 237.06459, found 237.06441.

*(Z)-1-(1-(4-fluorophenyl)-2-methoxyethyl)-2-hydroxydiazene 1-oxide* **(12)**

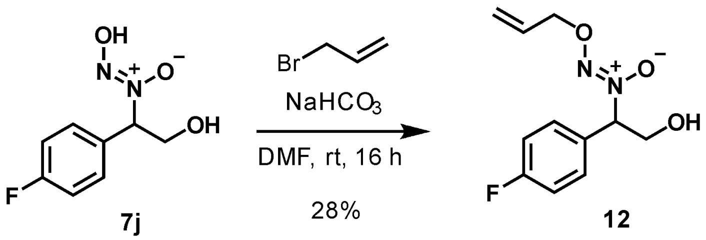


A flame-dried round-bottom flask, equipped with a magnetic stirring bar, was charged with the diazeniumdiolate **7j** (2.00 g, 10.00 mmol). An atmosphere of nitrogen was created. Anhydrous DMF (10 mL) and $NaHCO_3$ (1.00 g, 12.00 mmol) were added. The mixture was cooled to 0 °C and allyl bromide (0.91 mL, 10.50 mmol) was added dropwise. The mixture was allowed to warm to room temperature and stirred for 16 h. The reaction mixture was diluted with $H_2O$ (50 mL) and extracted three times in EtOAc (3x 50 mL). The combined organic layers were washed twice with a solution of 5% LiCl in $H_2O$, dried over $Na_2SO_4$, filtered, and concentrated under reduced pressure. The crude product **12** was purified by flash column chromatography ($SiO_2$; *n*-pent:EtOAc 70:30 to 40:60). **12**: Colourless solid (660 mg, 28%); **R*f*** ($SiO_2$; *n*-pent:EtOAc 50:50) = 0.30; **m.p.** = 66-67 °C; **IR** (neat, $cm^{-1}$): $\nu_{max}$ = 3421.3, 2937.3, 1606.0, 1511.3, 1228.7, 1054.2, 1014.6, 989.7; **$^1$H NMR** (400 MHz, $CDCl_3$): δ = 7.48 – 7.41 (m, 2H), 7.10 – 7.01 (m, 2H), 5.97 (ddt, *J* = 17.1,

10.4, 6.0 Hz, 1H), 5.42 – 5.25 (m, 3H), 4.75 (dt, $J$ = 6.0, 1.3 Hz, 2H), 4.50 (dd, $J$ = 12.2, 9.6 Hz, 1H), 3.88 (dd, $J$ = 12.3, 3.8 Hz, 1H), 2.46 (s, 1H); **$^{13}$C NMR** (101 MHz, $CDCl_3$) δ = 163.43 (d, $J$ = 248.8 Hz), 131.94, 129.70 (d, $J$ = 8.4 Hz), 128.88 (d, $J$ = 3.4 Hz), 120.05, 116.02 (d, $J$ = 21.7 Hz), 77.84, 75.14, 63.07; **$^{19}$F NMR** (377 MHz, $CDCl_3$) δ -111.66; **HRMS** (ESI+): m/z calc. for $C_{11}H_{13}FN_2O_3Na$ [M+Na$^+$] 263.08024, found 263.08005.

*(Z)-1-(1-(4-fluorophenyl)-2-methoxyethyl)-2-hydroxydiazene 1-oxide* **(13)**

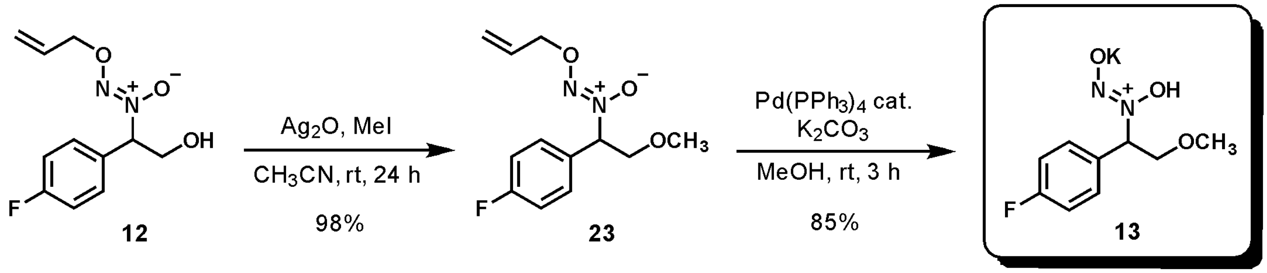


Step 1) A flame-dried round-bottom flask, equipped with a magnetic stirring bar, was charged with the primary alcohol **12** (96 mg, 0.40 mmol). An atmosphere of nitrogen was created. Anhydrous acetonitrile (4 mL), $Ag_2O$ (463 mg, 2.00 mmol) and iodomethane (0.25 mL, 4.00 mmol) were added successively. The flask was protected from light and the reaction mixture was stirred vigorously at room temperature for 24 h. The mixture was filtered through a short pad of Celite and concentrated under reduced pressure. The crude product **23** was purified by flash column chromatography ($SiO_2$; *n*-pent:EtOAc 90:10 to 70:30). **23**: Colourless oil (100 mg, 98%); **$R_f$** ($SiO_2$; *n*-pent:EtOAc 70:30) = 0.4; **IR** (neat, cm$^{-1}$): $\nu_{max}$ = 2932.5, 1606.4, 1511.3, 1227.6, 1127.4, 1011.4; **$^1$H NMR** (400 MHz, $CDCl_3$): δ 7.52 – 7.42 (m, 2H), 7.10 – 6.99 (m, 2H), 5.98 (ddt, $J$ = 17.3, 10.4, 5.9 Hz, 1H), 5.40 (dd, $J$ = 10.1, 3.9 Hz, 1H), 5.37 – 5.21 (m, 2H), 4.75 (dq, $J$ = 6.0, 1.2 Hz, 2H), 4.31 (t, $J$ = 10.3 Hz, 1H), 3.66 (dd, $J$ = 10.4, 3.9 Hz, 1H), 3.41 (s, 3H); **$^{13}$C NMR** (101 MHz, $CDCl_3$) δ 163.49 (d, $J$ = 249.1 Hz), 132.16, 129.93 (d, $J$ = 8.6 Hz), 128.77 (d, $J$ = 3.4 Hz), 119.65, 115.93 (d, $J$ = 21.7 Hz), 75.87, 74.91, 71.80, 59.41; **$^{19}$F NMR** (377 MHz, $CDCl_3$) δ -111.66; **HRMS** (ESI+): m/z calc. for $C_{12}H_{15}FN_2O_3Na$ [M+Na$^+$] 277.09589, found 277.09607.

Step 2) A flame-dried round-bottom flask, equipped with a magnetic stirring bar, was charged with **23** (89 mg, 0.35 mmol). An argon atmosphere was created and anhydrous MeOH (7 mL) was added. The solution was degassed by bubbling argon for 5 min, and tetrakis(triphenylphosphine)palladium(0) ($Pd(PPh_3)_4$, 8.1 mg 7.0 µmol, 2 mol%) was added. The resulting yellow solution was stirred at room temperature for 5 min. $K_2CO_3$ (290 mg, 2.10 mmol) was added and the reaction was stirred at room temperature for 3 h. The solvent was removed under reduced pressure. EtOH (5 mL) was added to dissolve the product and the emulsion ($K_2CO_3$ is insoluble in EtOH) was filtered through a 1 g Discovery SPE-18 column, washing in EtOH (4x 4 mL). The filtrate was concentrated under reduced pressure and the residue was triturated with acetonitrile (3x 5 mL). Concentration of the liquid under reduced pressure afforded the pure product **13**. Colourless oil (75 mg, 85%); **IR** (neat, cm$^{-1}$): $\nu_{max}$ = 3368.5, 1663.4, 1605.2, 1510.9, 1228.3, 1100.5; **$^1$H NMR** (400 MHz, $CD_3OD$) δ 7.56 – 7.47 (m, 2H), 7.13 – 7.02 (m, 2H), 5.46 (dd, $J$ = 9.4, 4.8 Hz, 1H), 4.28 (td, $J$ = 9.8, 0.9 Hz, 1H), 3.76 – 3.68 (m, 1H), 3.38 (d, $J$ = 0.9 Hz, 3H); **$^{13}$C NMR** (101 MHz, $CD_3OD$) δ 164.12 (d, $J$ = 244.8 Hz), 133.29 (d, $J$ = 3.2 Hz), 130.89 (d,

$J$ = 8.2 Hz), 116.08 (d, $J$ = 21.7 Hz), 73.11, 71.96, 59.04; $^{19}$**F NMR** (377 MHz, $CD_3OD$) δ -116.23; **HRMS** (ESI-): m/z calc. for $C_9H_{10}O_3N_2ClFK$ [M+$Cl^-$] 287.00065, found 287.00056.

*(Z)-1-(1-(4-fluorophenyl)-2-mercaptoethyl)-2-hydroxydiazene 1-oxide* **(14)**

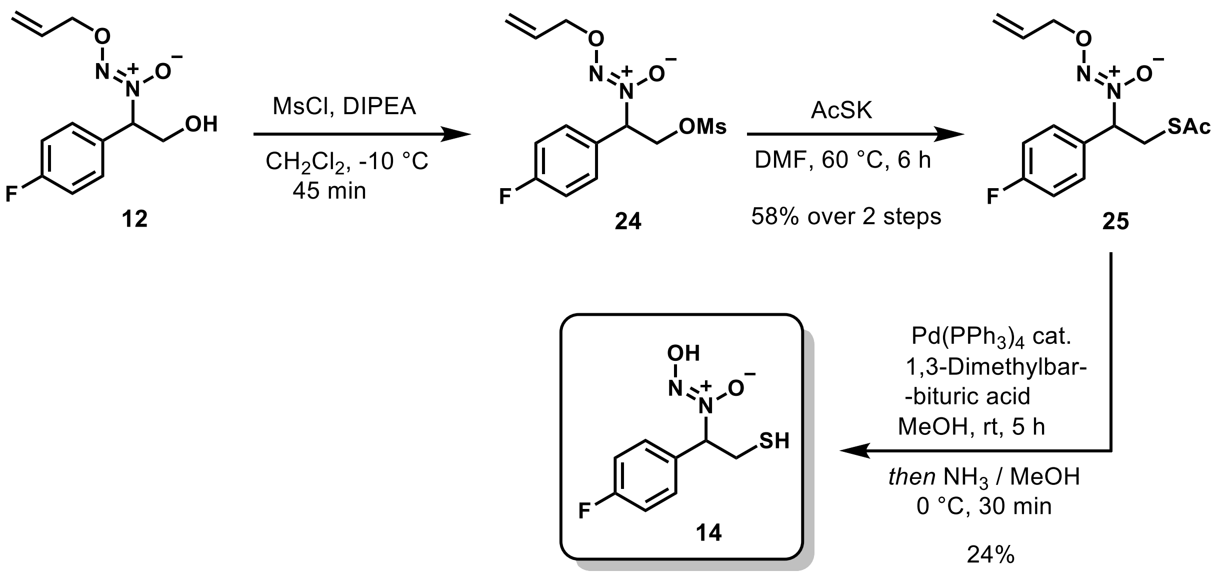


Step 1) A flame-dried round-bottom flask, equipped with a magnetic stirring bar, was charged with compound **12** (180 mg, 0.75 mmol). Anhydrous $CH_2Cl_2$ (3 mL) was added and the resulting solution was cooled to -10 °C (salt/ice/water bath). *N,N*-Diisopropylethylamine (0.14 mL, 0.825 mmol) was added, followed by dropwise addition of methanesulfonyl chloride (61 µL, 0.79 mmol). The reaction was stirred at -10 °C for 45 min before quenching with water (10 mL). The layers were separated, and the aqueous layer was extracted twice in $CH_2Cl_2$. The combined organic layers were washed with a saturated aqueous solution of $NaHCO_3$, and brine. The organic layer was dried over $Na_2SO_4$, filtered, and concentrated under reduced pressure to afford the crude product **24**, which was used in the next step without further purification.

Step 2) A flame-dried glass tube, equipped with a magnetic stirring bar, was charged with crude **24** (111 mg, 0.35 mmol theoretical). An atmosphere of nitrogen was created. Anhydrous DMF (1.0 mL) and potassium thioacetate (160 mg, 1.4 mmol) were added successively. The tube was sealed, and the reaction mixture was heated to 60 °C for 6 h. Upon cooling to room temperature, the solution was diluted with water and extracted twice in EtOAc. The combined organic layers were washed twice with LiCl (5%, aqueous), once with brine, dried over $Na_2SO_4$, filtered, and concentrated under reduced pressure. The crude product **25** was purified by flash column chromatography ($SiO_2$; *n*-pent:EtOAc 95:5 to 80:20). **25**: brown oil (60 mg, 58% over 2 steps); $\mathbf{R_f}$ ($SiO_2$; *n*-pent:EtOAc 80:20) = 0.4; **IR** (neat, $cm^{-1}$): $\nu_{max}$ = 2937.1, 1693.2, 1510.2, 1226.4, 1132.8, 1011.9; $^{1}$**H NMR** (400 MHz, $CDCl_3$): δ 7.53 – 7.46 (m, 2H), 7.11 – 7.02 (m, 2H), 5.96 (dddd, $J$ = 17.5, 10.4, 6.2, 5.6 Hz, 1H), 5.37 – 5.24 (m, 3H), 4.74 (dd, $J$ = 5.9, 1.4 Hz, 2H), 3.67 (dd, $J$ = 14.2, 10.1 Hz, 1H), 3.52 (dd, $J$ = 14.1, 4.9 Hz, 1H), 2.36 (d, $J$ = 0.6 Hz, 3H); $^{13}$**C NMR** (101 MHz, $CDCl_3$) δ 195.35, 163.48 (d, $J$ = 248.8 Hz), 132.04, 130.70 (d, $J$ = 3.3 Hz), 129.55 (d, $J$ = 8.7 Hz),

119.77, 116.00 (d, $J$ = 21.8 Hz), 75.60, 75.02, 31.66, 30.73; **$^{19}$F NMR** (377 MHz, $CDCl_3$) δ -111.49; **HRMS** (ESI+): m/z calc. for $C_{13}H_{15}FN_2O_3NaS$ $[M+Na]^+$ 321.06796, found 321.06810.

Step 3) A flame-dried round-bottom flask, equipped with a magnetic stirring bar, was charged with **25** (140 mg, 0.47 mmol). An argon atmosphere was created and anhydrous MeOH (9 mL), and 1,3-dimethyl barbituric acid (146 mg, 0.94 mmol) were successively added. The solution was degassed by bubbling argon for 5 min, and tetrakis(triphenylphosphine)palladium(0) ($Pd(PPh_3)_4$, 28 mg, 0.02 mmol, 5 mol%) was added. The reaction mixture was stirred at room temperature for 5 h. The crude was filtered through a short pad of Celite® with the aid of EtOAc. The filtrate was concentrated *in vacuo* and redissolved in dry methanol (5 mL) under an argon atmosphere. The solution was cooled to 0 °C and treated with 7N ammonia in methanol (2.5 mL). The reaction was stirred at 0 °C for 30 minutes. The reaction was concentrated *in vacuo*, resuspended in 50% $CH_3CN$ in $H_2O$ (5 mL), sonicated and filtered through a Discovery® DSC-18 SPE cartridge (bed weight 200 mg) with the aid of additional 50% $CH_3CN$ in $H_2O$ (15 mL). The filtrate was concentrated *in vacuo* and purified by reversed phase prep-HPLC (Synergi Hydro-RP column - Phenomenex®, elution gradient from pure $H_2O$ to 50% $CH_3CN$ in $H_2O$ over 35 min and another 5 min at 50% $CH_3CN$ in $H_2O$, flow = 15 mL/min, room temperature) to afford the pure product **14**. Colourless oil (24 mg, 24%); **IR** (neat, $cm^{-1}$): $\nu_{max}$ = 3307.1, 1599.9, 1509.6, 1222.2, 1159.2; **$^1$H NMR** (500 MHz, $CD_3OD$) δ 7.55 – 7.47 (m, 2H), 7.10 – 7.01 (m, 2H), 5.31 (dd, $J$ = 9.8, 5.2 Hz, 1H), 3.50 (dd, $J$ = 13.8, 9.8 Hz, 1H), 2.95 (dd, $J$ = 13.8, 5.2 Hz, 1H); **$^{13}$C NMR** (126 MHz, $CD_3OD$) δ 164.08 (d, $J$ = 245.4 Hz), 135.22 (d, $J$ = 3.3 Hz), 130.64 (d, $J$ = 8.2 Hz), 116.04 (d, $J$ = 21.7 Hz), 76.14, 27.98; **$^{19}$F NMR** (377 MHz, $CD_3OD$) δ -116.29; **HRMS** (ESI-): m/z calc. for $C_8H_8O_2N_2FS$ $[M-H]^-$ 215.02960, found 215.02980.

*(E)-1-(4-fluorophenyl)ethan-1-one oxime* **(15)**

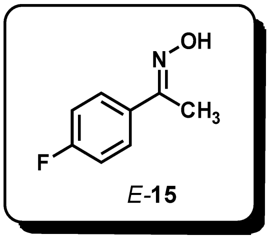


Prepared from commercial 4'-fluoroacetophenone (1.4 mL, 11.2 mmol) following the general method C. **15**: colourless solid (1.65 g, 96%); **R*f*** ($SiO_2$; *n*-pent:EtOAc 80:20) = 0.4; The spectral data matched the one reported in the literature.[4]

*(Z)-1-(1-(4-fluorophenyl)ethyl)-2-hydroxydiazene 1-oxide* **(16)**

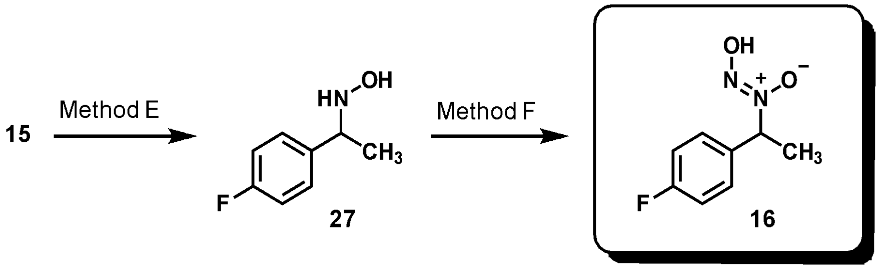


Step 1) Starting from oxime **15** (536 mg, 3.50 mmol), the corresponding hydroxylamine intermediate **27** was prepared according to a modified version of the general method E (crude product was purified by silica gel flash column chromatography, eluting in *n*-pent:EtOAc 50:50 to pure EtOAc). **27**: colourless solid (510 mg, 94%); **R*f*** ($SiO_2$; EtOAc) = 0.50 (poorly-UV active, stained with $KMnO_4$); **m.p.** = 78-79 °C; **IR** (neat, $cm^{-1}$): $\nu_{max}$ = 3247.6, 2974.8, 1604.2, 1509.9, 1224.6, 832.3; **$^1$H NMR** (400 MHz, $CD_3OD$) δ 7.42 – 7.32 (m, 2H), 7.08 – 6.98 (m, 2H), 4.04 (q, *J* = 6.7 Hz, 1H), 1.32 (d, *J* = 6.7 Hz, 3H); **$^{13}$C NMR** (101 MHz, $CD_3OD$) δ 163.47 (d, *J* = 243.4 Hz), 140.28 (d, *J* = 3.1 Hz), 130.10 (d, *J* = 8.0 Hz), 115.88 (d, *J* = 21.3 Hz), 62.26, 19.93; **$^{19}$F NMR** (377 MHz, $CD_3OD$) δ -118.02; **HRMS** (ESI+): m/z calc. for $C_8H_{11}FNO$ [M+H$^+$] 156.08192, found 156.08172.

Step 2) The title compound **16** was prepared from pure **27** (132 mg, 0.85 mmol) according to a modified general method F (after concentration of the reaction mixture, the obtained residue was dissolved in acetonitrile and washed three times with *n*-hexane). **16**: yellow oil (147 mg, 94%, contains 5% of hydroxylamine **27**); **IR** (neat, $cm^{-1}$): $\nu_{max}$ = 2994.0, 1605.4, 1512.5, 1453.0, 1230.1, 1049.5, 840.7; **$^1$H NMR** (400 MHz, $CD_3OD$) δ 7.56 – 7.48 (m, 2H), 7.15 – 7.07 (m, 2H), 5.57 (q, *J* = 7.0 Hz, 1H), 1.79 (d, *J* = 7.0 Hz, 3H); **$^{13}$C NMR** (101 MHz, $CD_3OD$) δ 164.44 (d, *J* = 246.5 Hz), 134.46 (d, *J* = 3.3 Hz), 130.56 (d, *J* = 8.6 Hz), 116.41 (d, *J* = 22.0 Hz), 72.23, 18.92; **$^{19}$F NMR** (376 MHz, $CD_3OD$) δ -115.03; **HRMS** (ESI-): m/z calc. for $C_8H_8FN_2O_2$ [M-H$^+$] 183.05753, found 183.05723.

NMR-spectra

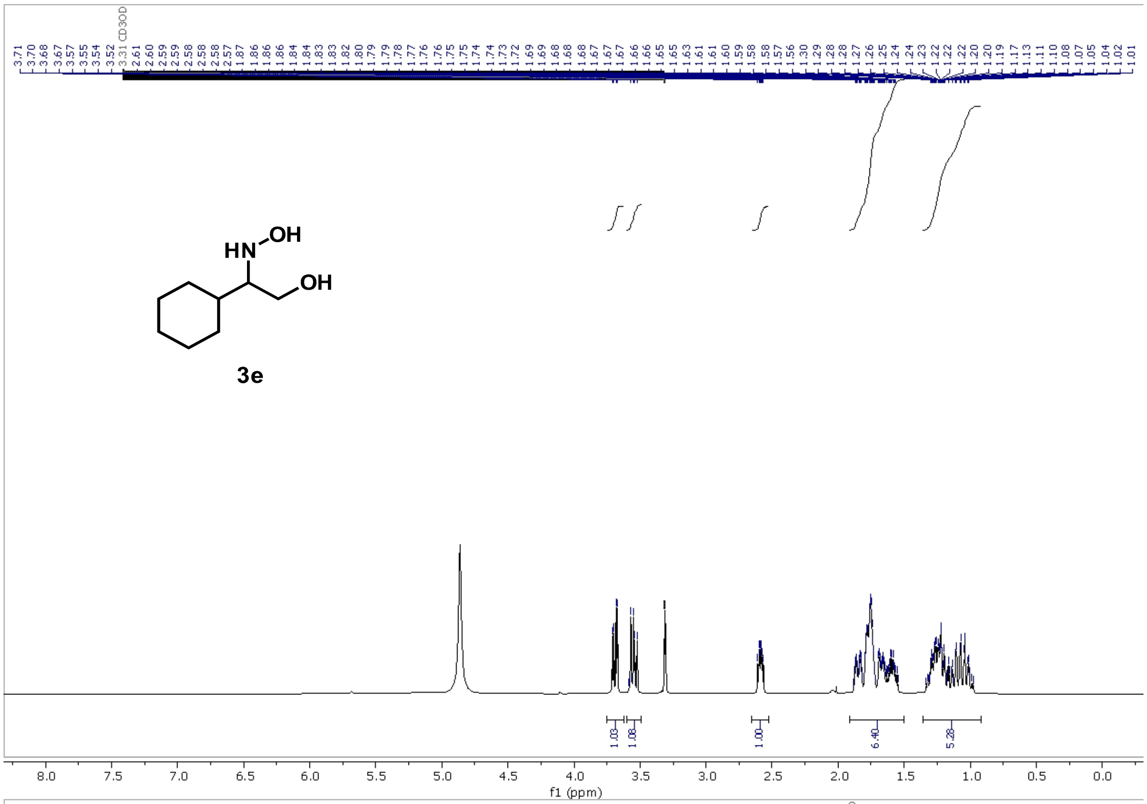

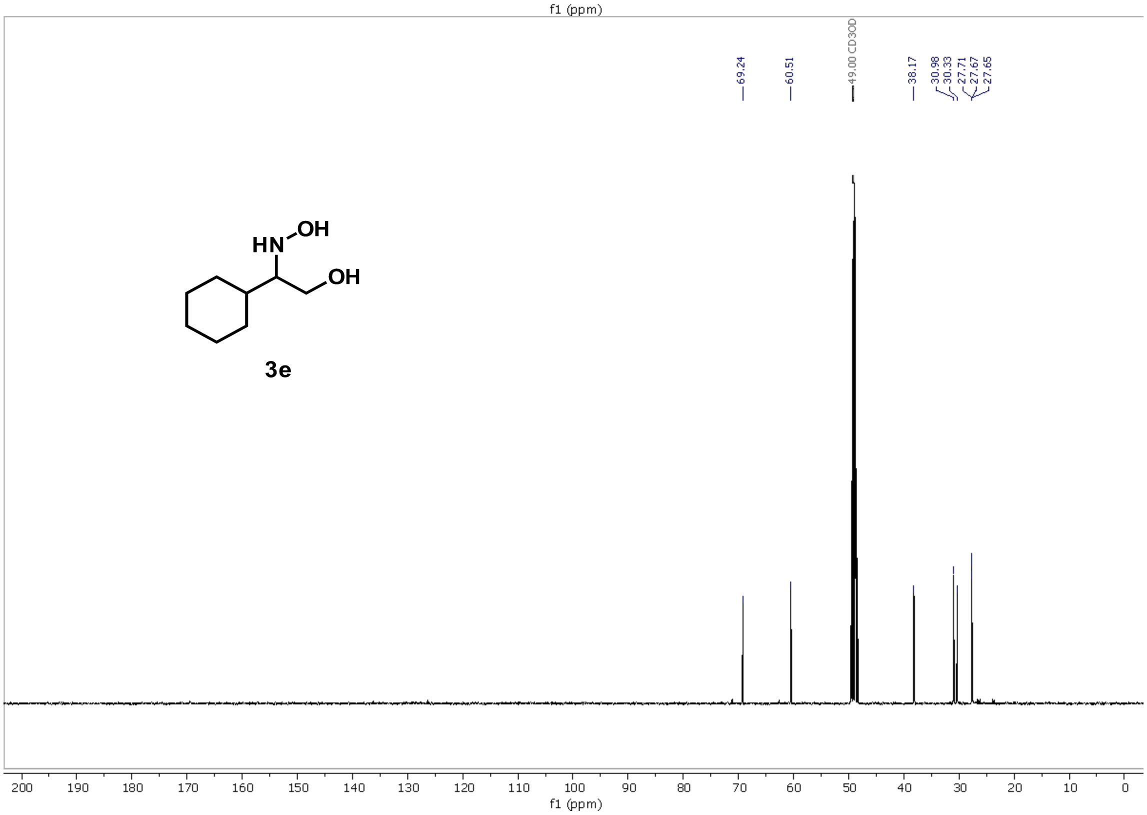

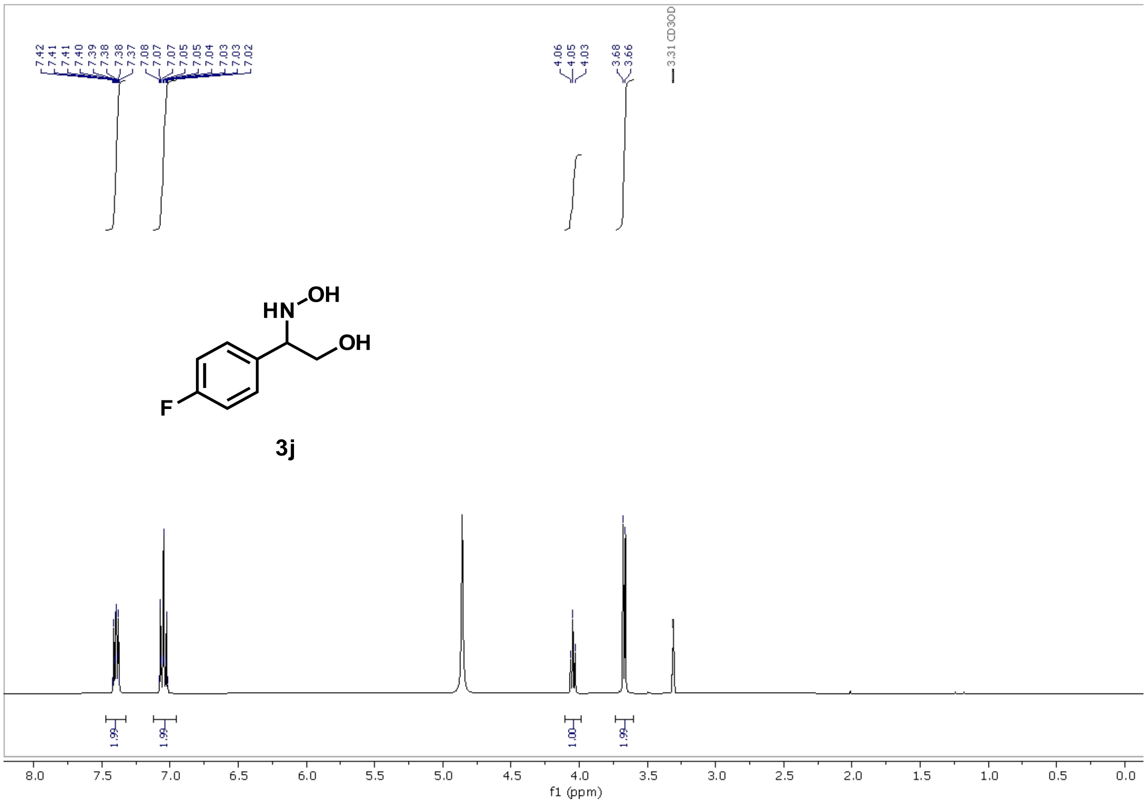
7.42
7.41
7.40
7.39
7.38
7.38
7.37
7.08
7.07
7.07
7.05
7.05
7.04
7.03
7.03
7.02
4.06
4.05
4.03
3.68
3.66
3.31 CD3OD
HN
OH
OH
F
3j
1.99
1.99
1.00
1.99
8.0
7.5
7.0
6.5
6.0
5.5
5.0
4.5
4.0
3.5
3.0
2.5
2.0
1.5
1.0
0.5
0.0
f1 (ppm)

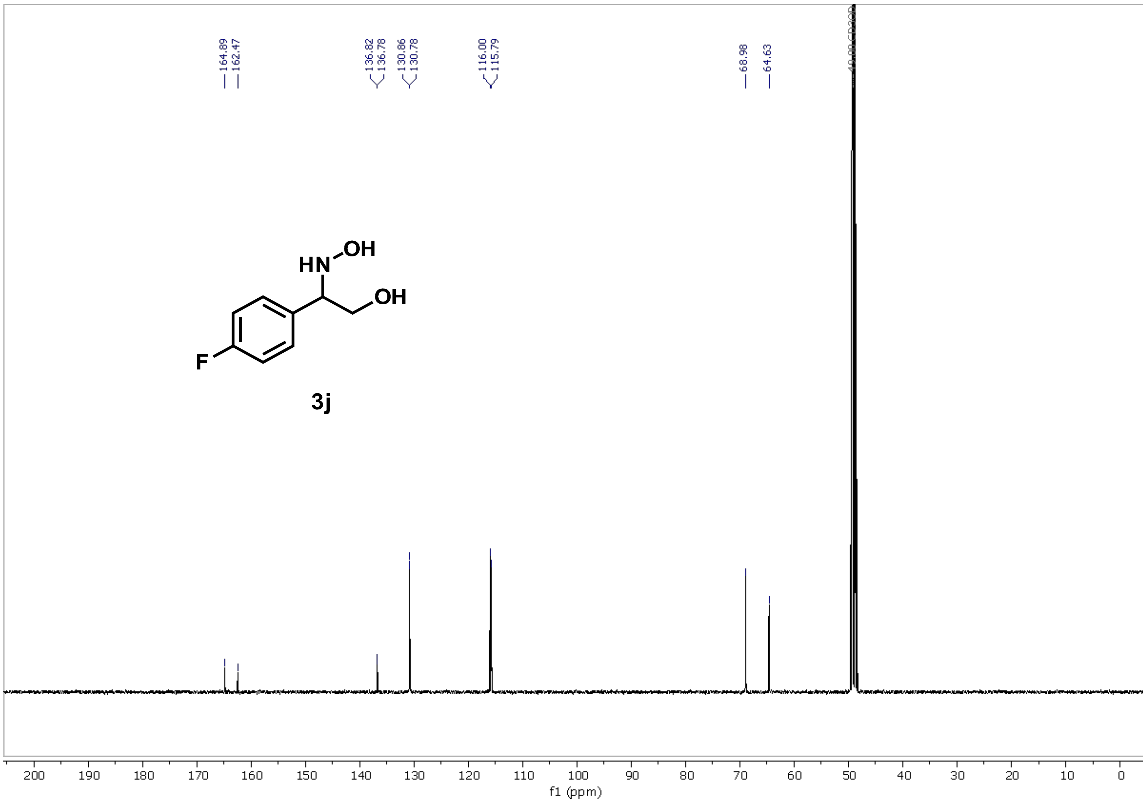
164.89
162.47
136.82
136.78
130.86
130.78
116.00
115.79
68.98
64.63
HN
OH
OH
F
3j
200
190
180
170
160
150
140
130
120
110
100
90
80
70
60
50
40
30
20
10
0
f1 (ppm)

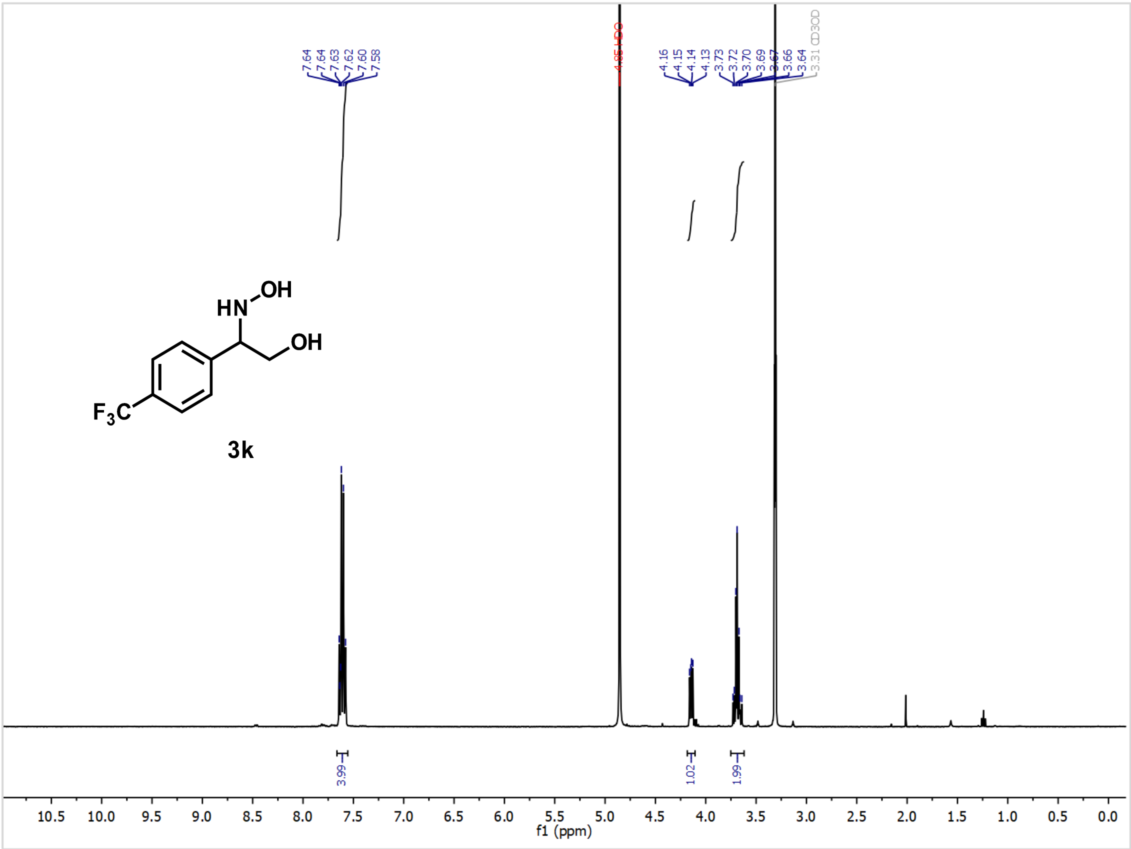
7.64
7.64
7.63
7.62
7.60
7.58
4.85 HDO
4.16
4.15
4.14
4.13
3.73
3.72
3.70
3.69
3.67
3.66
3.64
3.31 CD3OD
HN
OH
OH
F3C
3k
3.99
1.02
1.99
f1 (ppm)

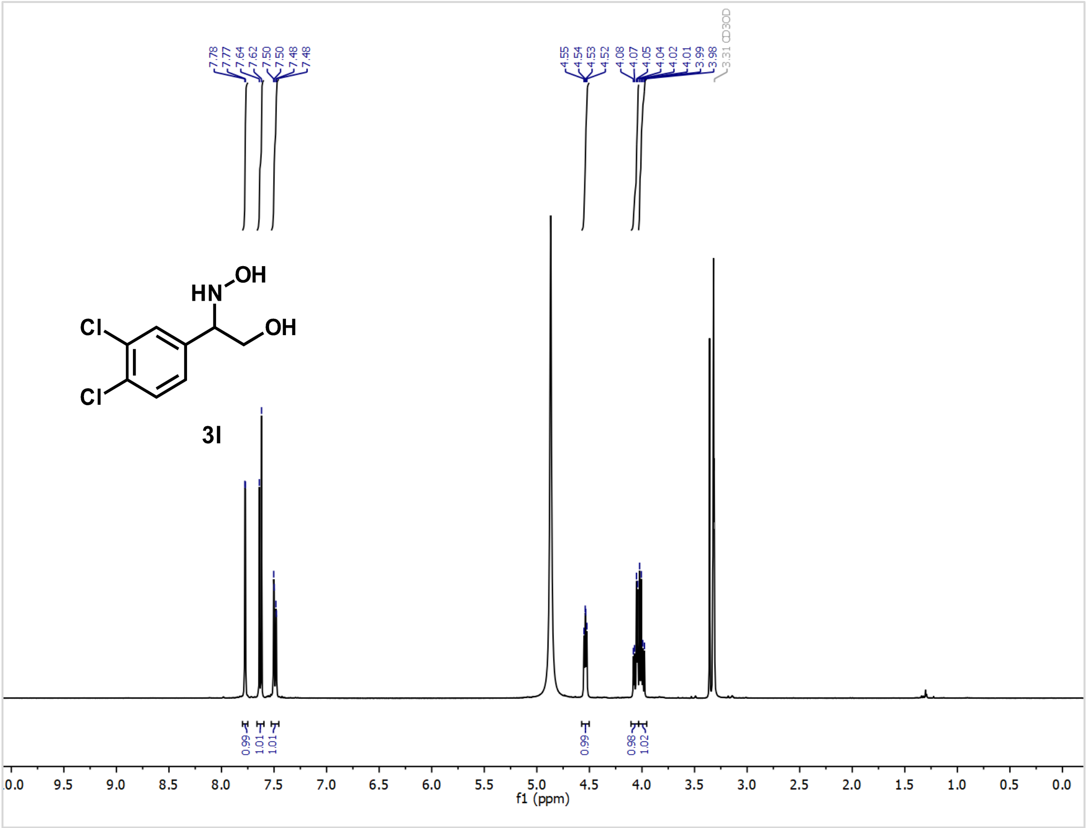
7.78
7.77
7.64
7.62
7.50
7.50
7.48
7.48
4.55
4.54
4.53
4.52
4.08
4.07
4.05
4.04
4.02
4.01
3.99
3.98
3.31 CD3OD
HN
OH
Cl
OH
Cl
3l
0.99
1.01
1.01
0.99
0.98
1.02
10.0 9.5 9.0 8.5 8.0 7.5 7.0 6.5 6.0 5.5 5.0 4.5 4.0 3.5 3.0 2.5 2.0 1.5 1.0 0.5 0.0
f1 (ppm)

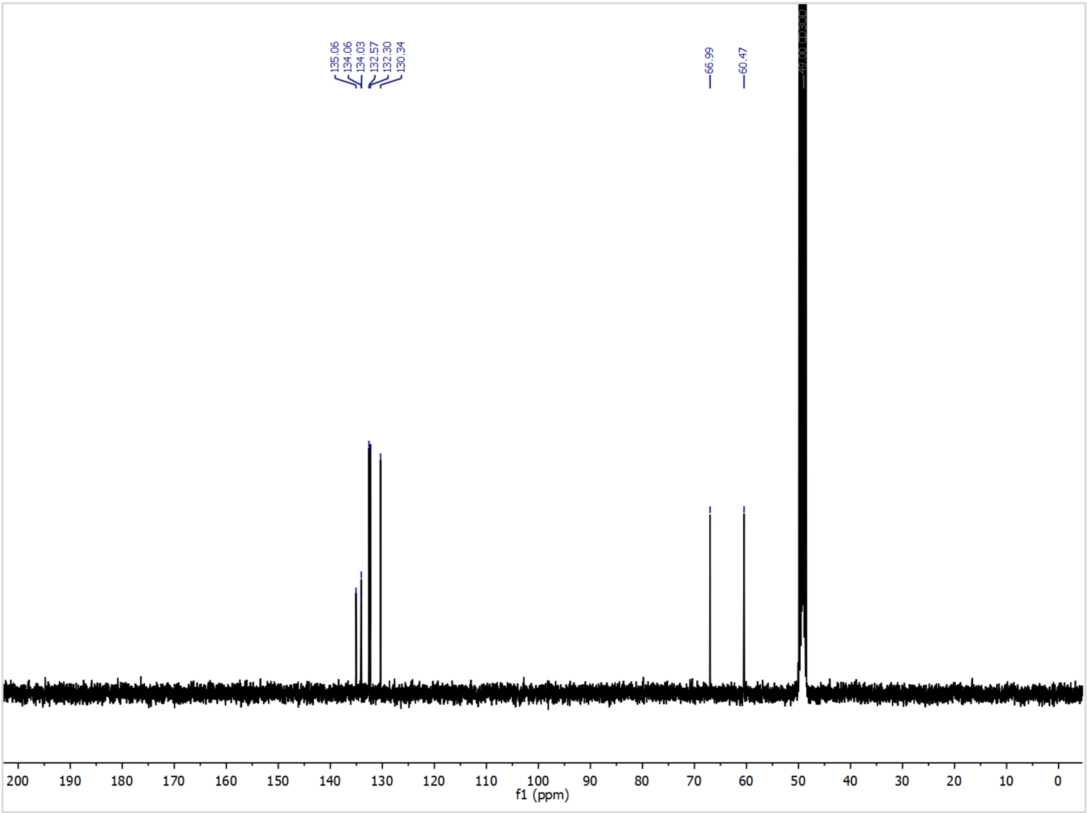
135.06
134.06
134.03
132.57
132.30
130.34
66.99
60.47
49.00 CD3OD
200 190 180 170 160 150 140 130 120 110 100 90 80 70 60 50 40 30 20 10 0
f1 (ppm)

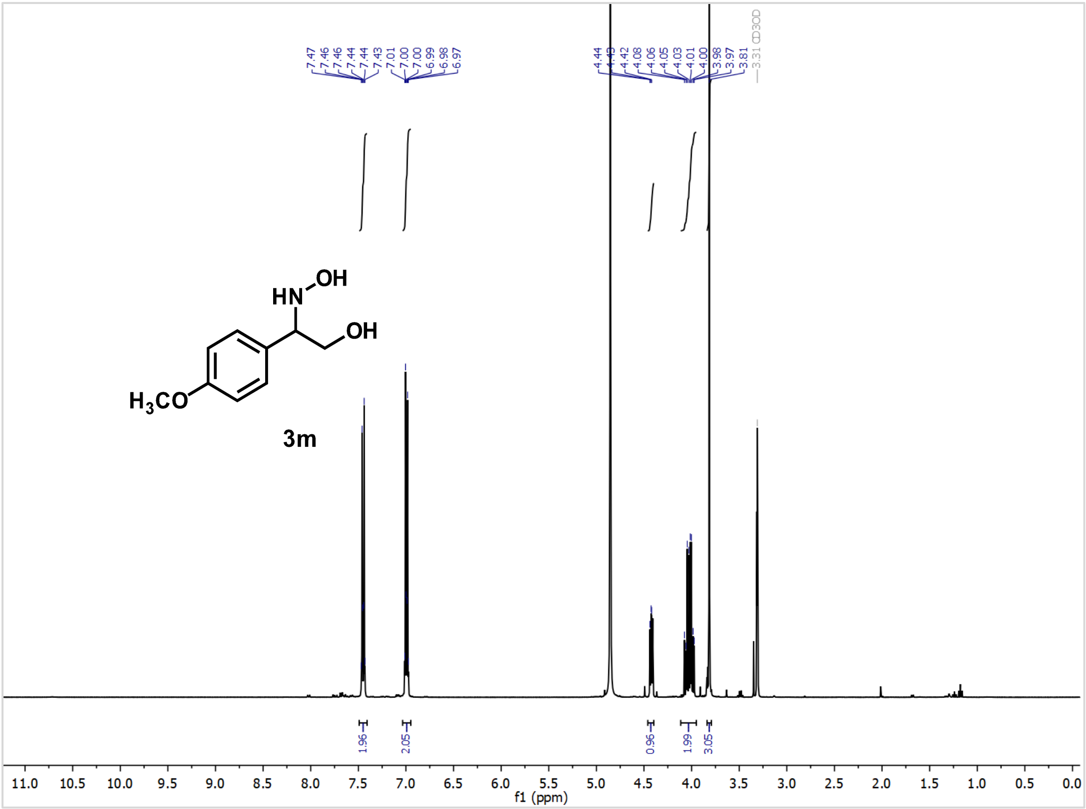
7.47
7.46
7.46
7.44
7.44
7.43
7.01
7.00
7.00
6.99
6.98
6.97
4.44
4.42
4.08
4.06
4.05
4.03
4.01
4.00
3.98
3.97
3.81
3.31 CD3OD
HN
OH
OH
H3CO
3m
1.96
2.05
0.96
1.99
3.05
11.0 10.5 10.0 9.5 9.0 8.5 8.0 7.5 7.0 6.5 6.0 5.5 5.0 4.5 4.0 3.5 3.0 2.5 2.0 1.5 1.0 0.5 0.0
f1 (ppm)

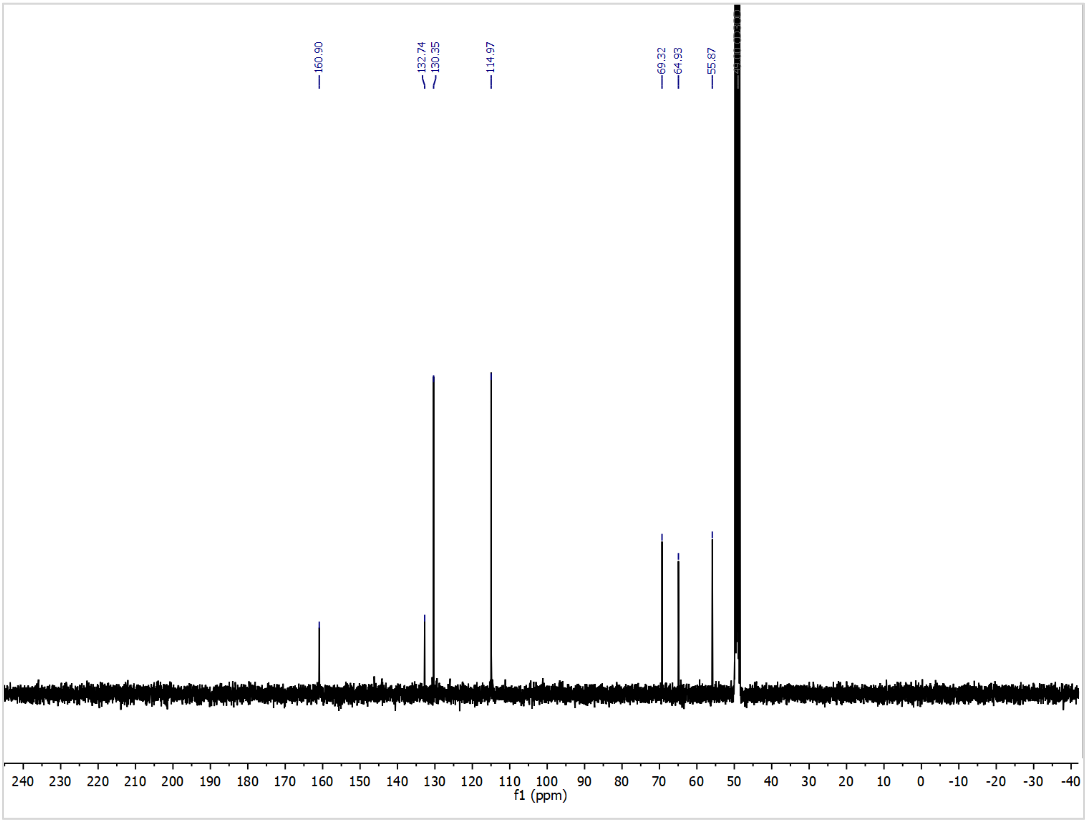
160.90
132.74
130.35
114.97
69.32
64.93
55.87
240 230 220 210 200 190 180 170 160 150 140 130 120 110 100 90 80 70 60 50 40 30 20 10 0 -10 -20 -30 -40
f1 (ppm)

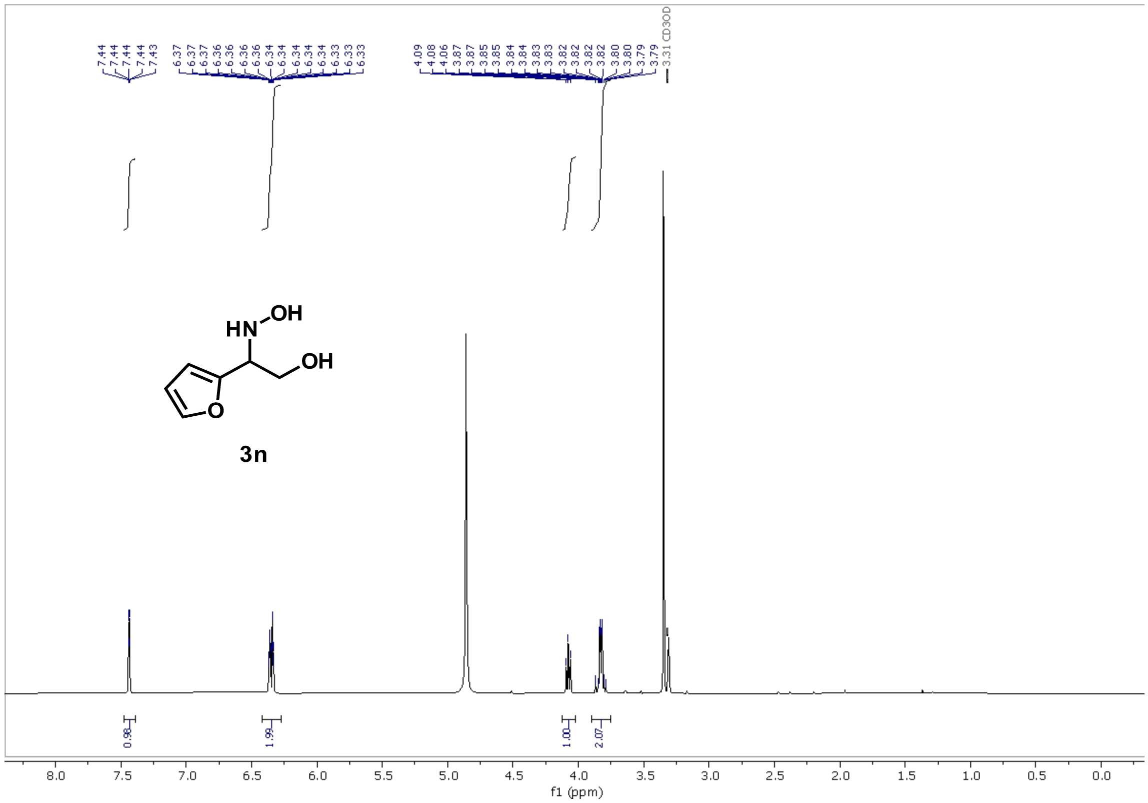
7.44
7.44
7.44
7.44
7.43
6.37
6.37
6.37
6.36
6.36
6.36
6.34
6.34
6.34
6.34
6.34
6.33
6.33
6.33
4.09
4.08
4.06
3.87
3.87
3.85
3.85
3.84
3.84
3.83
3.83
3.82
3.82
3.82
3.82
3.80
3.80
3.79
3.79
3.31 CD3OD
HN
OH
OH
O
3n
0.98
1.99
1.00
2.07
8.0
7.5
7.0
6.5
6.0
5.5
5.0
4.5
4.0
3.5
3.0
2.5
2.0
1.5
1.0
0.5
0.0
f1 (ppm)

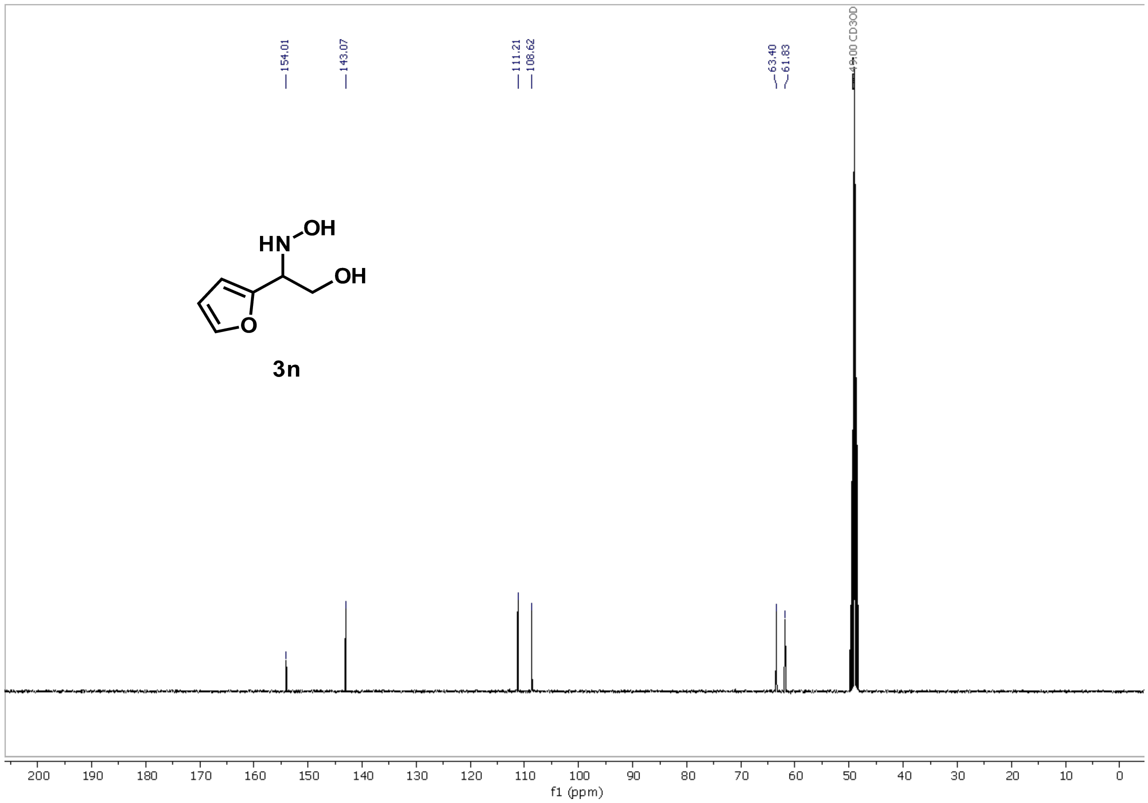
154.01
143.07
111.21
108.62
63.40
61.83
49.00 CD3OD
HN
OH
OH
O
3n
200
190
180
170
160
150
140
130
120
110
100
90
80
70
60
50
40
30
20
10
0
f1 (ppm)

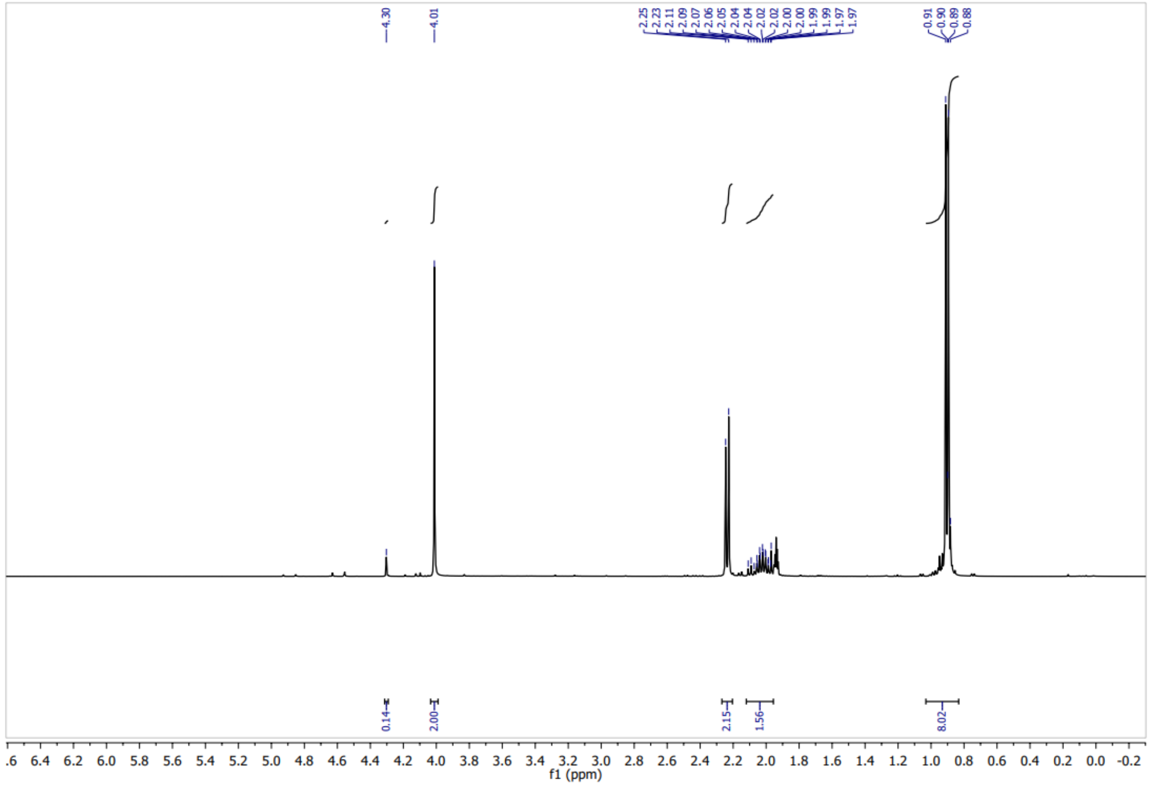
4.30
4.01
2.25 2.23 2.11 2.09 2.07 2.06 2.05 2.04 2.04 2.02 2.02 2.00 2.00 1.99 1.99 1.97 1.97
0.91 0.90 0.89 0.88
0.14
2.00
2.15
1.56
8.02
f1 (ppm)

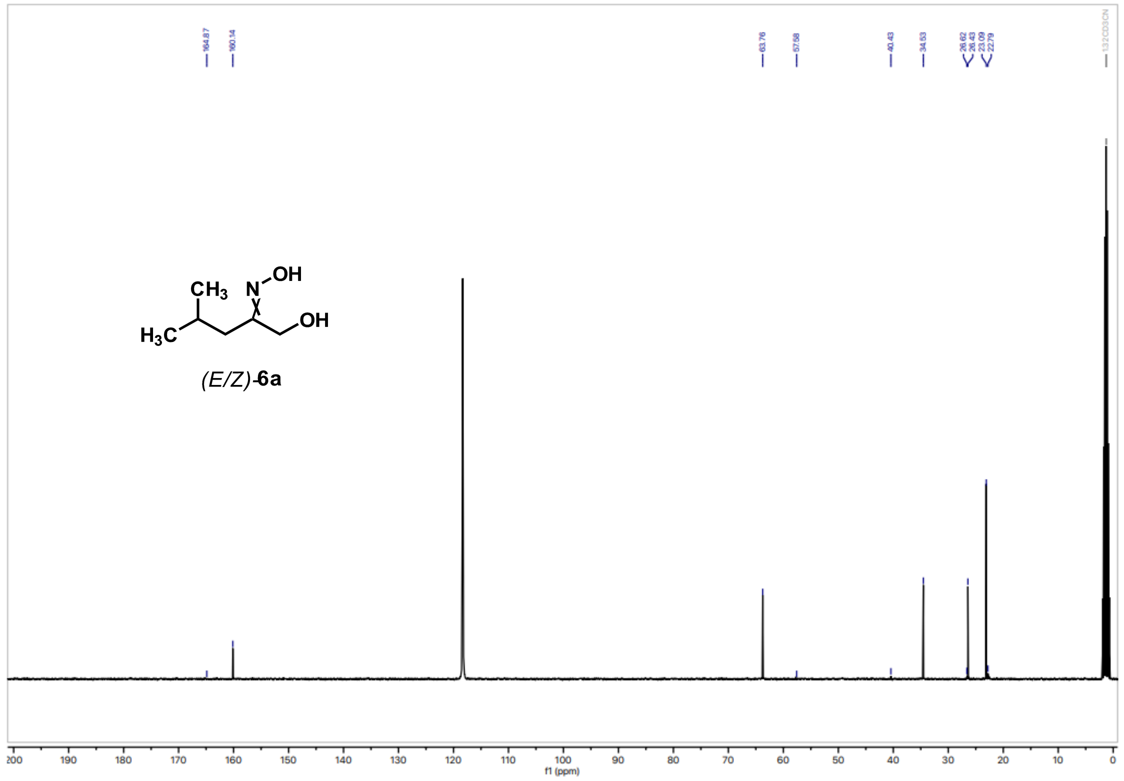
164.87
160.14
63.76
57.58
40.43
34.53
26.62 26.43 23.09 22.79
CH3 N OH
H3C OH
(E/Z)-6a
f1 (ppm)

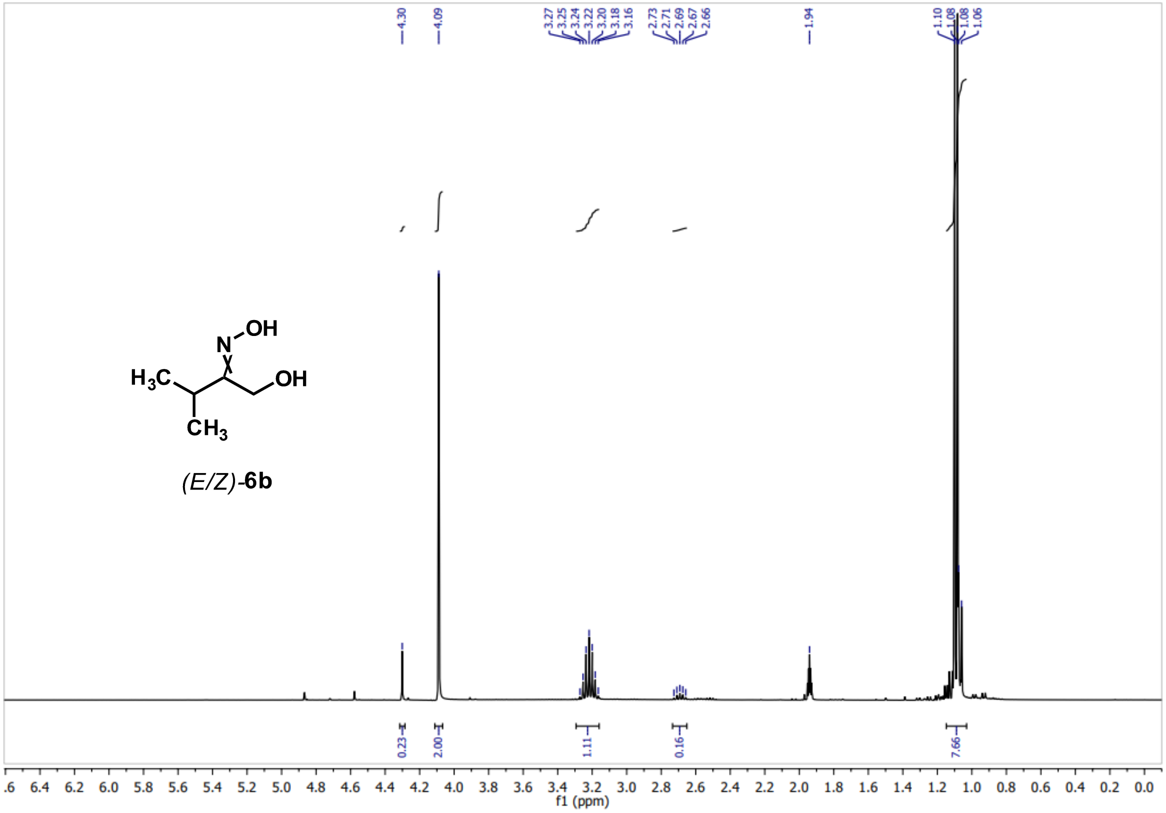
4.30
4.09
3.27 3.25 3.24 3.22 3.20 3.18 3.16
2.73 2.71 2.69 2.67 2.66
1.94
1.10 1.08 1.08 1.06
(E/Z)-6b
0.23
2.00
1.11
0.16
7.66
f1 (ppm)

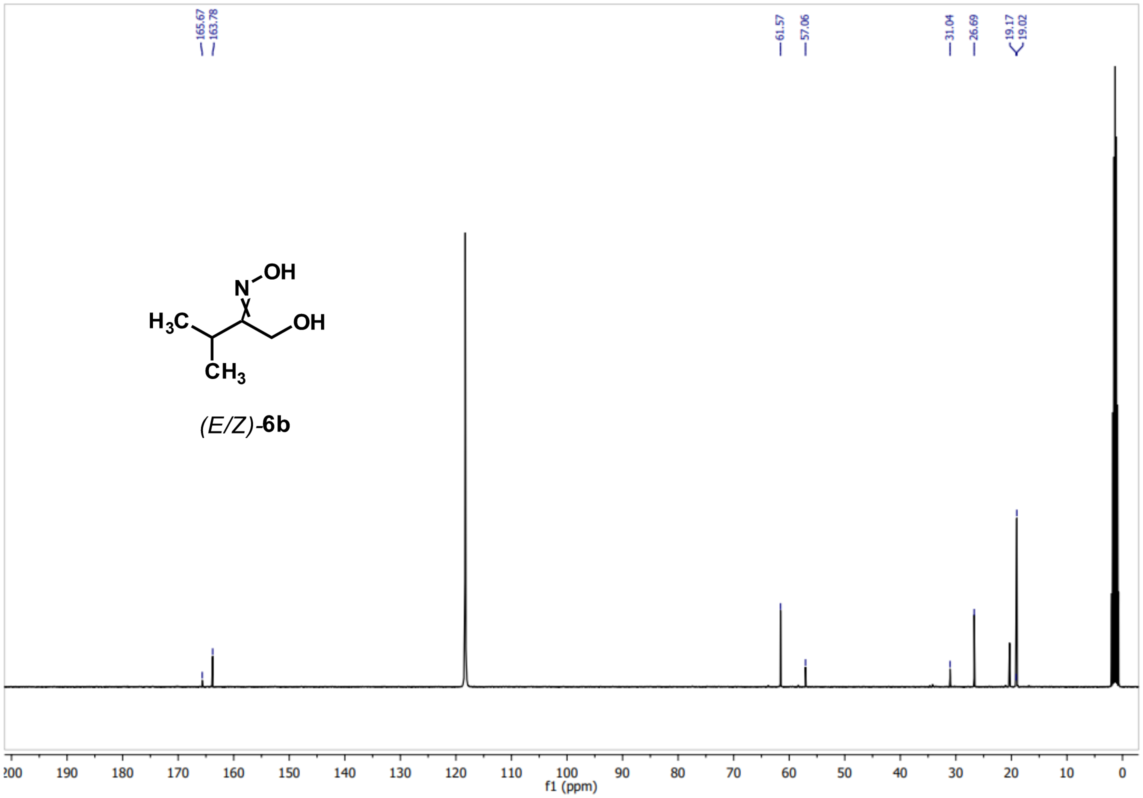
165.67
163.78
61.57
57.06
31.04
26.69
19.17
19.02
(E/Z)-6b
f1 (ppm)

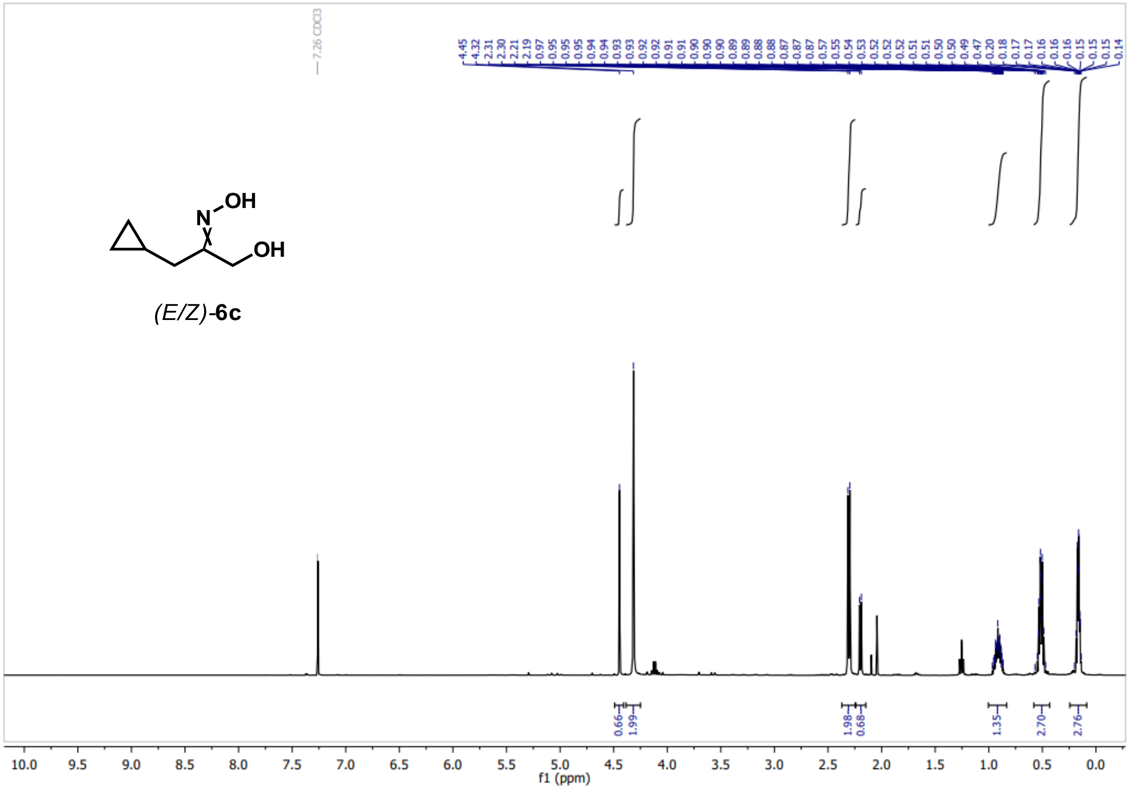

(E/Z)-**6c**

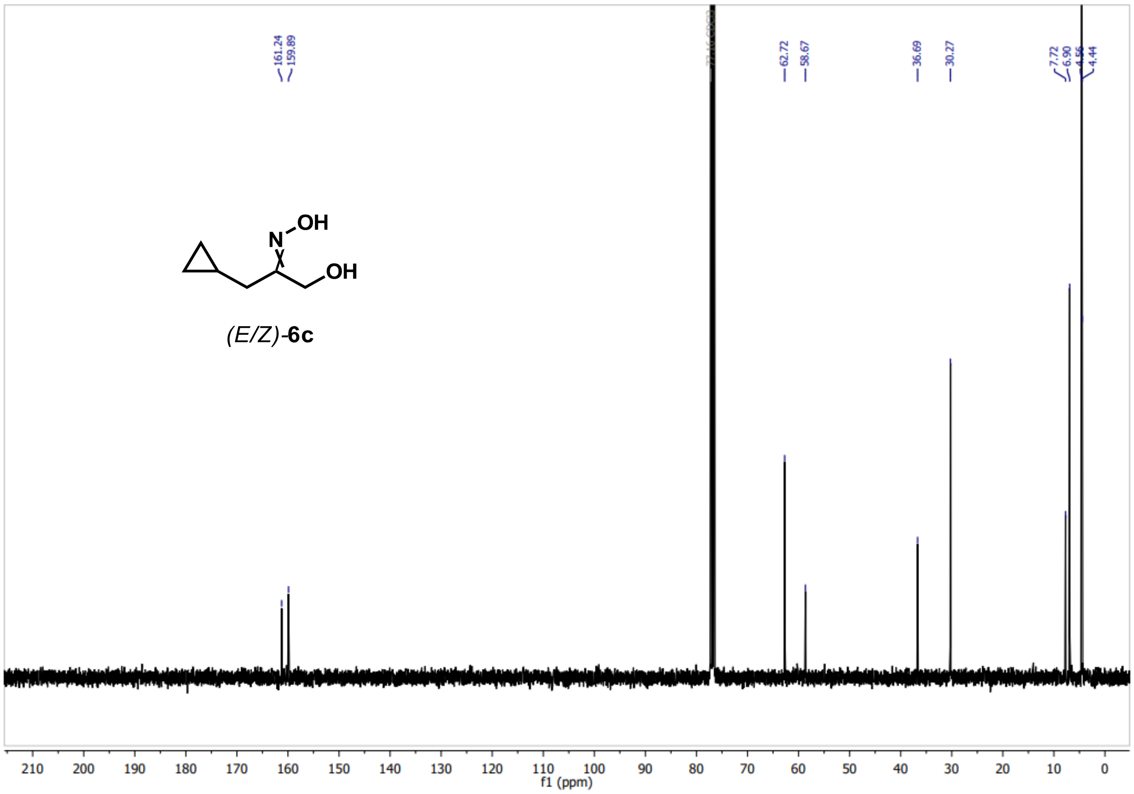

(E/Z)-**6c**

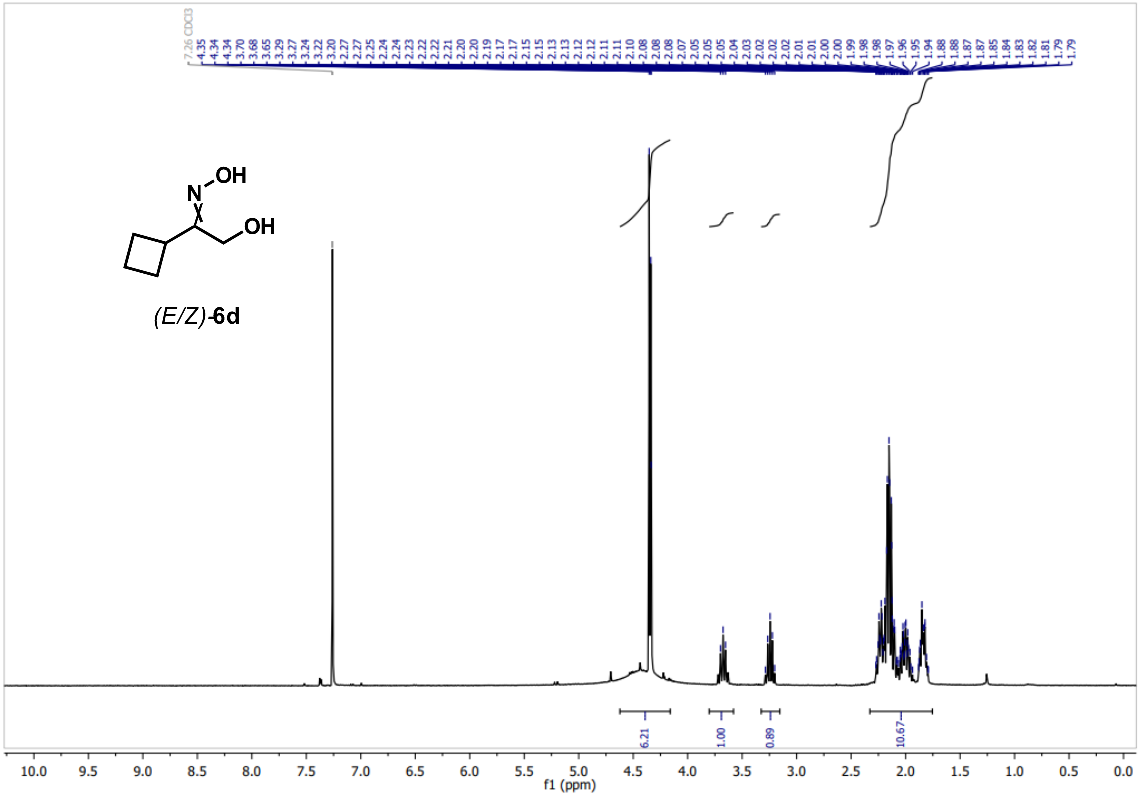
(E/Z)-6d
N
OH
OH
6.21
1.00
0.89
10.67
f1 (ppm)

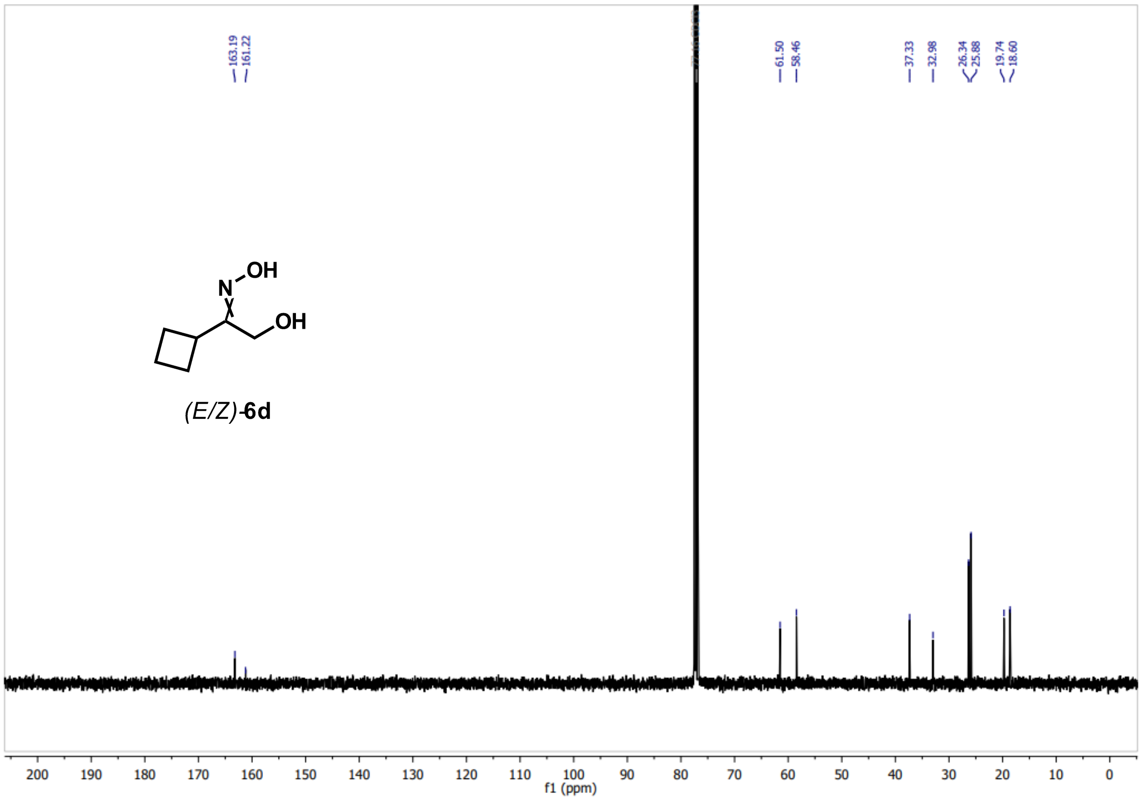
163.19
161.22
61.50
58.46
37.33
32.98
26.34
25.88
19.74
18.60
(E/Z)-6d
N
OH
OH
f1 (ppm)

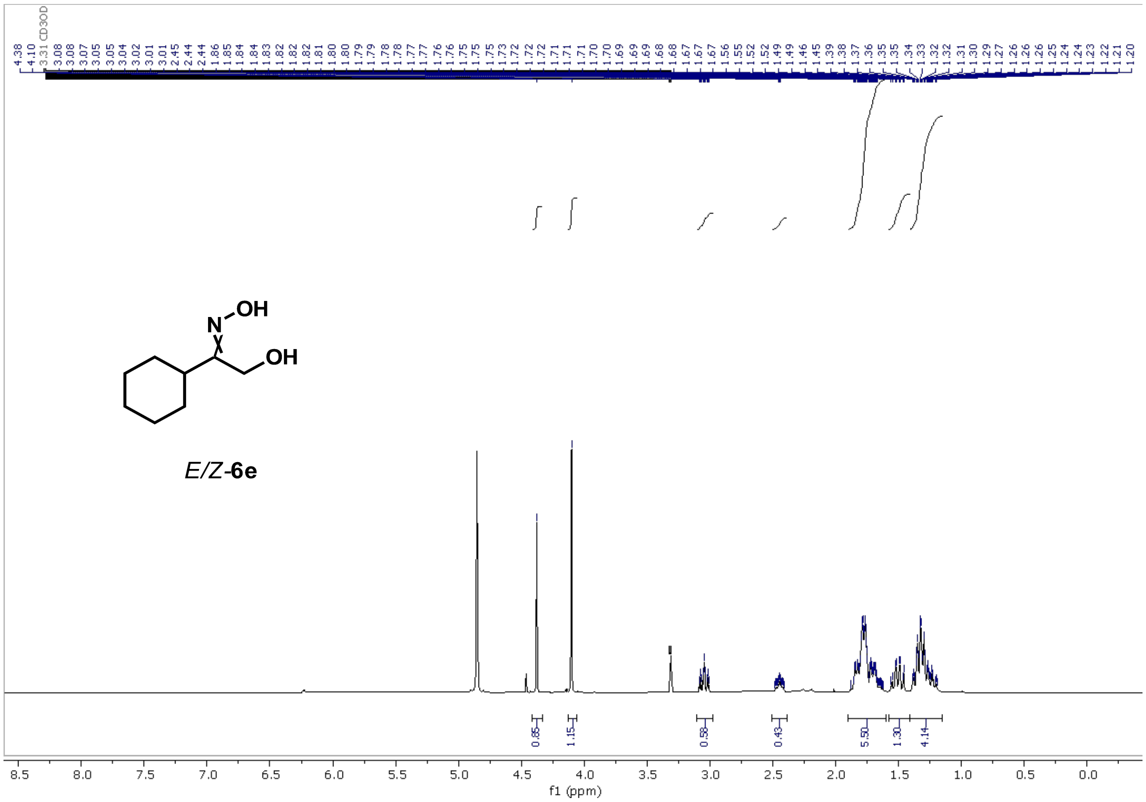

*E*/*Z*-**6e**

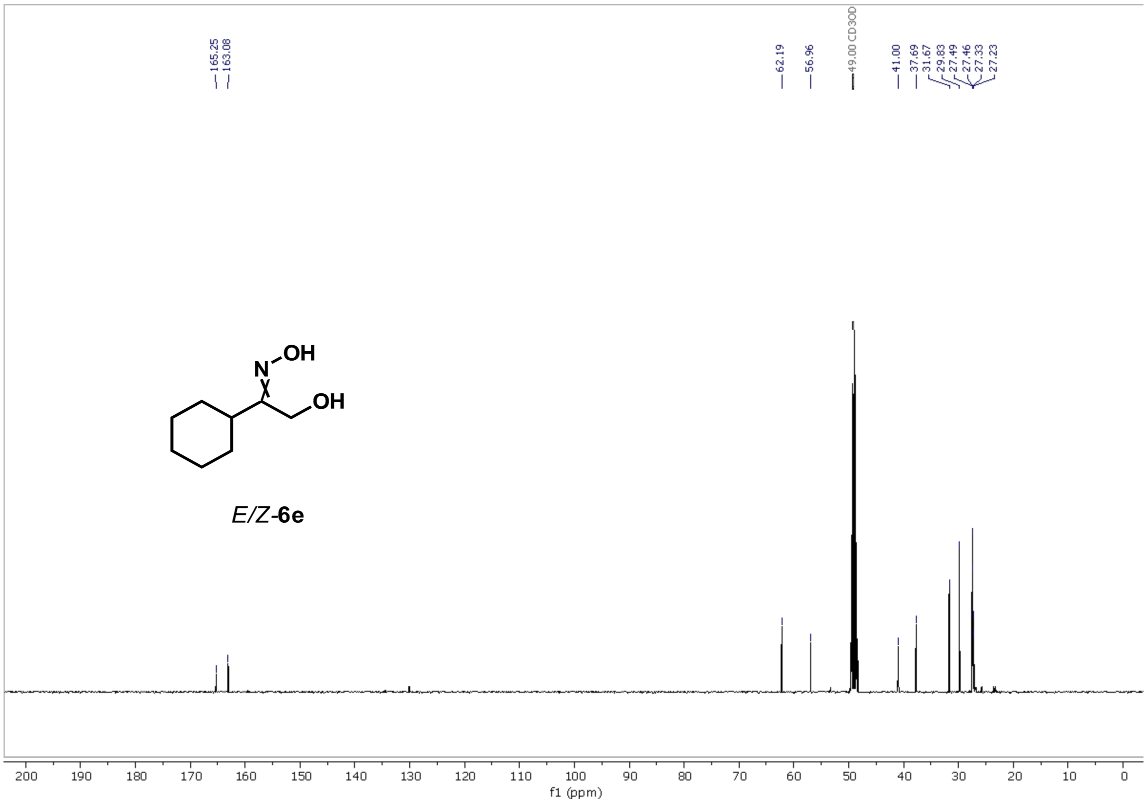

*E*/*Z*-**6e**

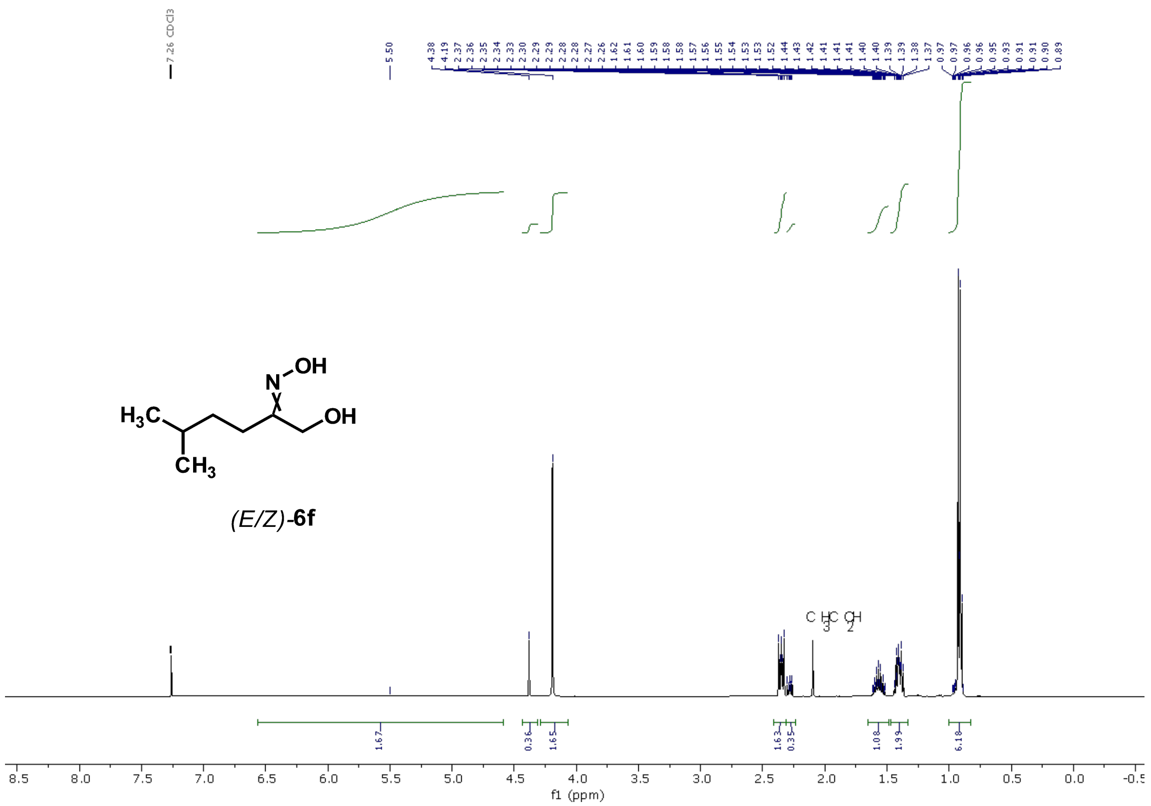
7.26 CDCl3
5.50
(E/Z)-6f
f1 (ppm)

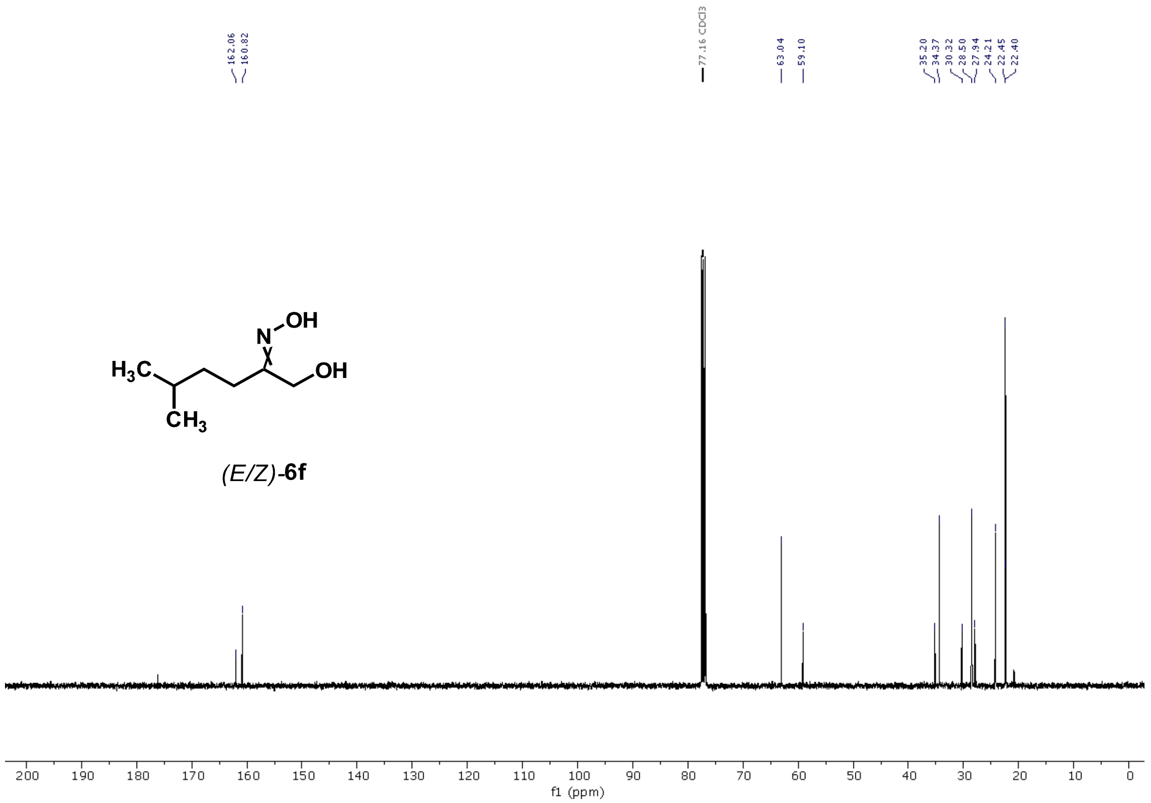
162.06
160.82
77.16 CDCl3
63.04
59.10
35.20
34.37
30.32
28.50
27.94
24.21
22.45
22.40
(E/Z)-6f
f1 (ppm)

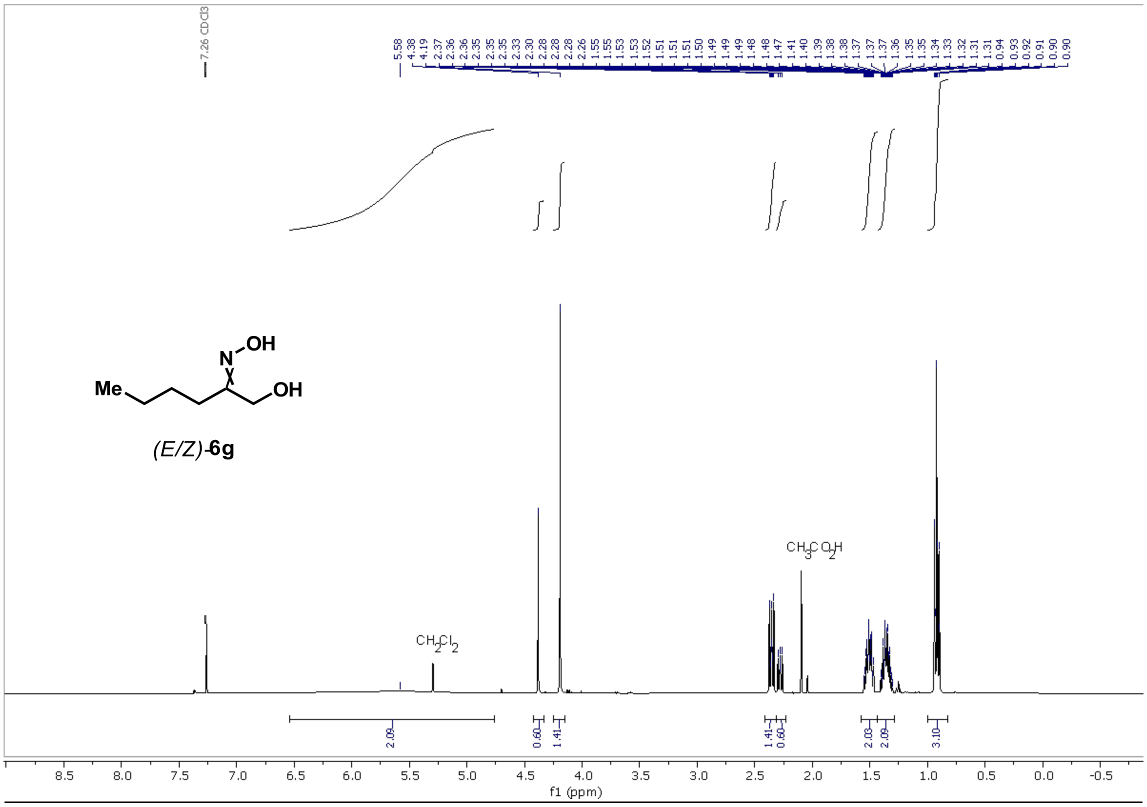
7.26 CDCl3
5.58
4.38
4.19
2.37
2.36
2.36
2.35
2.35
2.35
2.33
2.30
2.28
2.28
2.28
2.26
1.55
1.55
1.53
1.53
1.52
1.51
1.51
1.51
1.50
1.49
1.49
1.49
1.48
1.48
1.47
1.41
1.40
1.39
1.38
1.38
1.38
1.37
1.37
1.37
1.36
1.35
1.35
1.34
1.33
1.32
1.31
1.31
0.94
0.93
0.93
0.92
0.91
0.90
0.90
N
OH
Me
OH
(E/Z)-6g
CH3CO2H
CH2Cl2
2.09
0.60
1.41
1.41
0.60
2.03
2.09
3.10
8.5
8.0
7.5
7.0
6.5
6.0
5.5
5.0
4.5
4.0
3.5
3.0
2.5
2.0
1.5
1.0
0.5
0.0
-0.5
f1 (ppm)

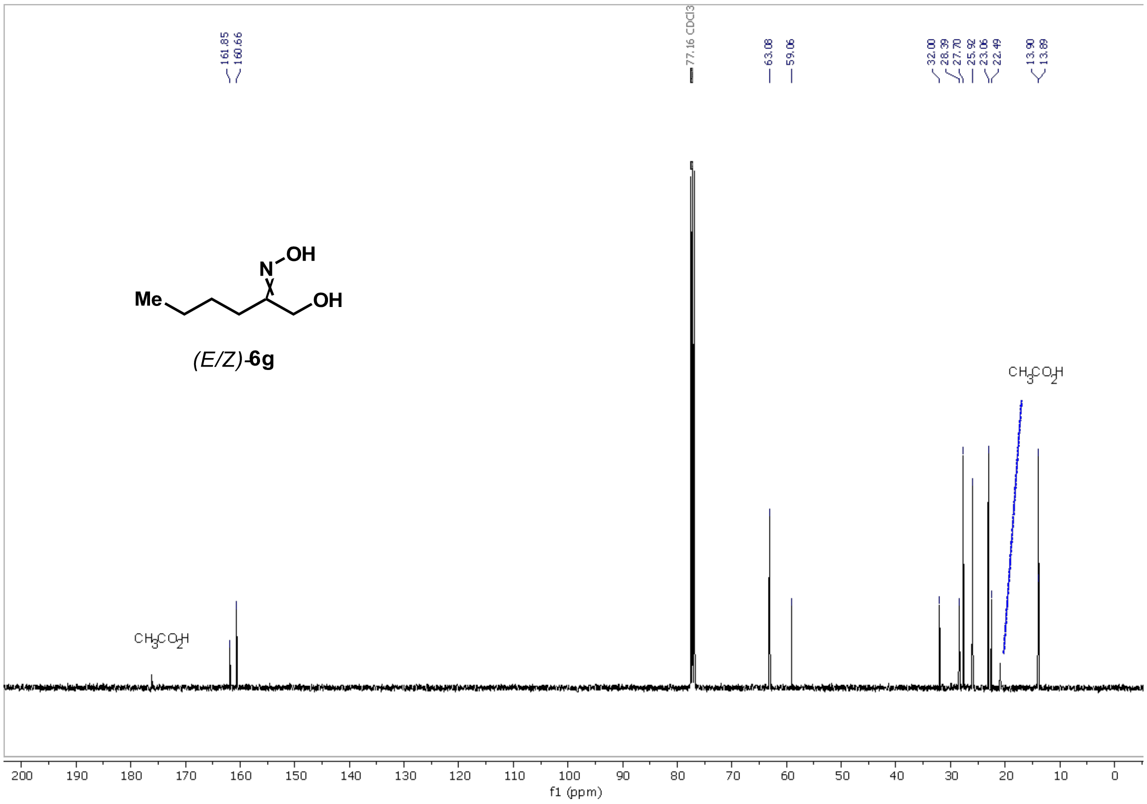
161.85
160.66
77.16 CDCl3
63.08
59.06
32.00
28.39
27.70
25.92
23.06
22.49
13.90
13.89
N
OH
Me
OH
(E/Z)-6g
CH3CO2H
CH3CO2H
200
190
180
170
160
150
140
130
120
110
100
90
80
70
60
50
40
30
20
10
0
f1 (ppm)

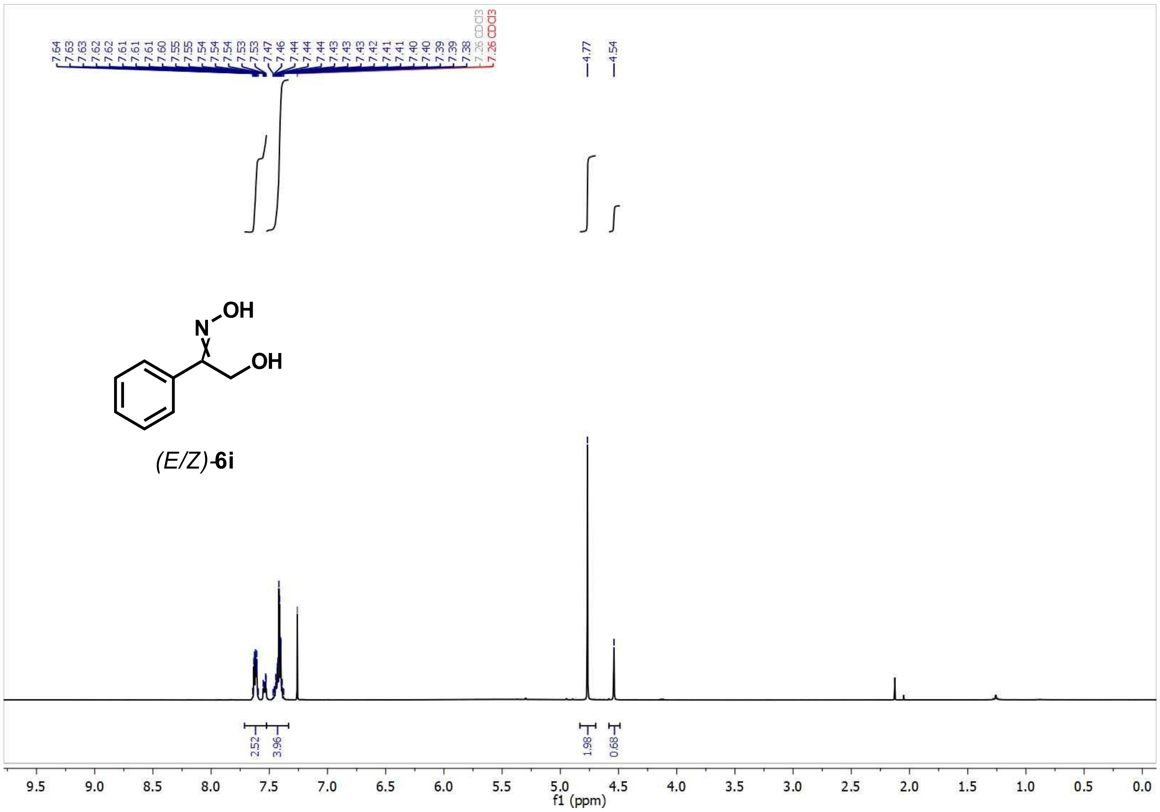
7.64
7.63
7.63
7.62
7.62
7.61
7.61
7.61
7.60
7.55
7.55
7.54
7.54
7.54
7.53
7.53
7.47
7.46
7.44
7.44
7.44
7.43
7.43
7.43
7.42
7.41
7.40
7.40
7.39
7.39
7.38
7.26 CDCl3
7.26 CDCl3
4.77
4.54
N
OH
OH
(E/Z)-6i
2.52
3.96
1.98
0.68
9.5 9.0 8.5 8.0 7.5 7.0 6.5 6.0 5.5 5.0 4.5 4.0 3.5 3.0 2.5 2.0 1.5 1.0 0.5 0.0
f1 (ppm)

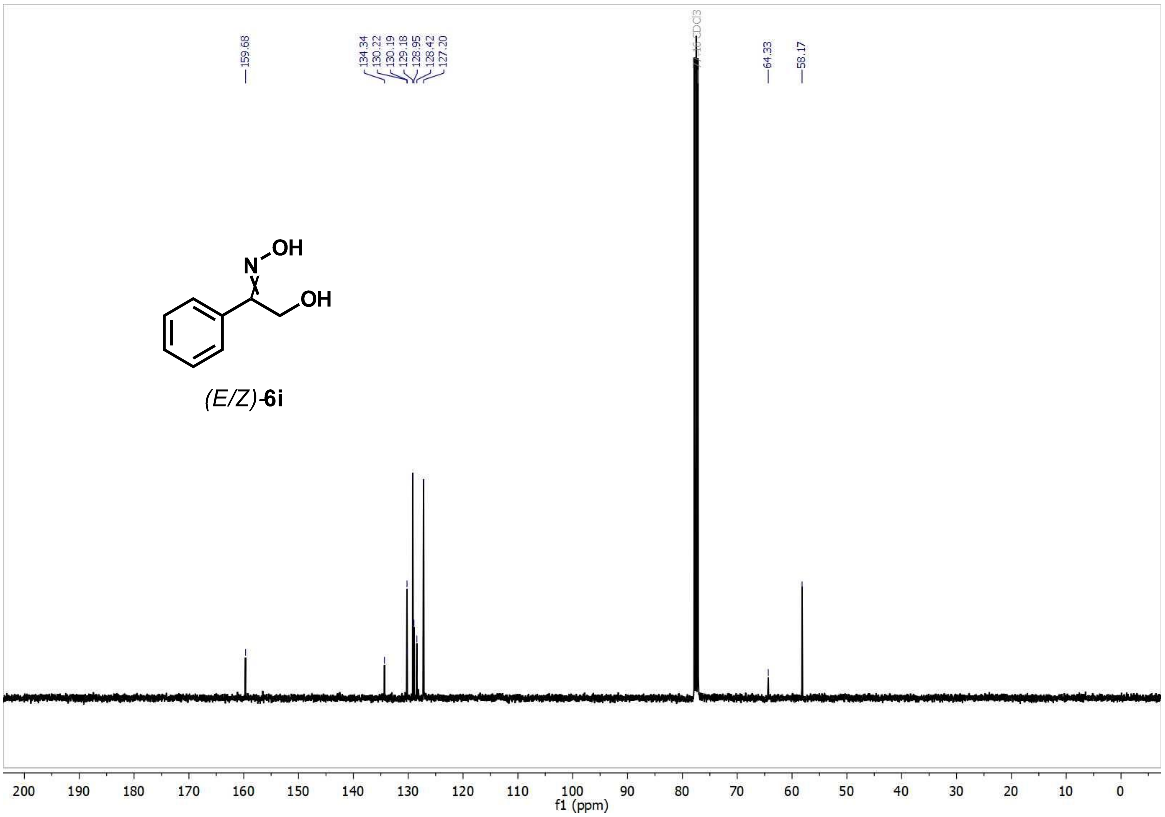
159.68
134.34
130.22
130.19
129.18
128.95
128.42
127.20
CDCl3
64.33
58.17
N
OH
OH
(E/Z)-6i
200 190 180 170 160 150 140 130 120 110 100 90 80 70 60 50 40 30 20 10 0
f1 (ppm)

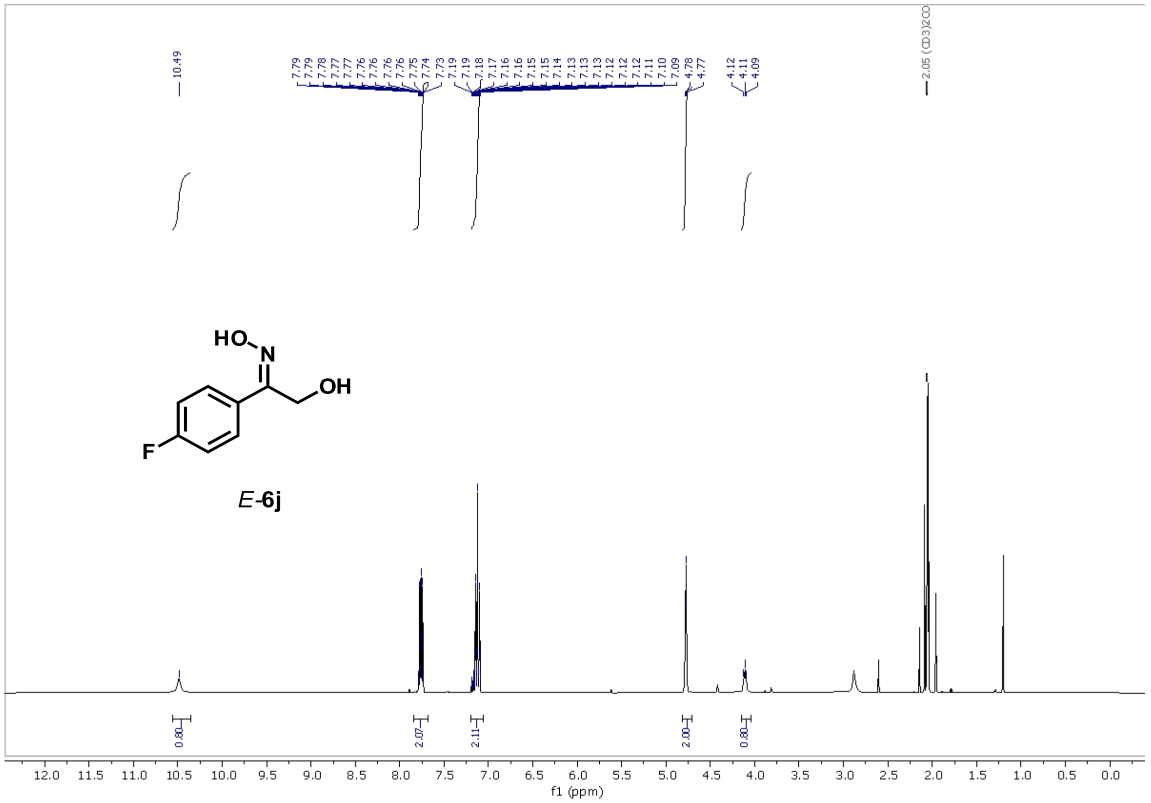

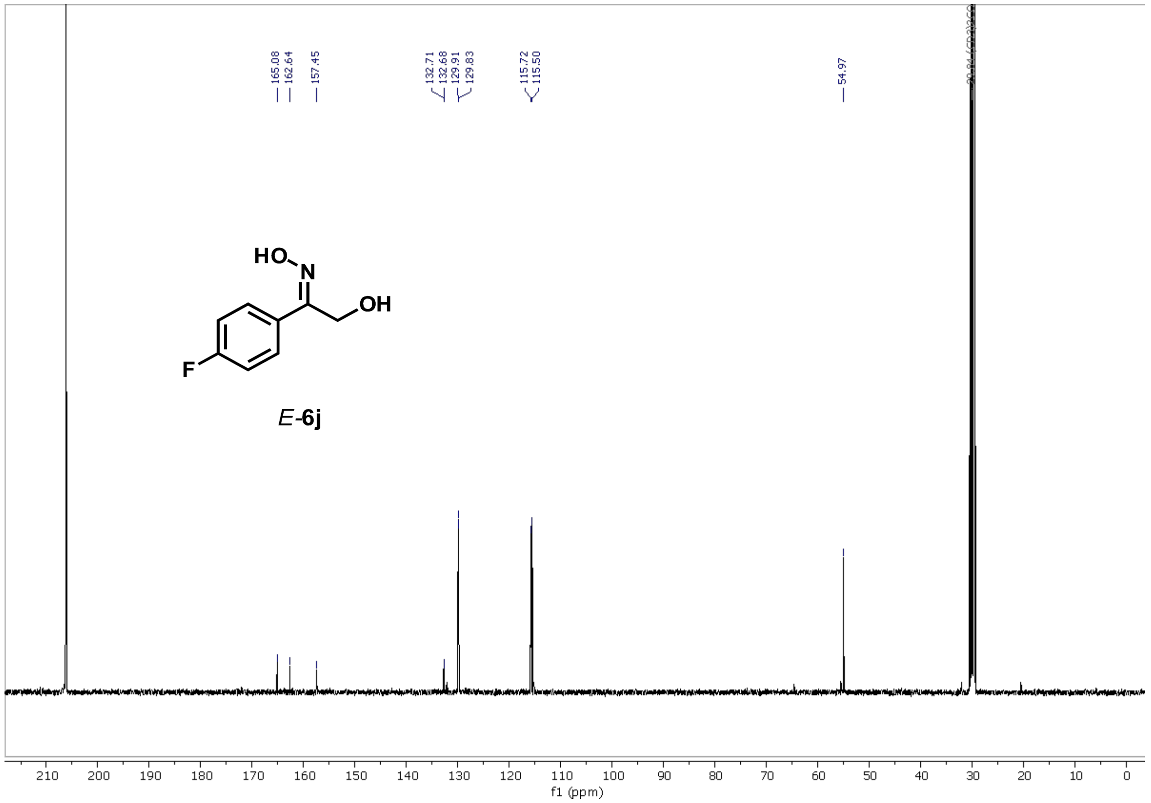

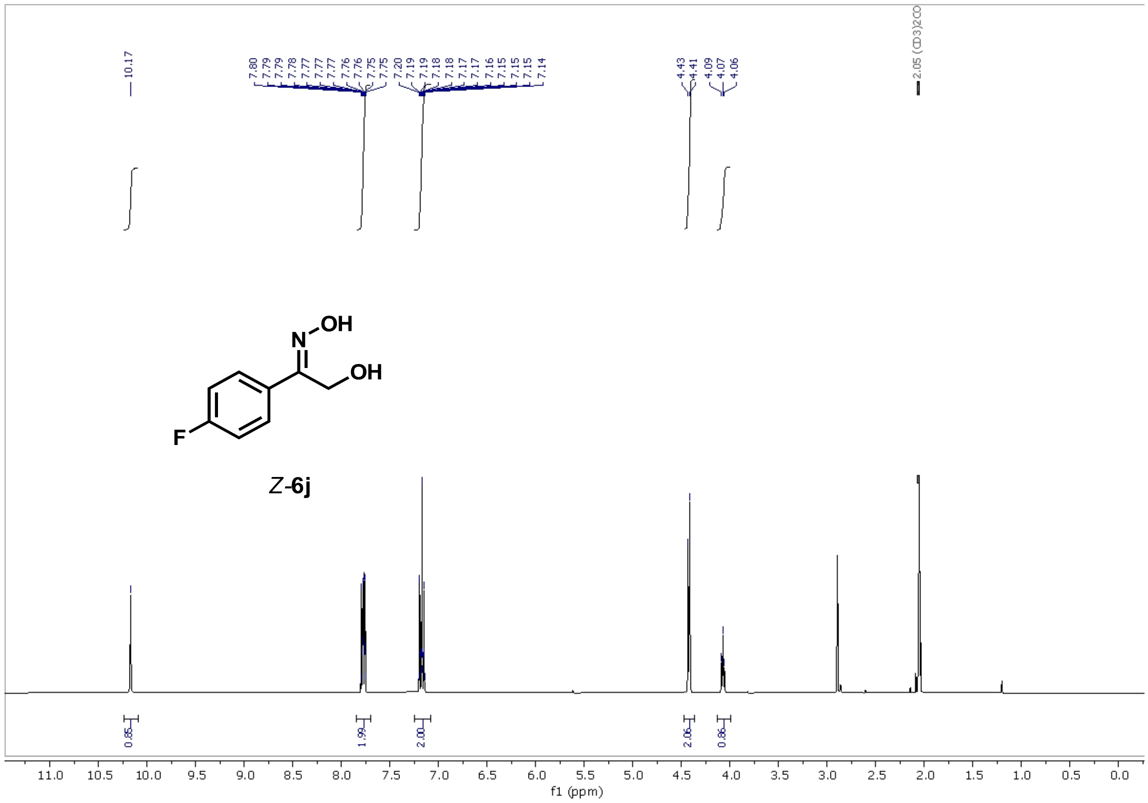

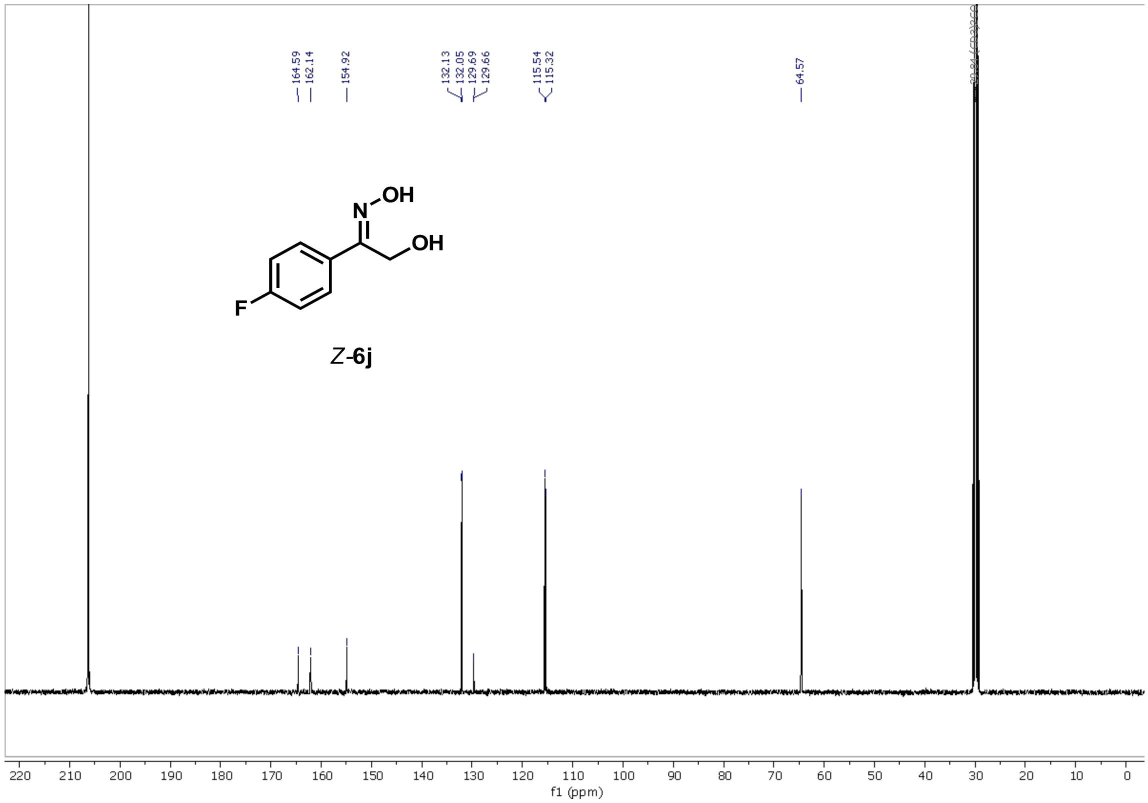

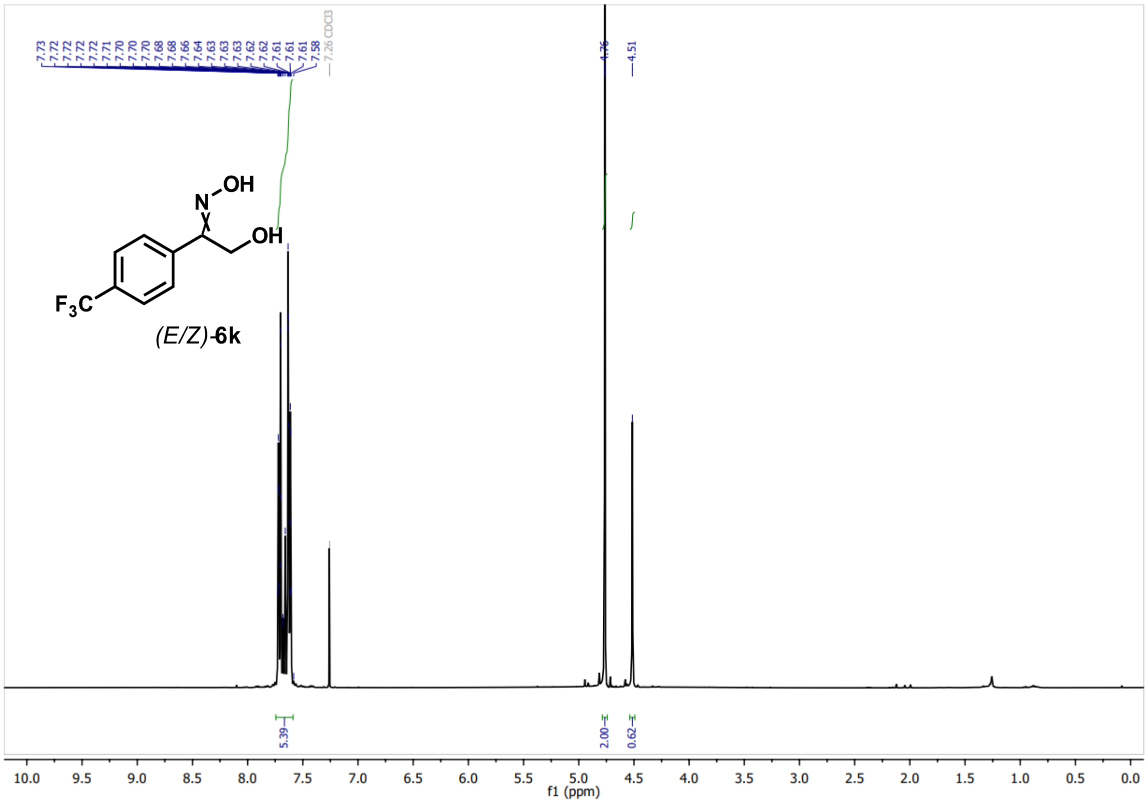

7.73
7.72
7.72
7.72
7.72
7.71
7.70
7.70
7.70
7.68
7.68
7.66
7.64
7.63
7.63
7.63
7.62
7.62
7.61
7.61
7.61
7.58
7.26 CDCl3
4.76
4.51
(E/Z)-6k
5.39
2.00
0.62
f1 (ppm)


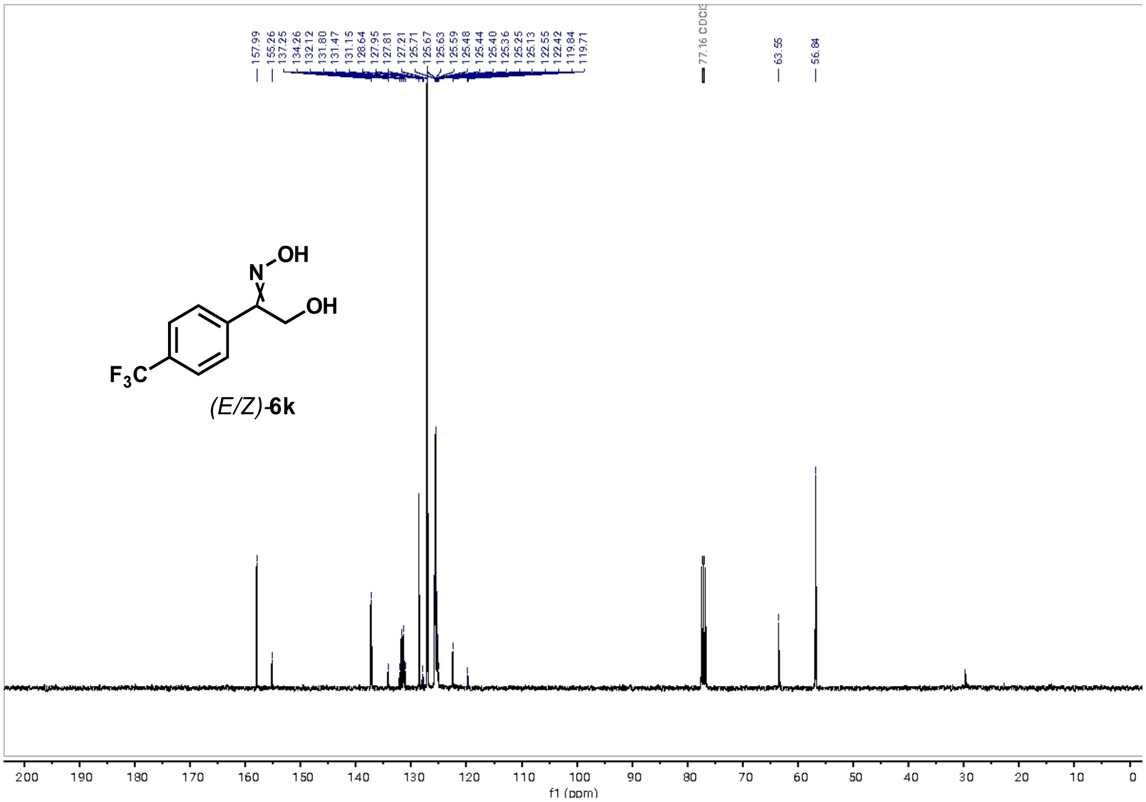

157.99
155.26
137.25
134.26
132.12
131.80
131.47
131.15
128.64
127.95
127.81
127.21
125.71
125.67
125.63
125.59
125.48
125.44
125.40
125.36
125.25
125.13
122.55
122.42
119.84
119.71
77.16 CDCl3
63.55
56.84
(E/Z)-6k
f1 (ppm)

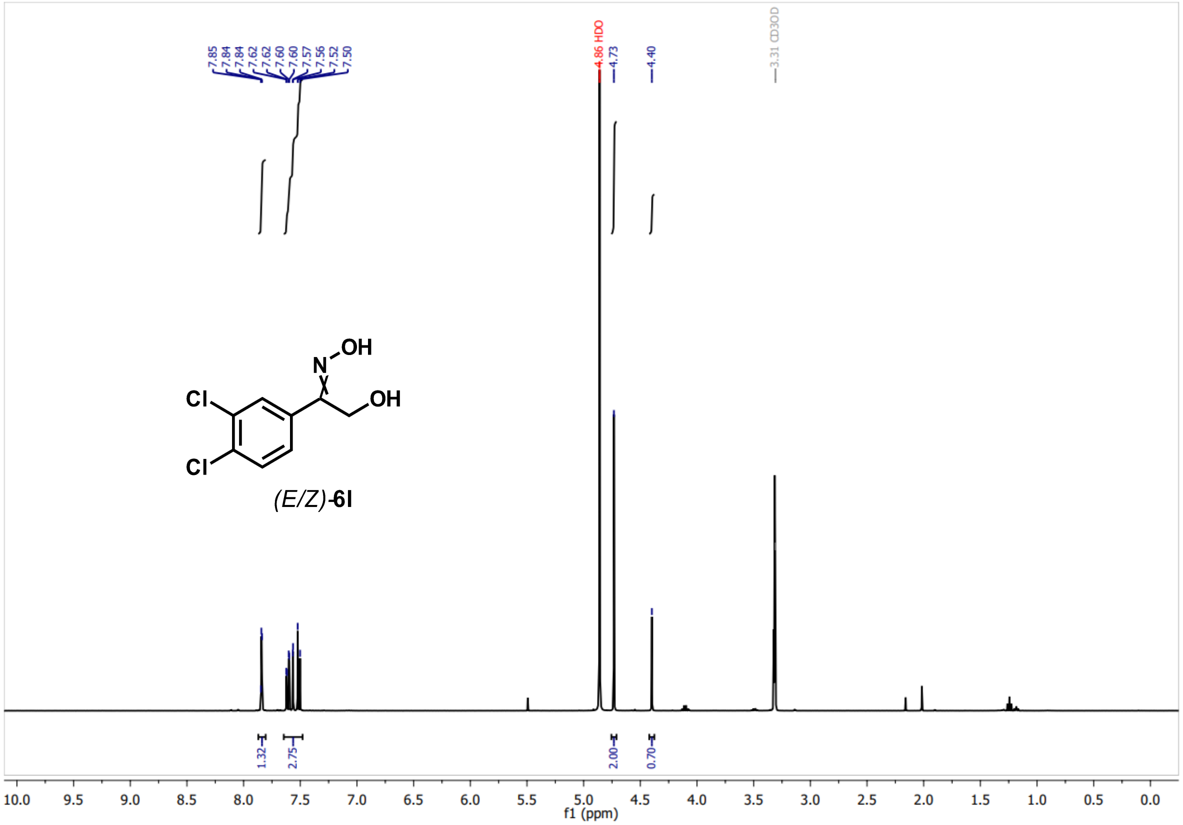
7.85
7.84
7.84
7.62
7.62
7.60
7.60
7.57
7.56
7.52
7.50
4.86 HDO
4.73
4.40
3.31 CD3OD
Cl
Cl
N
OH
OH
(E/Z)-6l
1.32
2.75
2.00
0.70
f1 (ppm)

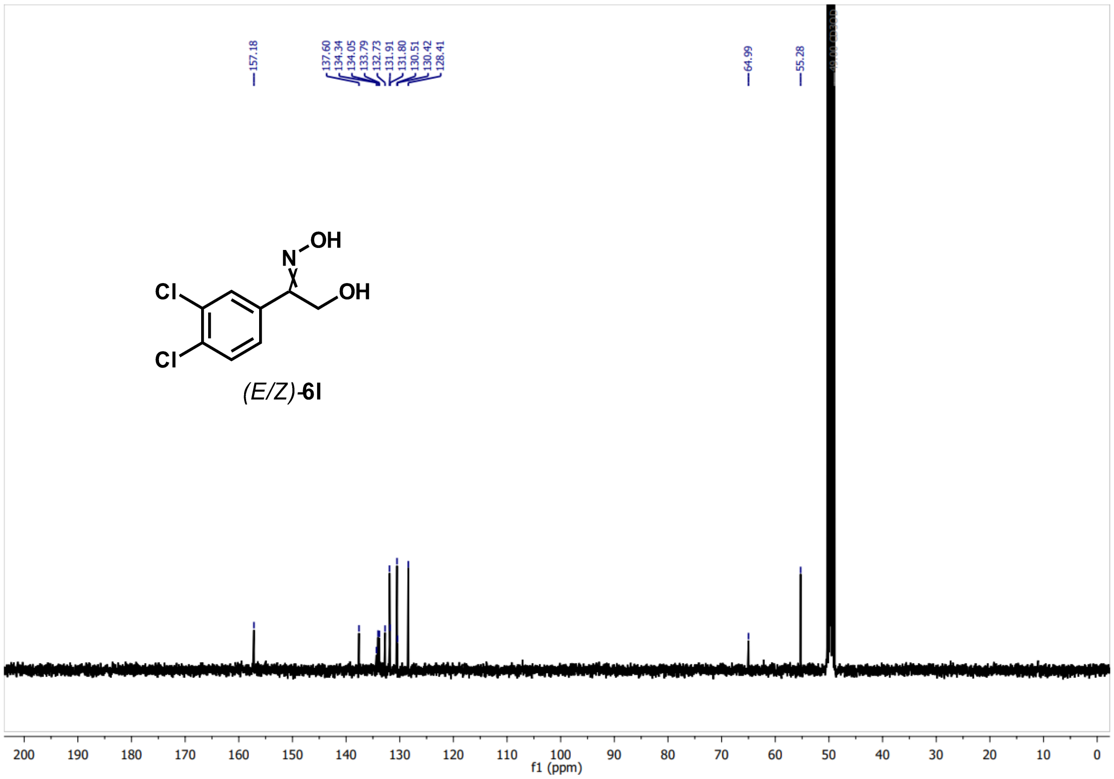
157.18
137.60
134.34
134.05
133.79
132.73
131.91
131.80
130.51
130.42
128.41
64.99
55.28
Cl
Cl
N
OH
OH
(E/Z)-6l
f1 (ppm)

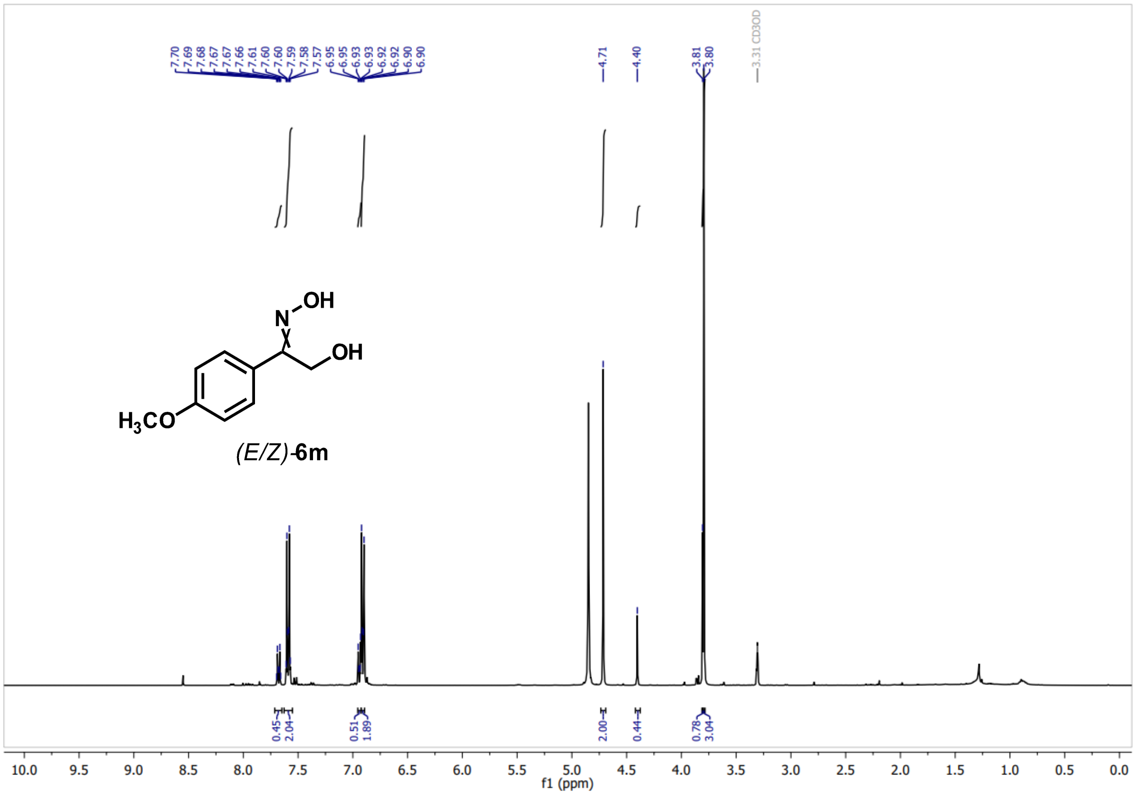
OH
N
OH
H3CO
(E/Z)-6m
7.70 7.69 7.68 7.67 7.67 7.66 7.61 7.60 7.60 7.59 7.58 7.57 6.95 6.95 6.93 6.93 6.92 6.92 6.90 6.90
4.71
4.40
3.81 3.80
3.31 CD3OD
0.45 2.04 0.51 1.89 2.00 0.44 0.78 3.04
f1 (ppm)

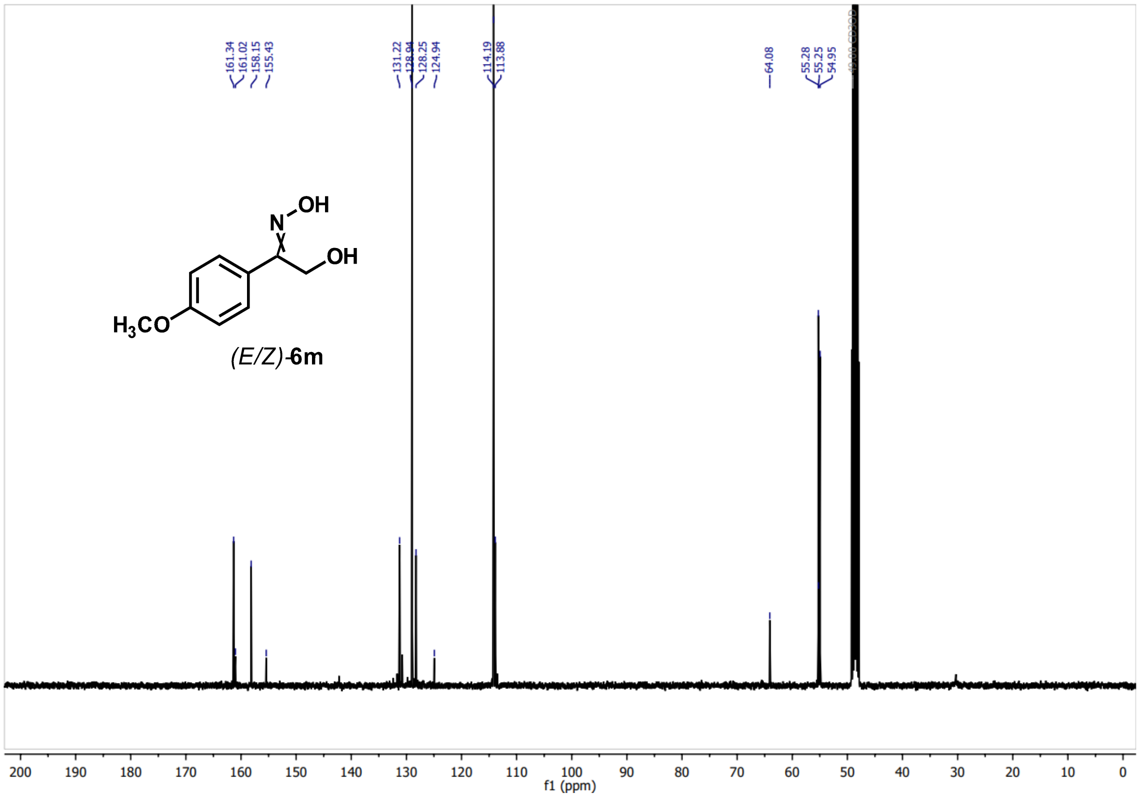
OH
N
OH
H3CO
(E/Z)-6m
161.34 161.02 158.15 155.43
131.22 128.94 128.25 124.94
114.19 113.88
64.08
55.28 55.25 54.95
f1 (ppm)

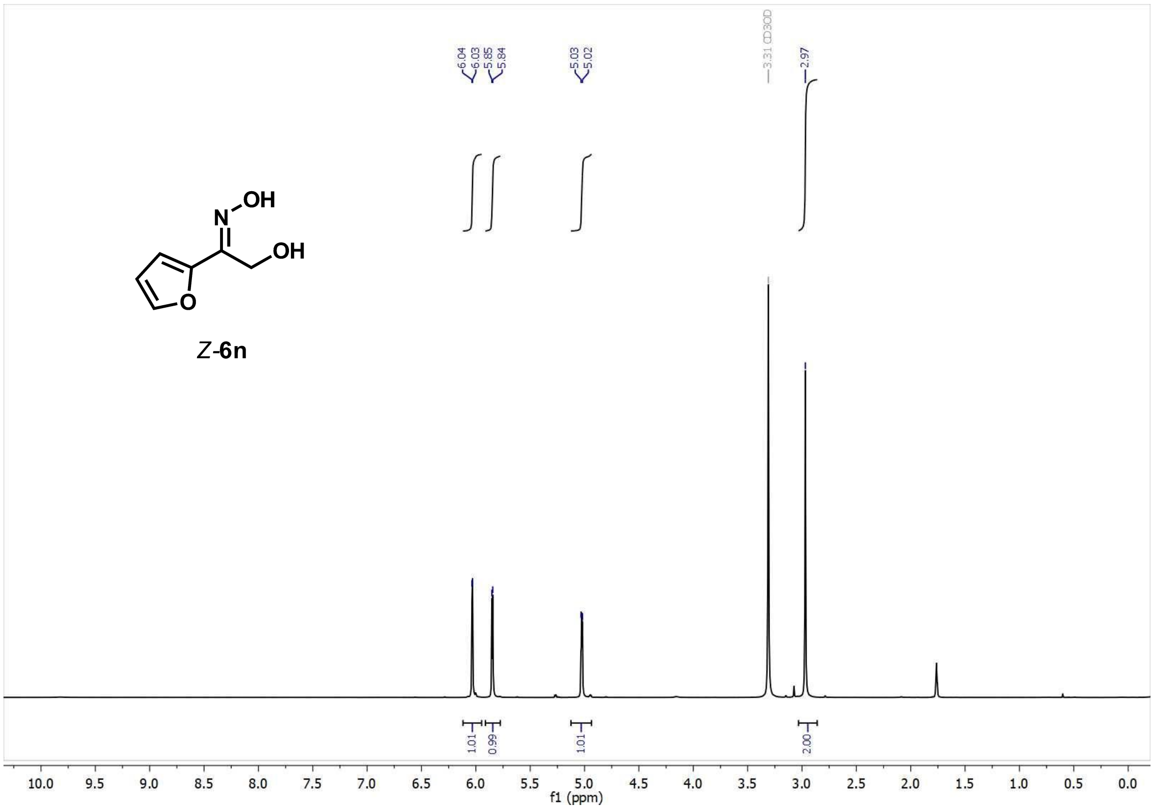
6.04
6.03
5.85
5.84
5.03
5.02
3.31 CD3OD
2.97
N
OH
OH
O
Z-6n
1.01
0.99
1.01
2.00
10.0 9.5 9.0 8.5 8.0 7.5 7.0 6.5 6.0 5.5 5.0 4.5 4.0 3.5 3.0 2.5 2.0 1.5 1.0 0.5 0.0
f1 (ppm)

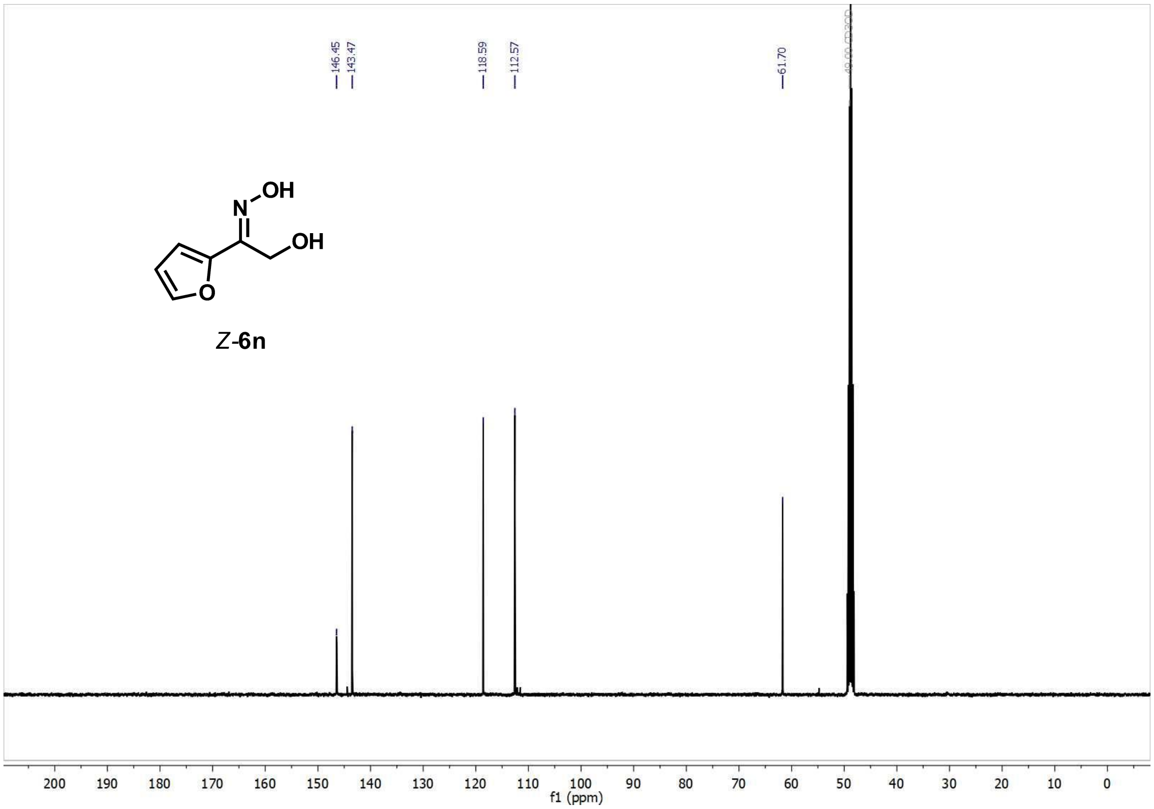
146.45
143.47
118.59
112.57
61.70
N
OH
OH
O
Z-6n
200 190 180 170 160 150 140 130 120 110 100 90 80 70 60 50 40 30 20 10 0
f1 (ppm)

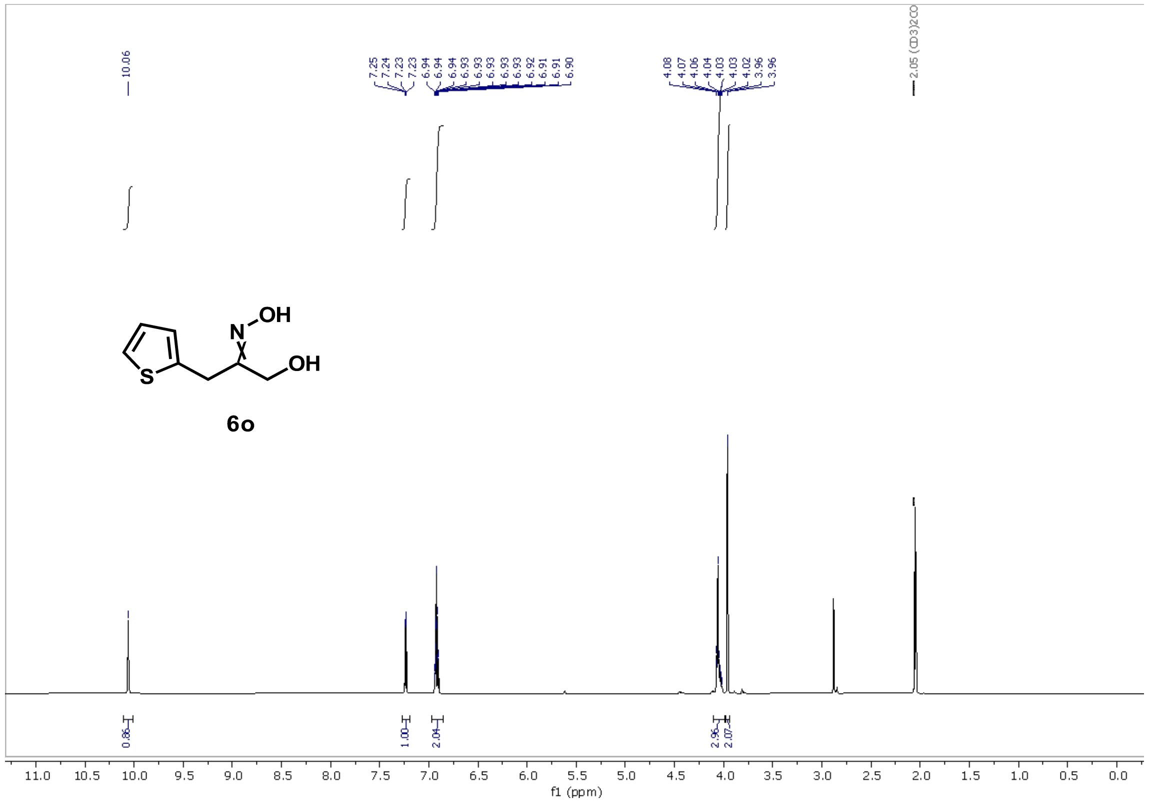

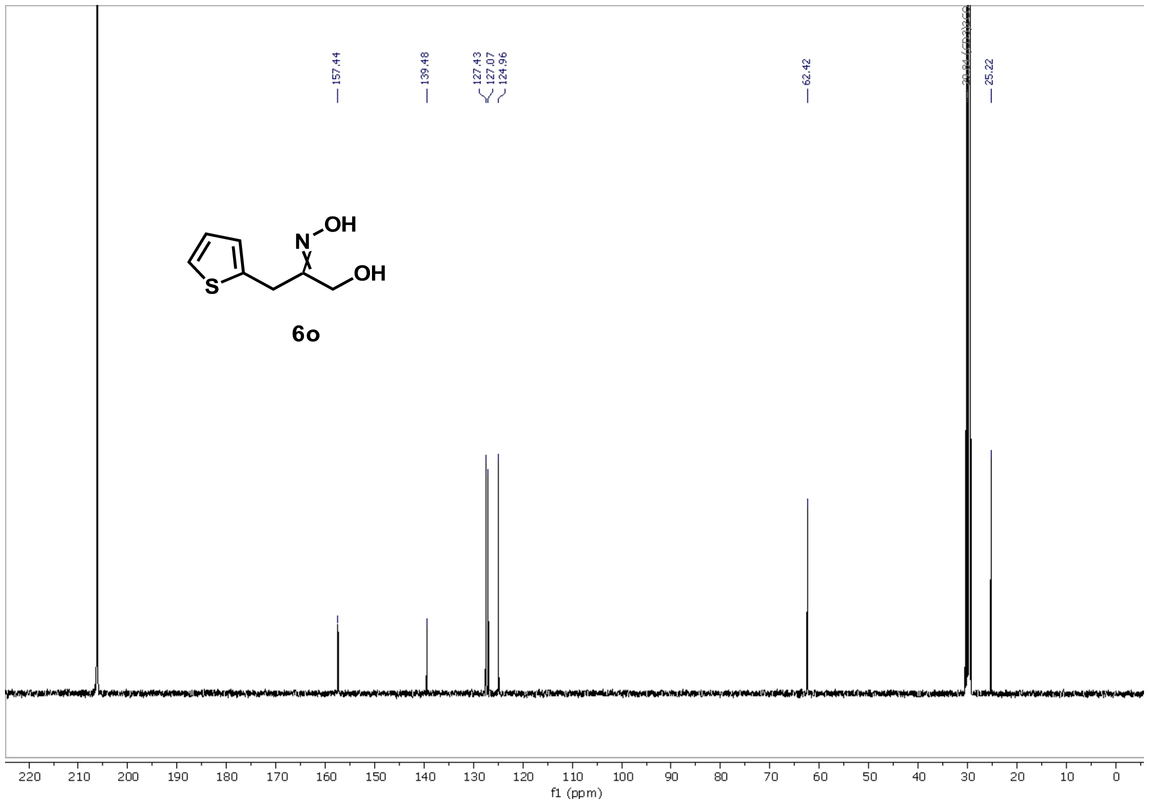

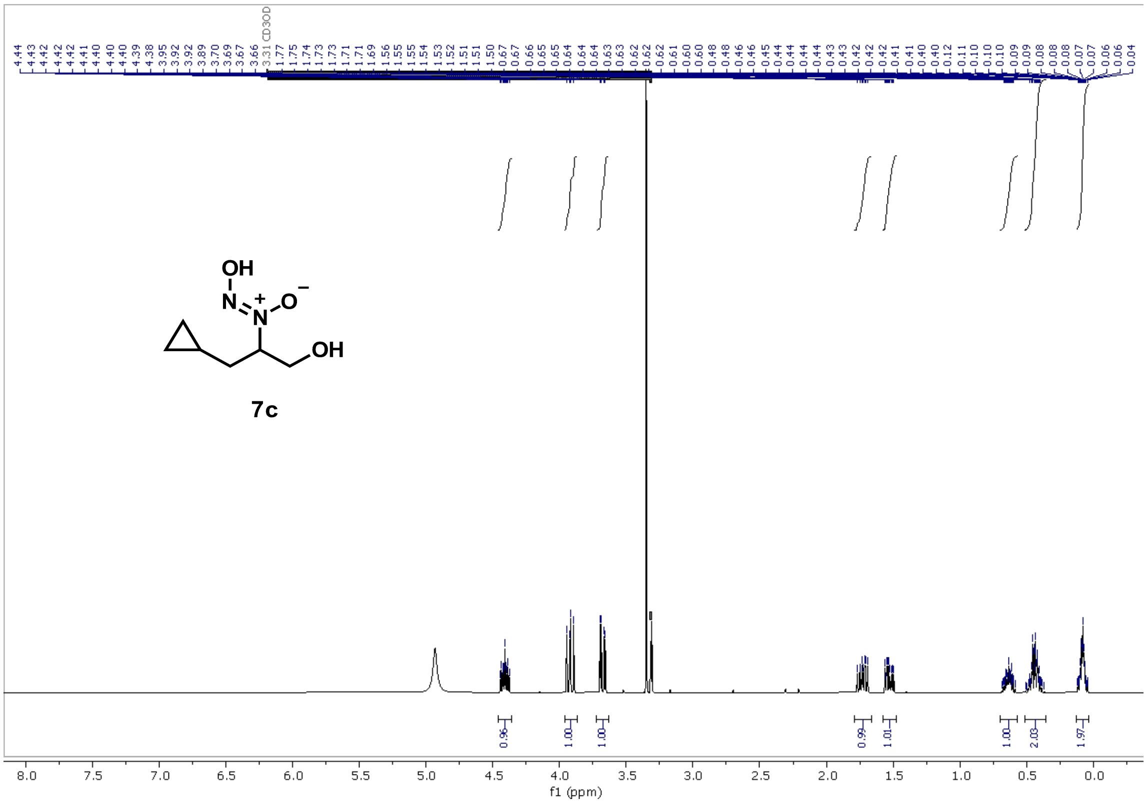
CD3OD
OH
N
N
O
OH
7c
f1 (ppm)

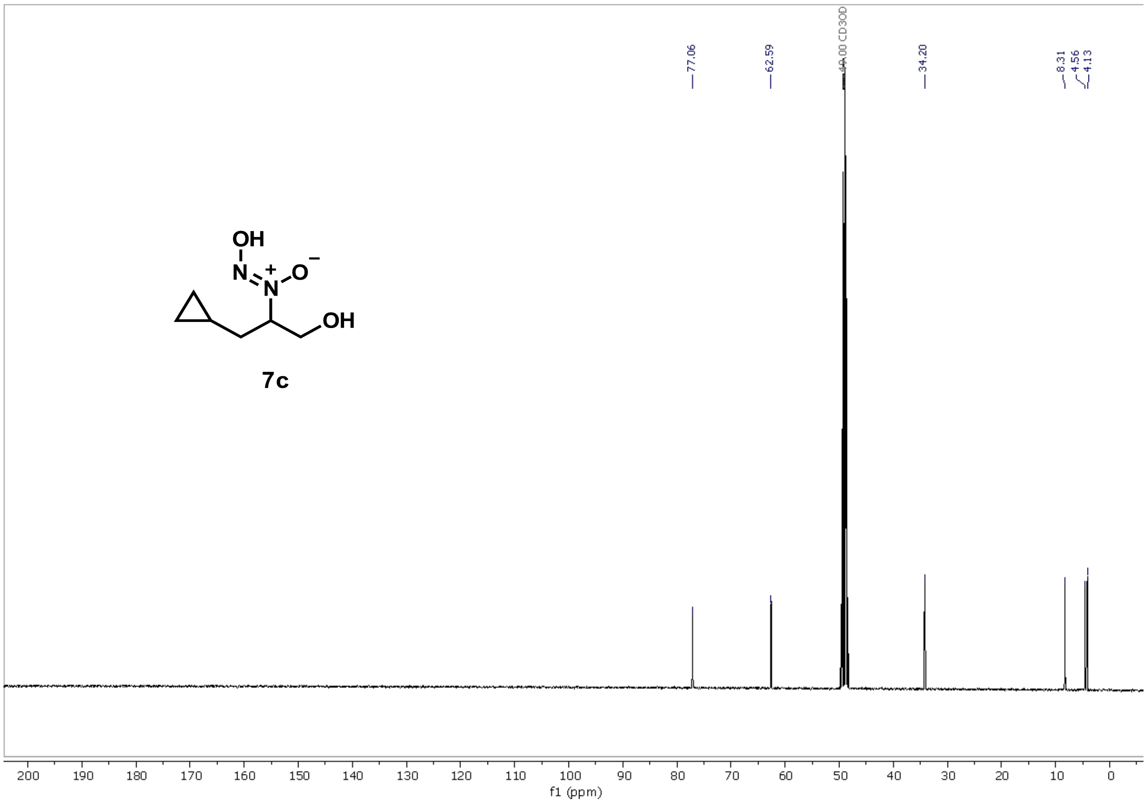
77.06
62.59
49.00 CD3OD
34.20
8.31
4.56
4.13
OH
N
N
O
OH
7c
f1 (ppm)

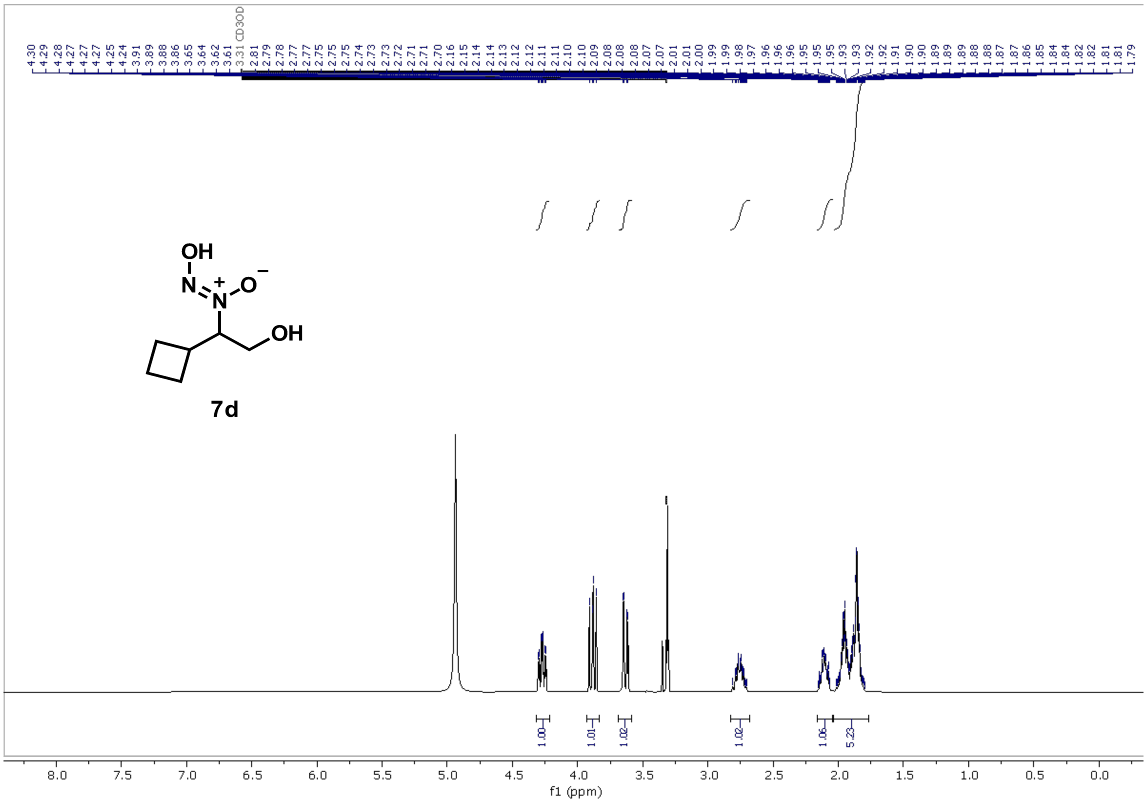
7d
f1 (ppm)

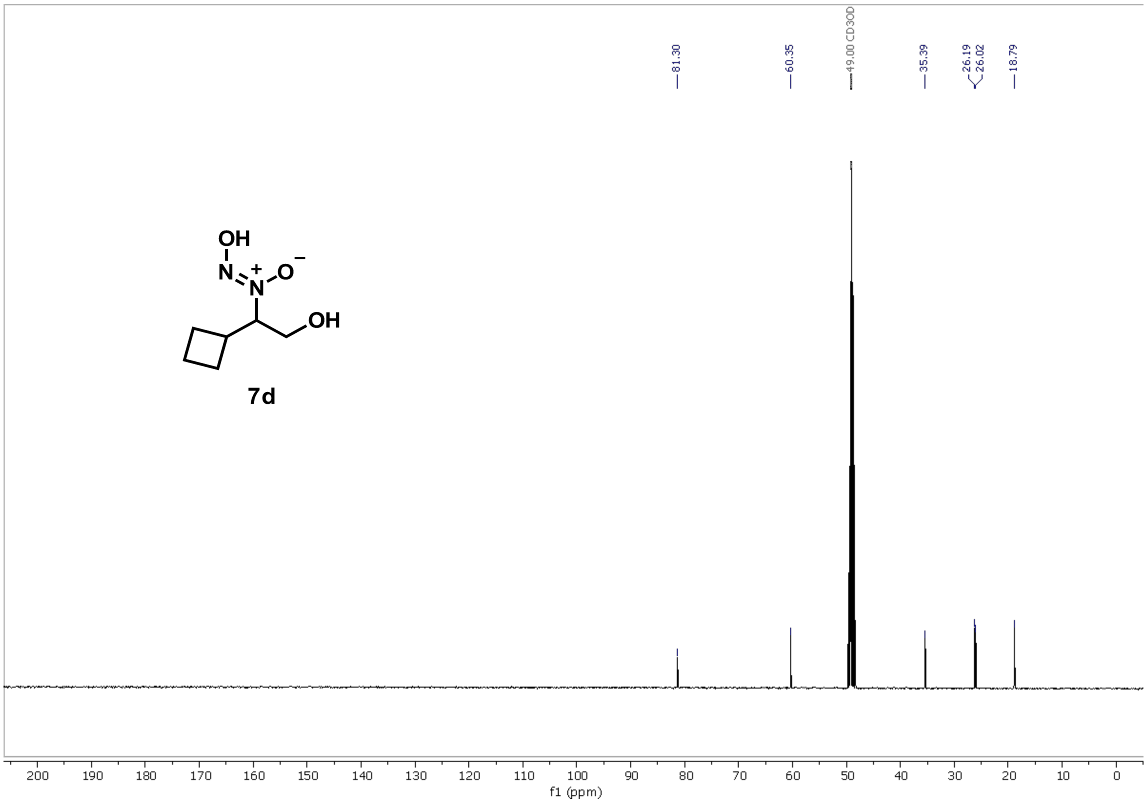
7d
81.30
60.35
49.00 CD3OD
35.39
26.19
26.02
18.79
f1 (ppm)

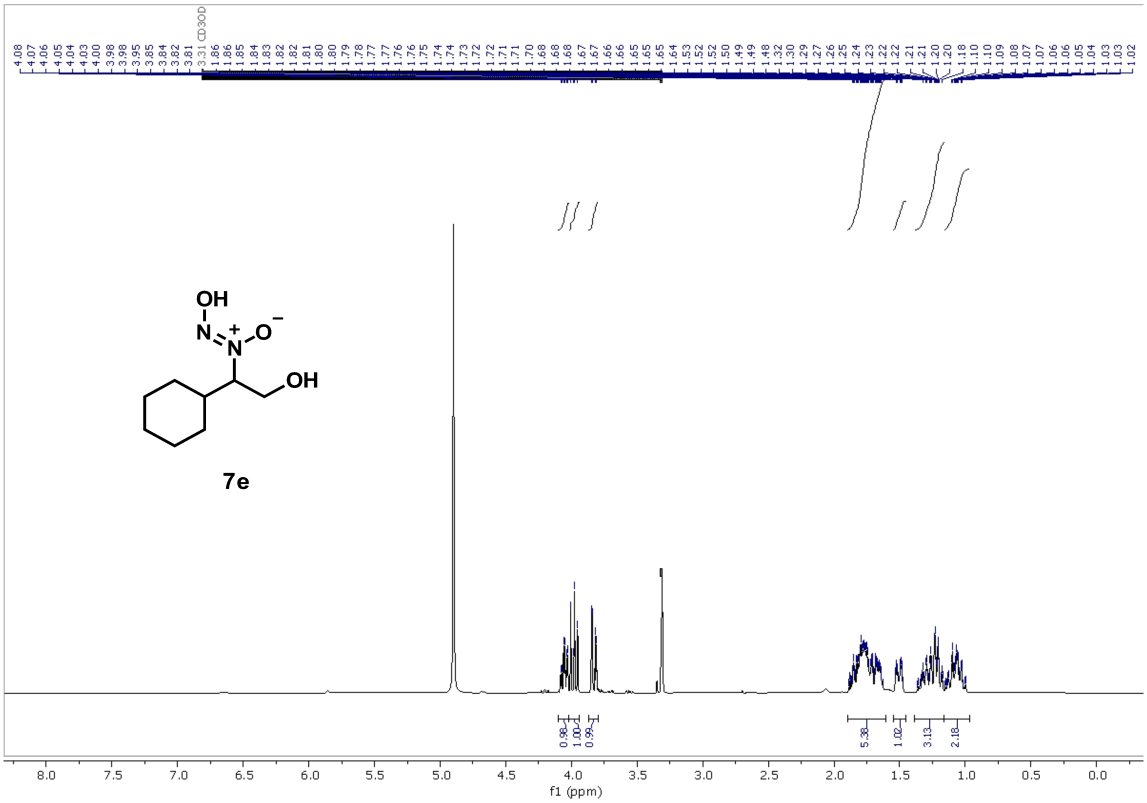

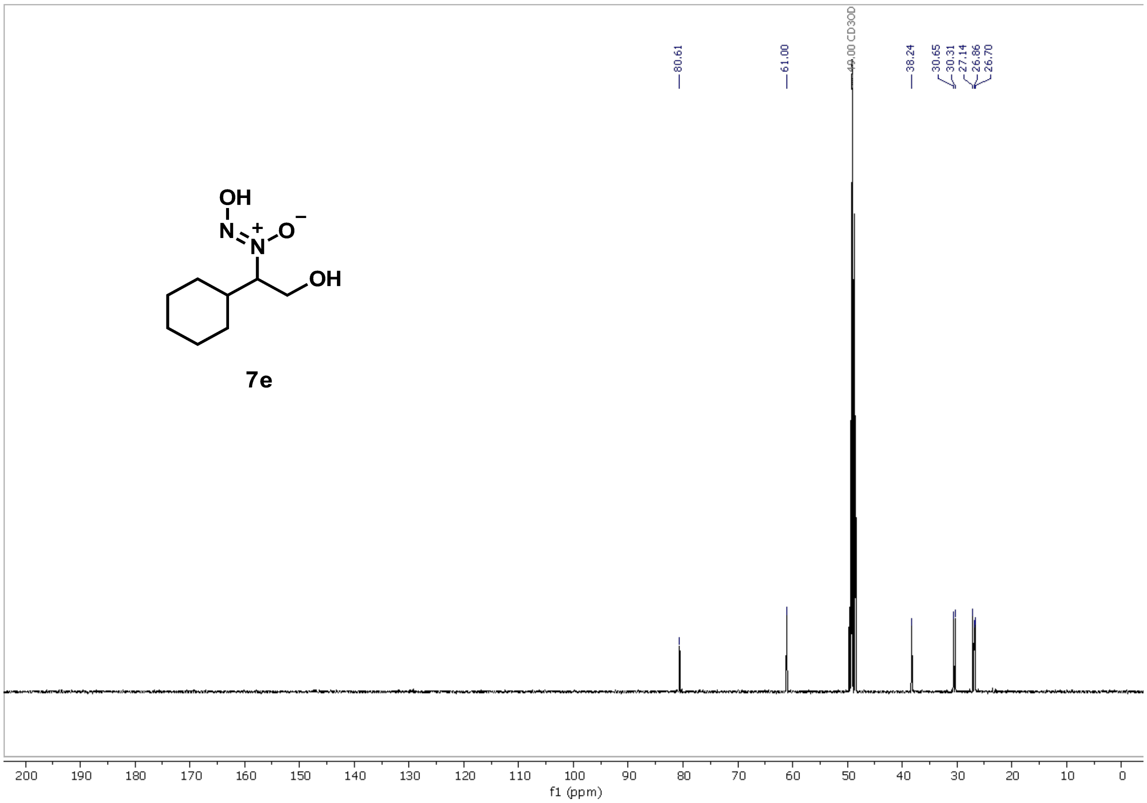

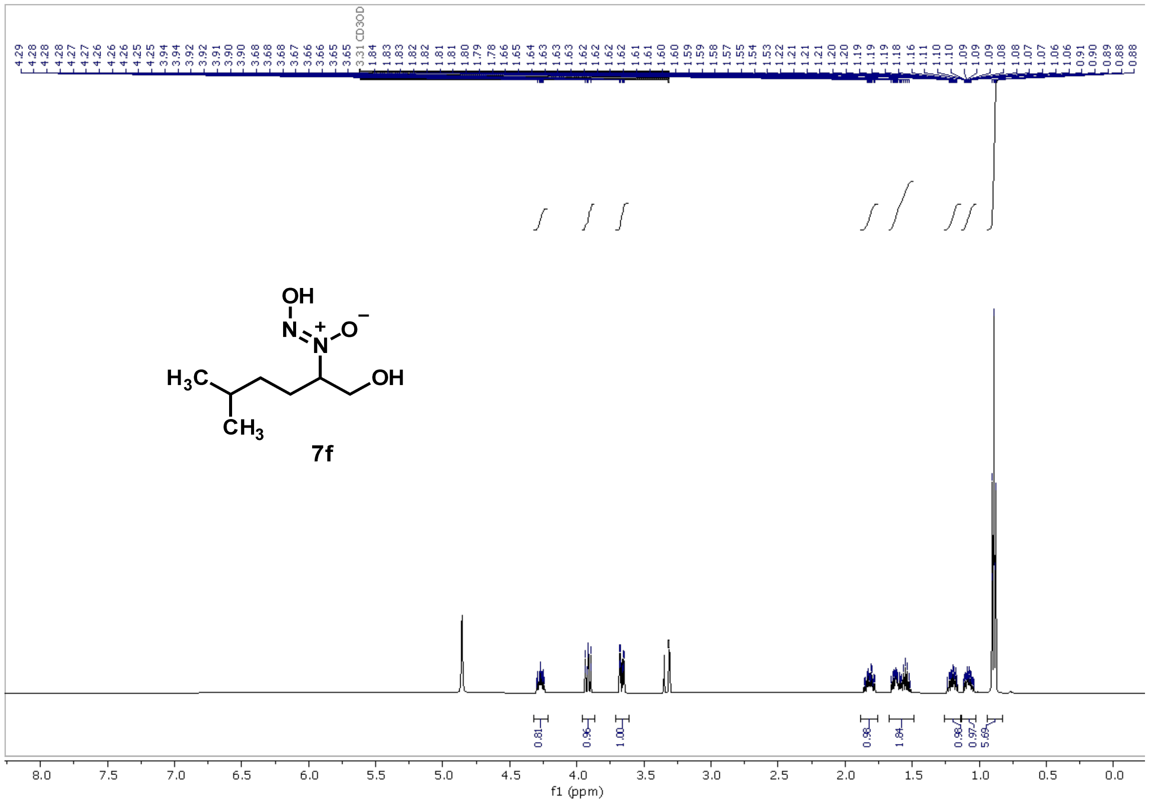

CD3OD
OH
N
O
H3C
OH
CH3
7f
0.81
0.96
1.00
0.98
1.84
0.98
0.97
5.69
f1 (ppm)


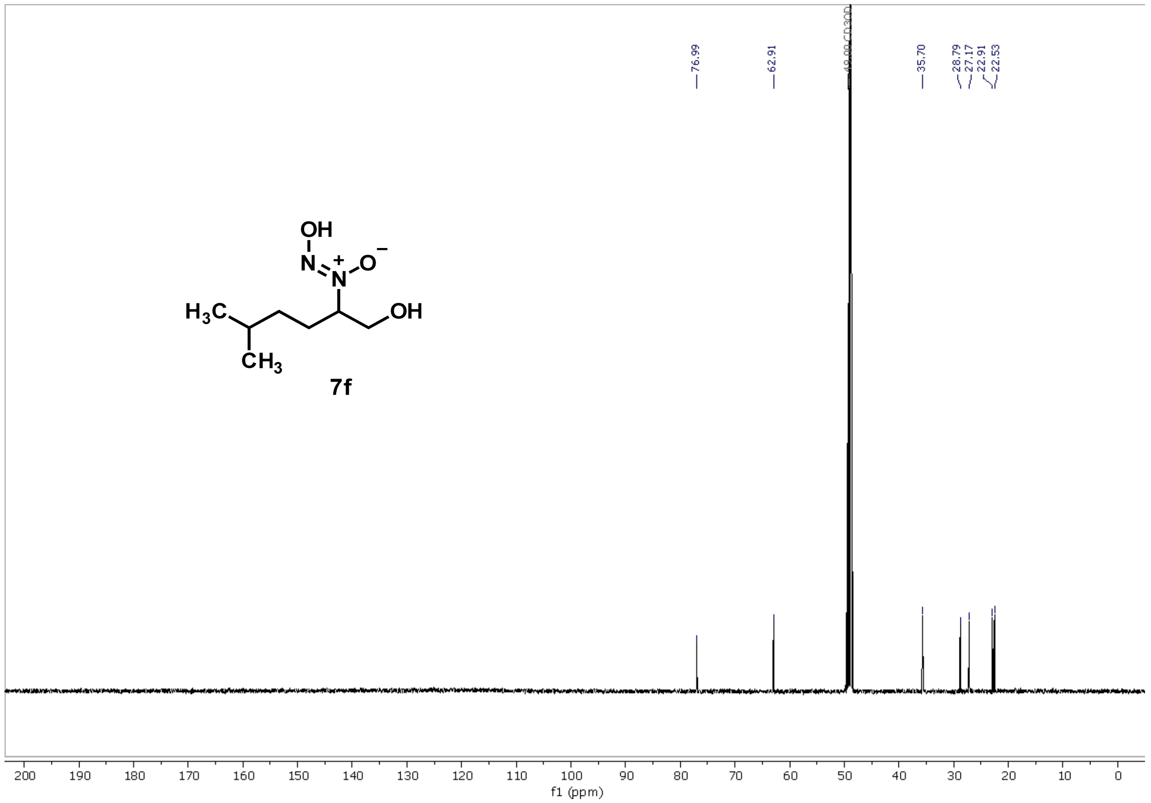

76.99
62.91
CD3OD
35.70
28.79
27.17
22.91
22.53
OH
N
O
H3C
OH
CH3
7f
f1 (ppm)

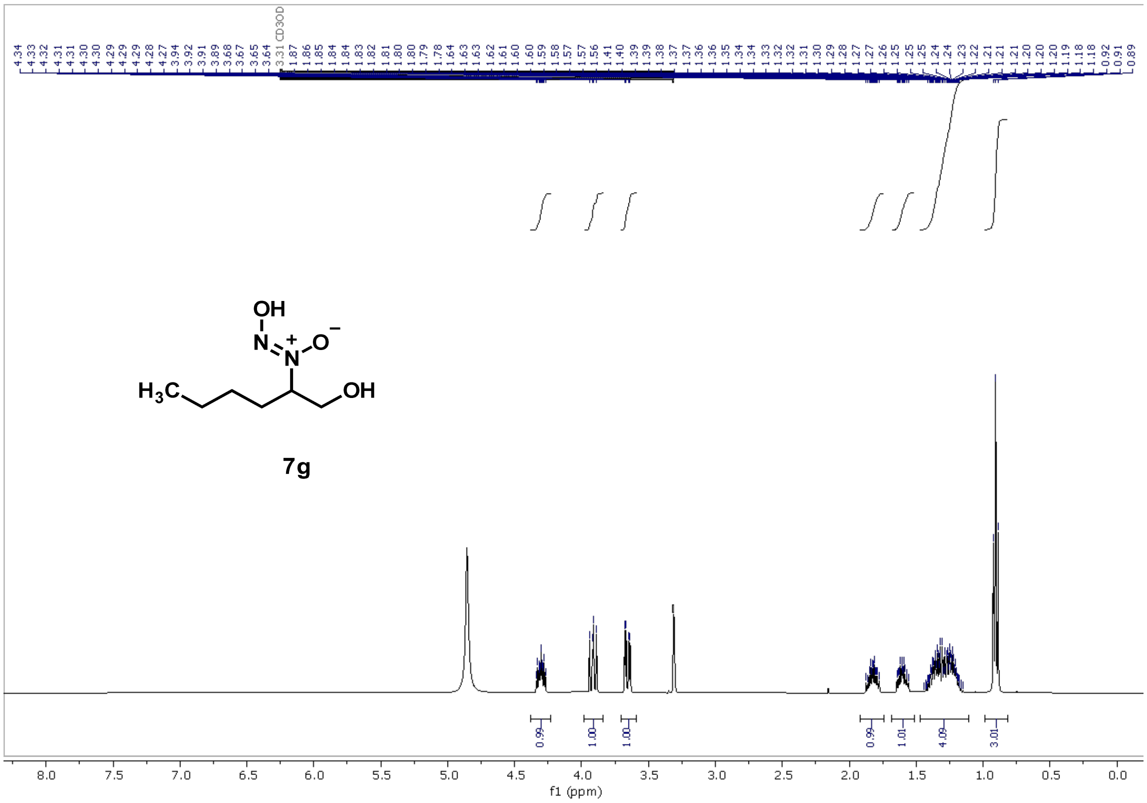
4.34 4.33 4.32 4.31 4.31 4.30 4.30 4.29 4.29 4.28 4.27 3.94 3.92 3.91 3.89 3.68 3.67 3.65 3.64 3.31 CD3OD 1.87 1.86 1.85 1.84 1.84 1.83 1.82 1.81 1.80 1.80 1.79 1.78 1.64 1.63 1.63 1.62 1.61 1.60 1.60 1.59 1.58 1.57 1.57 1.56 1.41 1.40 1.39 1.39 1.38 1.37 1.37 1.36 1.36 1.35 1.34 1.34 1.33 1.32 1.32 1.31 1.30 1.29 1.28 1.27 1.27 1.26 1.25 1.25 1.24 1.24 1.23 1.22 1.21 1.21 1.21 1.20 1.20 1.19 1.18 1.18 0.92 0.91 0.89
OH
N
N
O
$H_3C$
OH
7g
0.99
1.00
1.00
0.99
1.01
4.09
3.01
8.0 7.5 7.0 6.5 6.0 5.5 5.0 4.5 4.0 3.5 3.0 2.5 2.0 1.5 1.0 0.5 0.0
f1 (ppm)

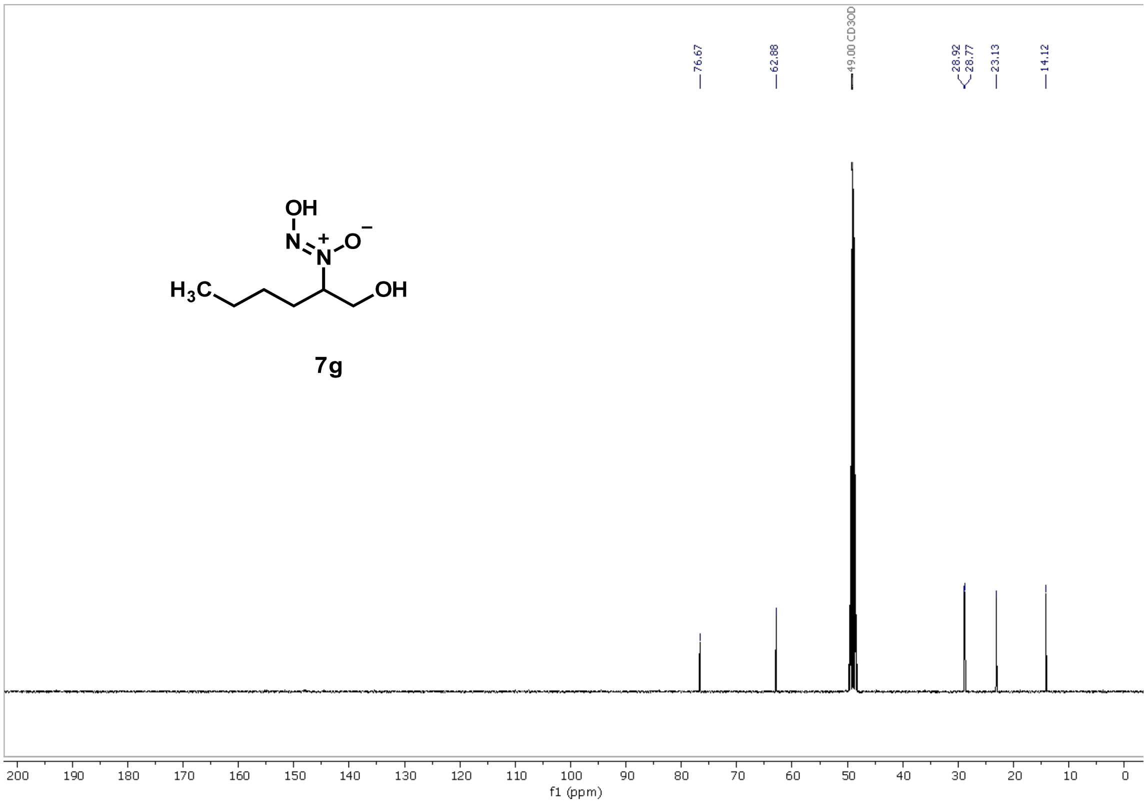
76.67
62.88
49.00 CD3OD
28.92
28.77
23.13
14.12
OH
N
N
O
$H_3C$
OH
7g
200 190 180 170 160 150 140 130 120 110 100 90 80 70 60 50 40 30 20 10 0
f1 (ppm)

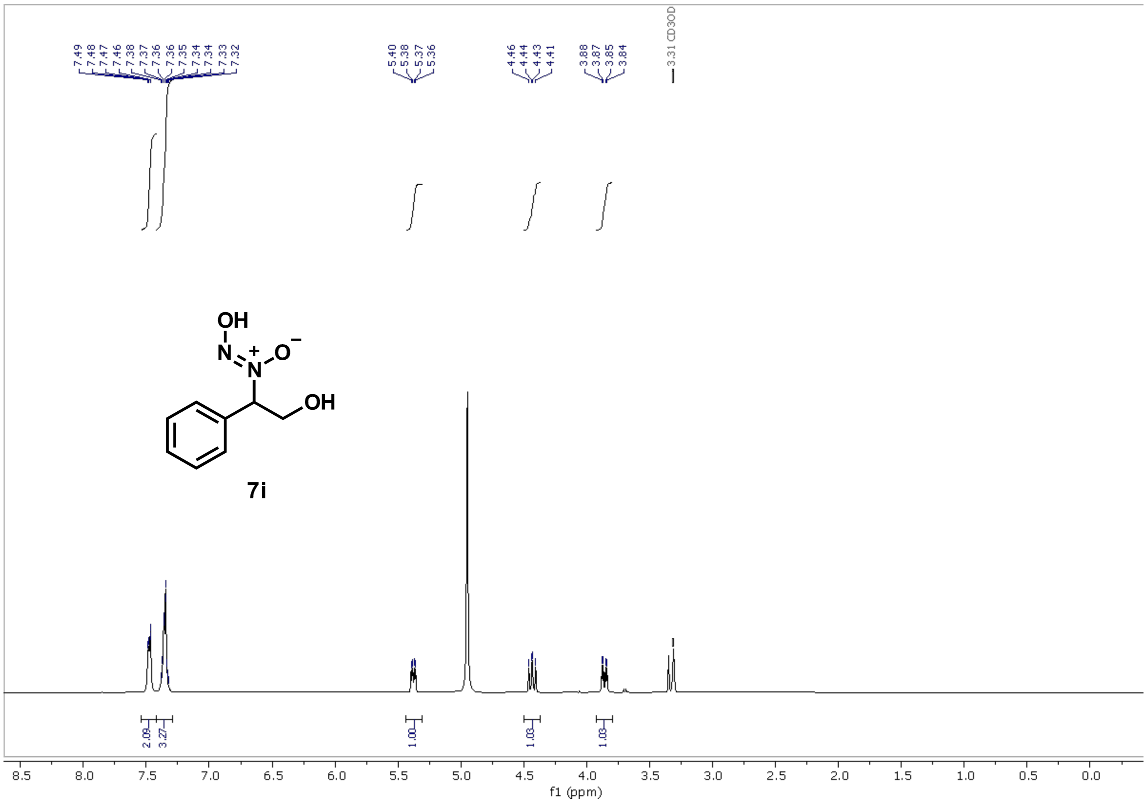
7.49
7.48
7.47
7.46
7.38
7.37
7.36
7.36
7.35
7.34
7.34
7.33
7.32
5.40
5.38
5.37
5.36
4.46
4.44
4.43
4.41
3.88
3.87
3.85
3.84
3.31 CD3OD
OH
N
N
O
+
−
OH
7i
2.09
3.27
1.00
1.03
1.03
8.5
8.0
7.5
7.0
6.5
6.0
5.5
5.0
4.5
4.0
3.5
3.0
2.5
2.0
1.5
1.0
0.5
0.0
f1 (ppm)

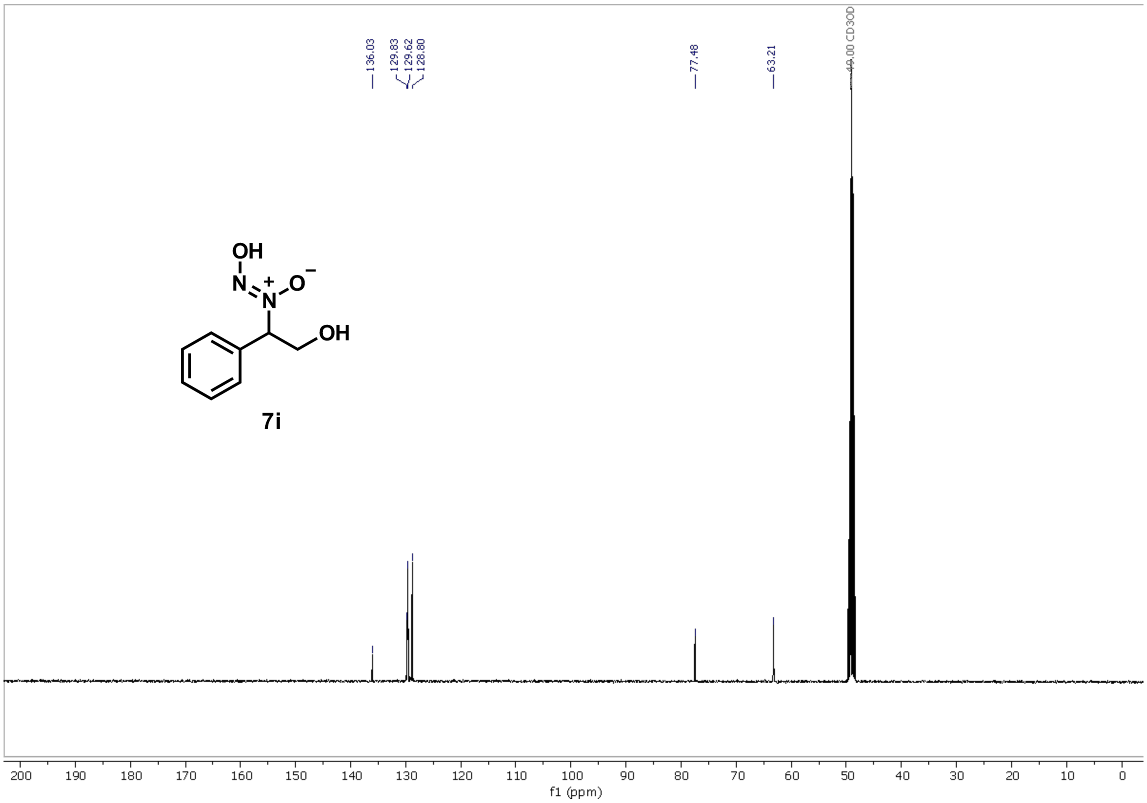
136.03
129.83
129.62
128.80
77.48
63.21
49.00 CD3OD
OH
N
N
O
+
−
OH
7i
200
190
180
170
160
150
140
130
120
110
100
90
80
70
60
50
40
30
20
10
0
f1 (ppm)

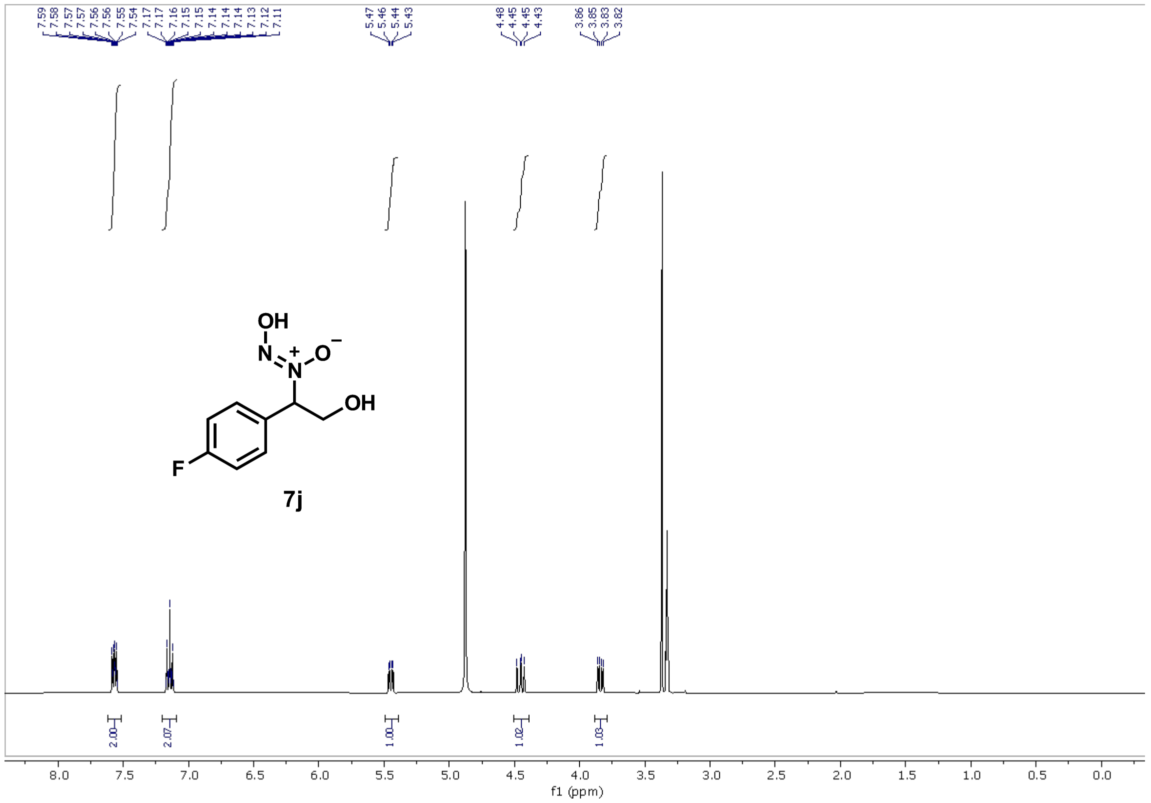
7j
f1 (ppm)

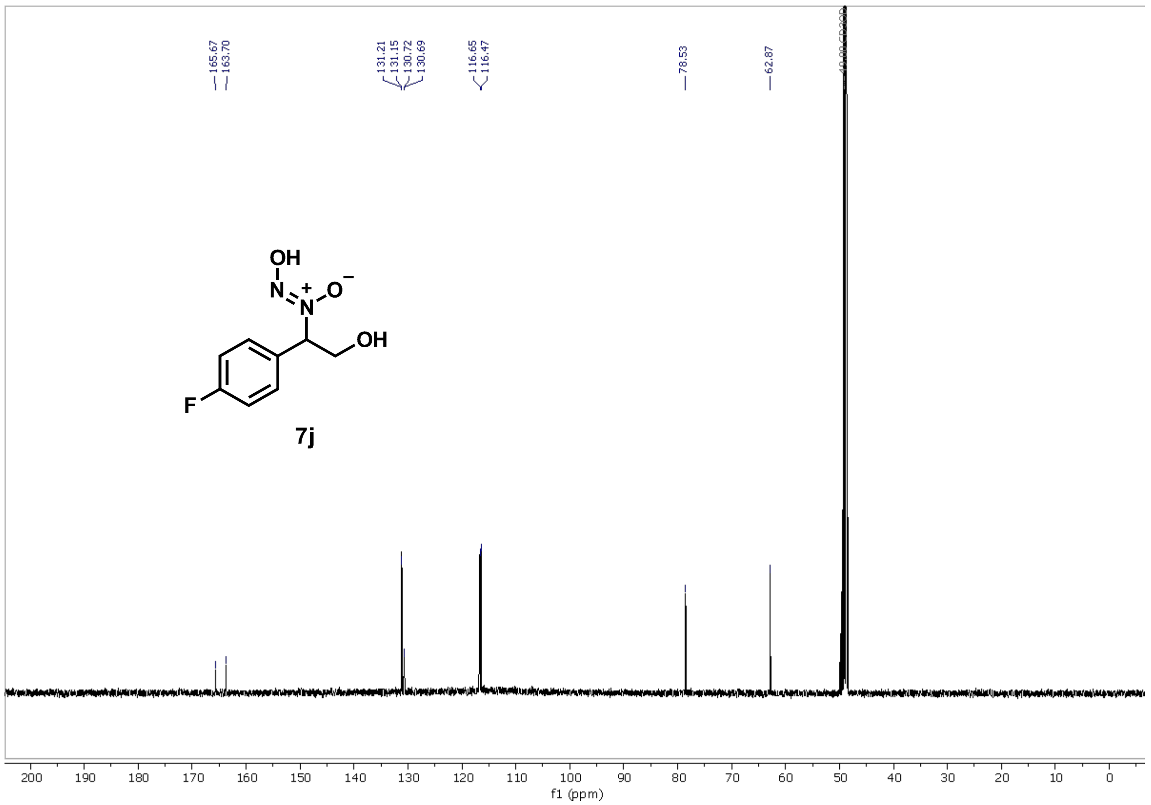
7j
f1 (ppm)

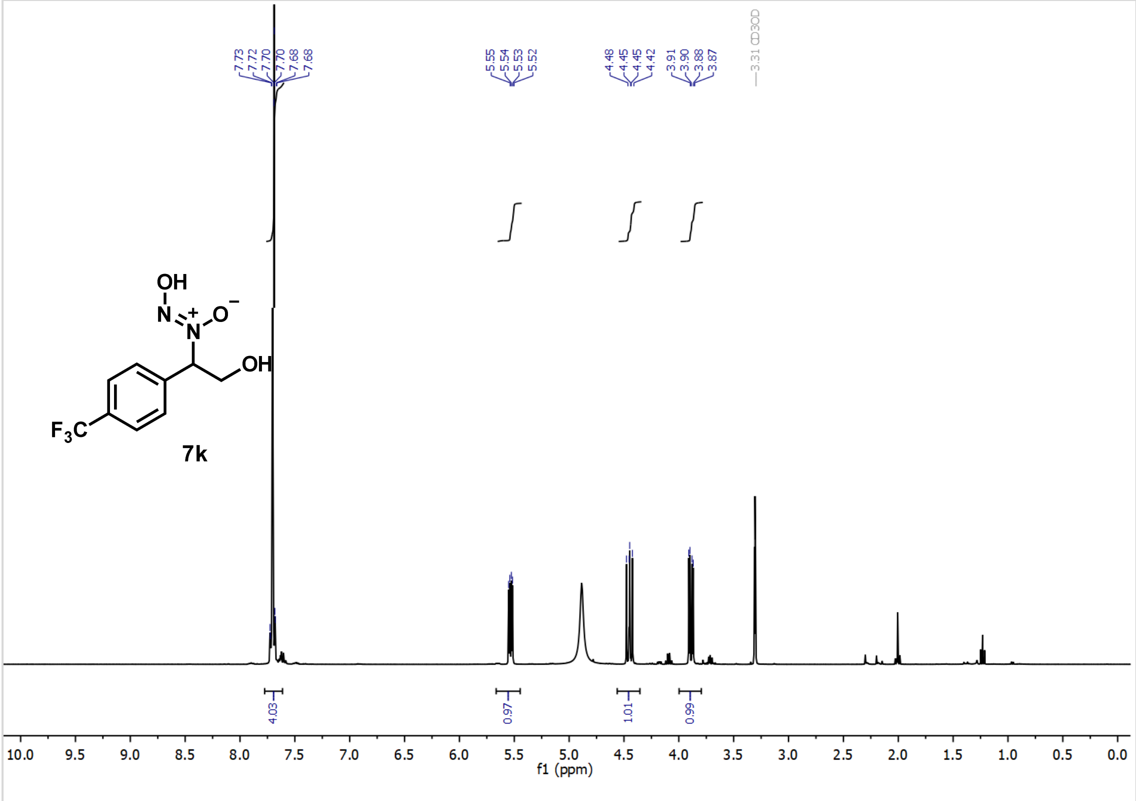
7.73
7.72
7.70
7.70
7.68
7.68
5.55
5.54
5.53
5.52
4.48
4.45
4.45
4.42
3.91
3.90
3.88
3.87
3.31 CD3OD
OH
N
N
O
OH
F3C
7k
4.03
0.97
1.01
0.99
f1 (ppm)

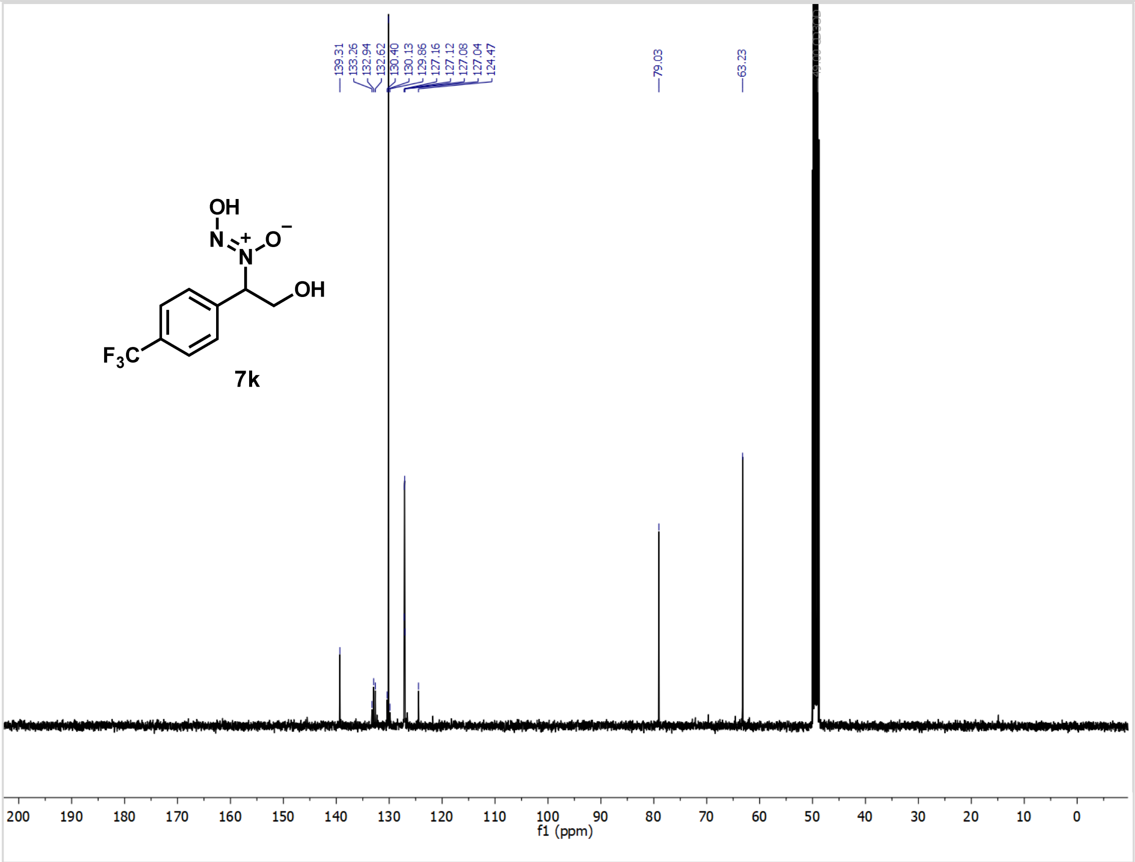
139.31
133.26
132.94
132.62
130.40
130.13
129.86
127.16
127.12
127.08
127.04
124.47
79.03
63.23
OH
N
N
O
OH
F3C
7k
f1 (ppm)

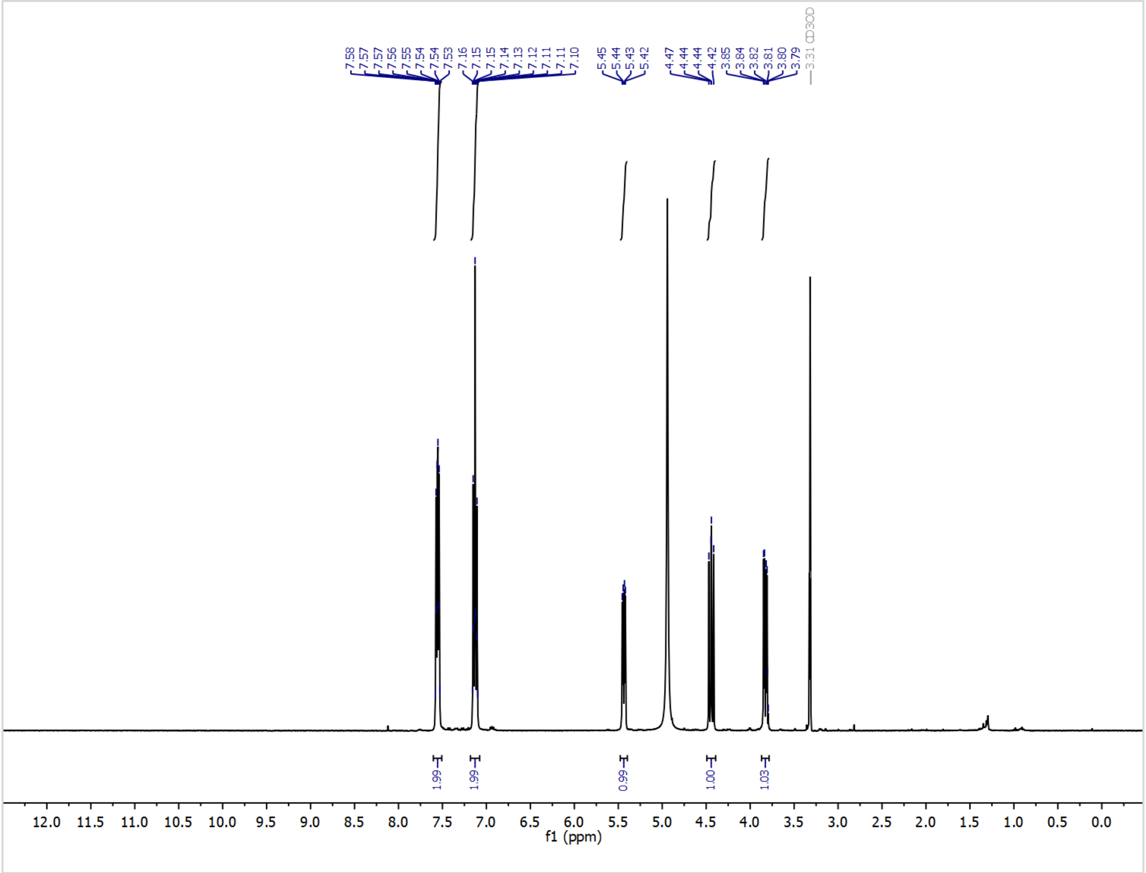
7.58
7.57
7.57
7.56
7.55
7.54
7.54
7.53
7.16
7.15
7.15
7.14
7.13
7.12
7.11
7.11
7.10
5.45
5.44
5.43
5.42
4.47
4.44
4.44
4.42
3.85
3.84
3.82
3.81
3.80
3.79
3.31 CD3OD
1.99
1.99
0.99
1.00
1.03
12.0 11.5 11.0 10.5 10.0 9.5 9.0 8.5 8.0 7.5 7.0 6.5 6.0 5.5 5.0 4.5 4.0 3.5 3.0 2.5 2.0 1.5 1.0 0.5 0.0
f1 (ppm)

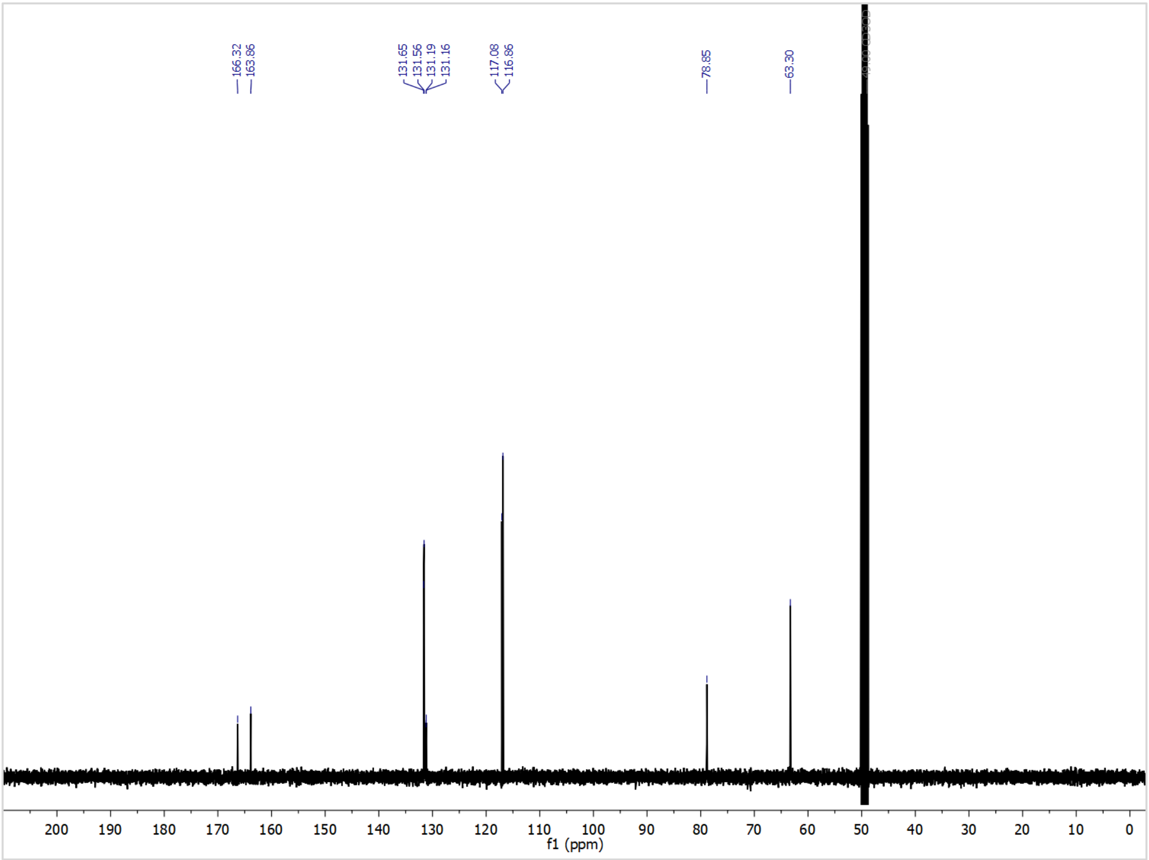
166.32
163.86
131.65
131.56
131.19
131.16
117.08
116.86
78.85
63.30
200 190 180 170 160 150 140 130 120 110 100 90 80 70 60 50 40 30 20 10 0
f1 (ppm)

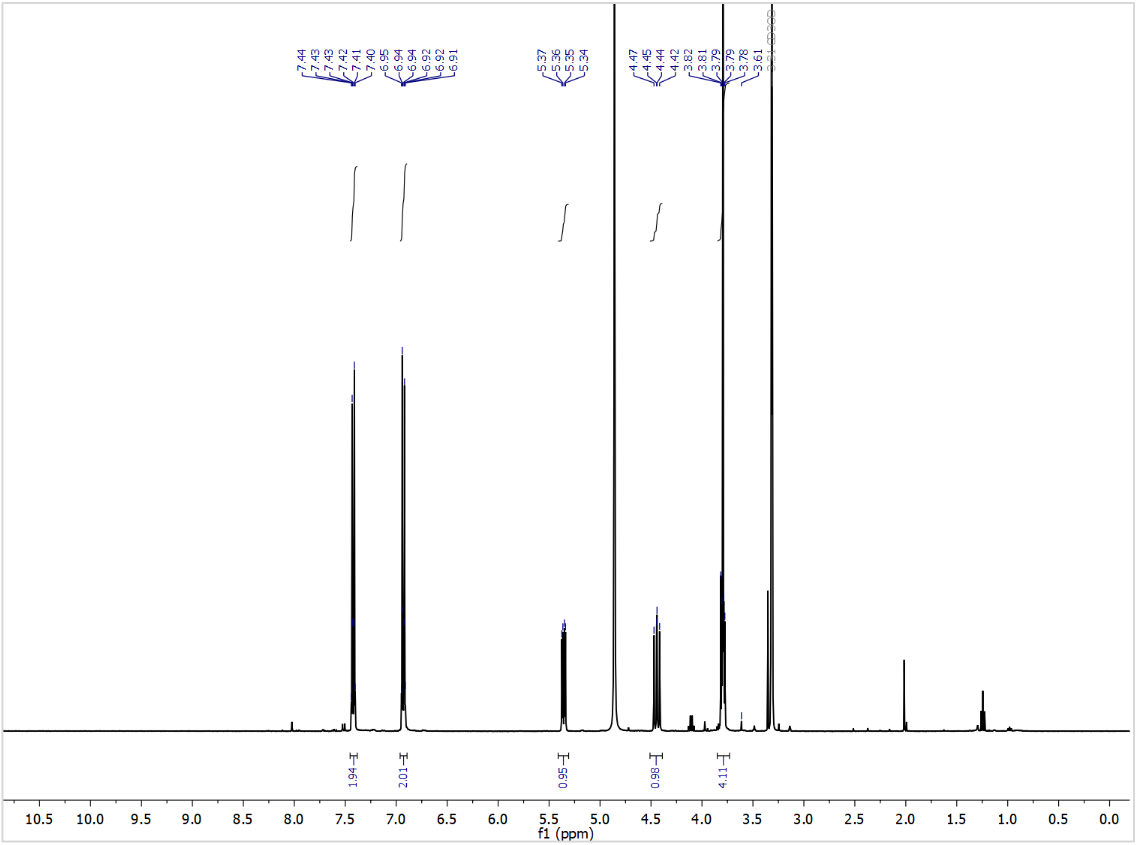
7.44
7.43
7.43
7.42
7.41
7.40
6.95
6.94
6.94
6.92
6.92
6.91
5.37
5.36
5.35
5.34
4.47
4.45
4.44
4.42
3.82
3.81
3.79
3.79
3.78
3.61
1.94
2.01
0.95
0.98
4.11
10.5 10.0 9.5 9.0 8.5 8.0 7.5 7.0 6.5 6.0 5.5 5.0 4.5 4.0 3.5 3.0 2.5 2.0 1.5 1.0 0.5 0.0
f1 (ppm)

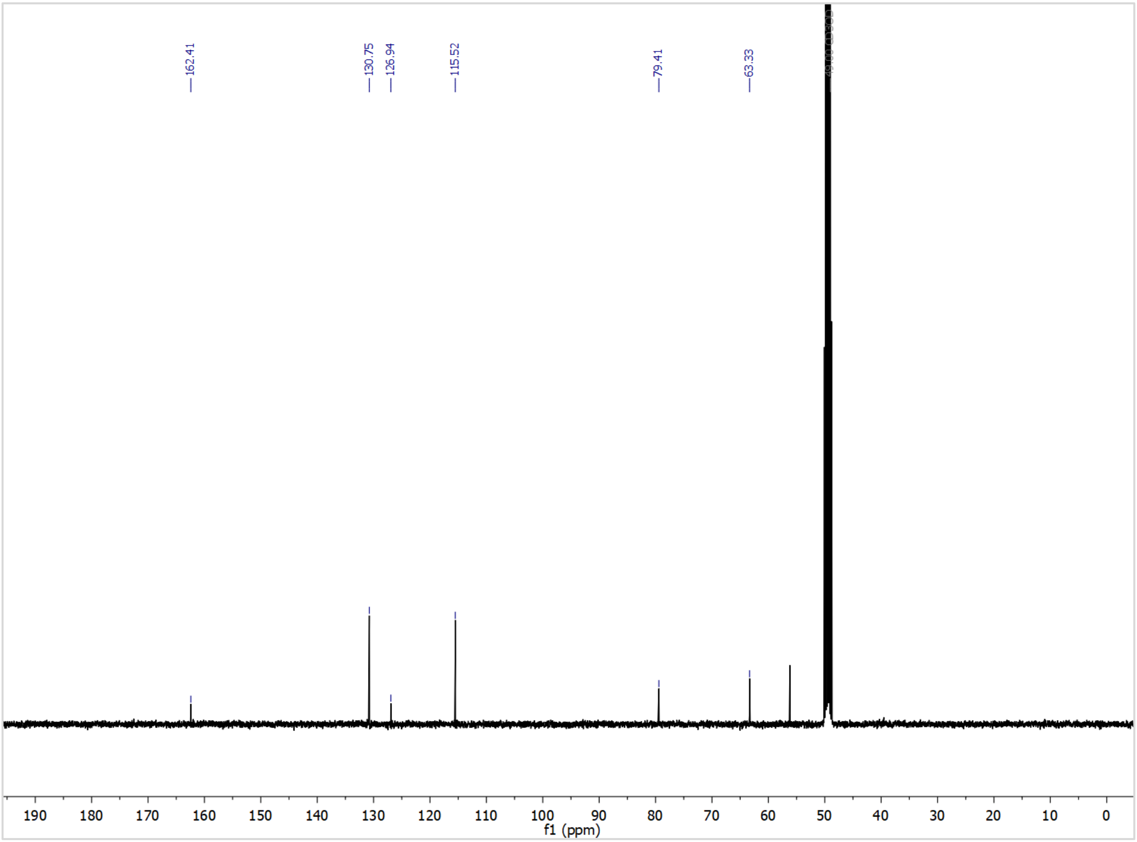
162.41
130.75
126.94
115.52
79.41
63.33
190 180 170 160 150 140 130 120 110 100 90 80 70 60 50 40 30 20 10 0
f1 (ppm)

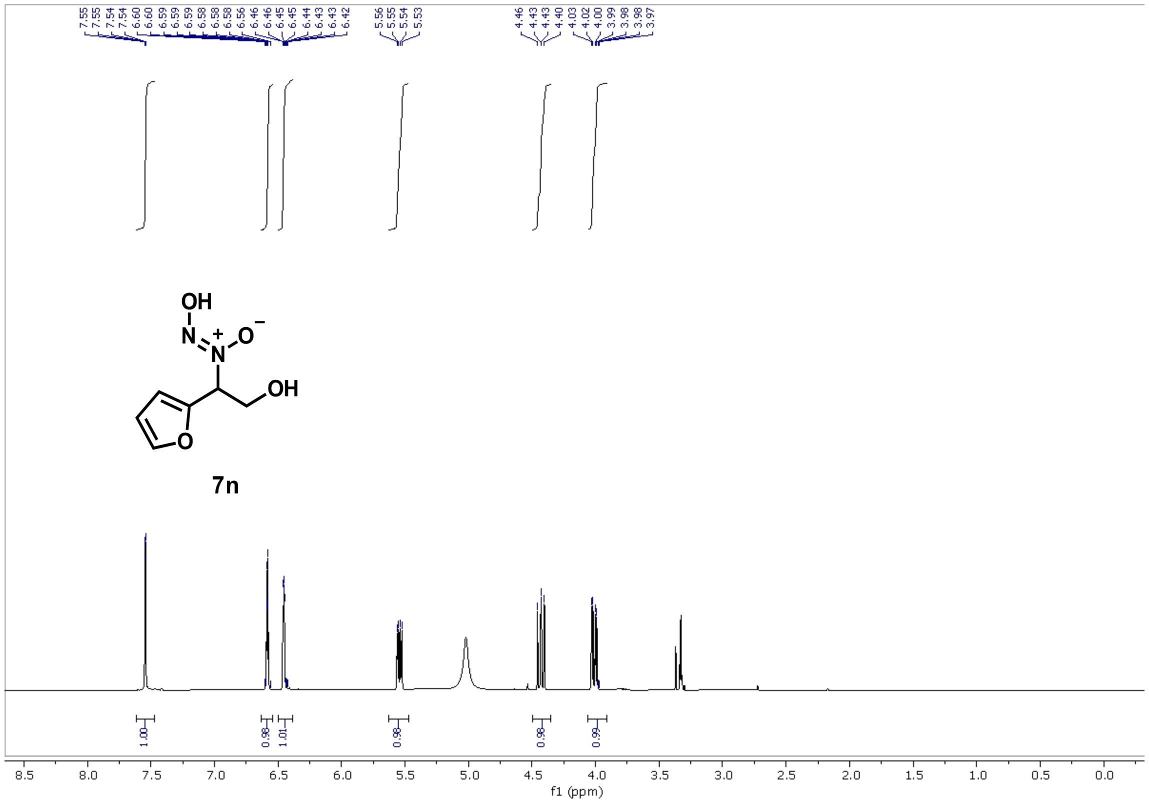

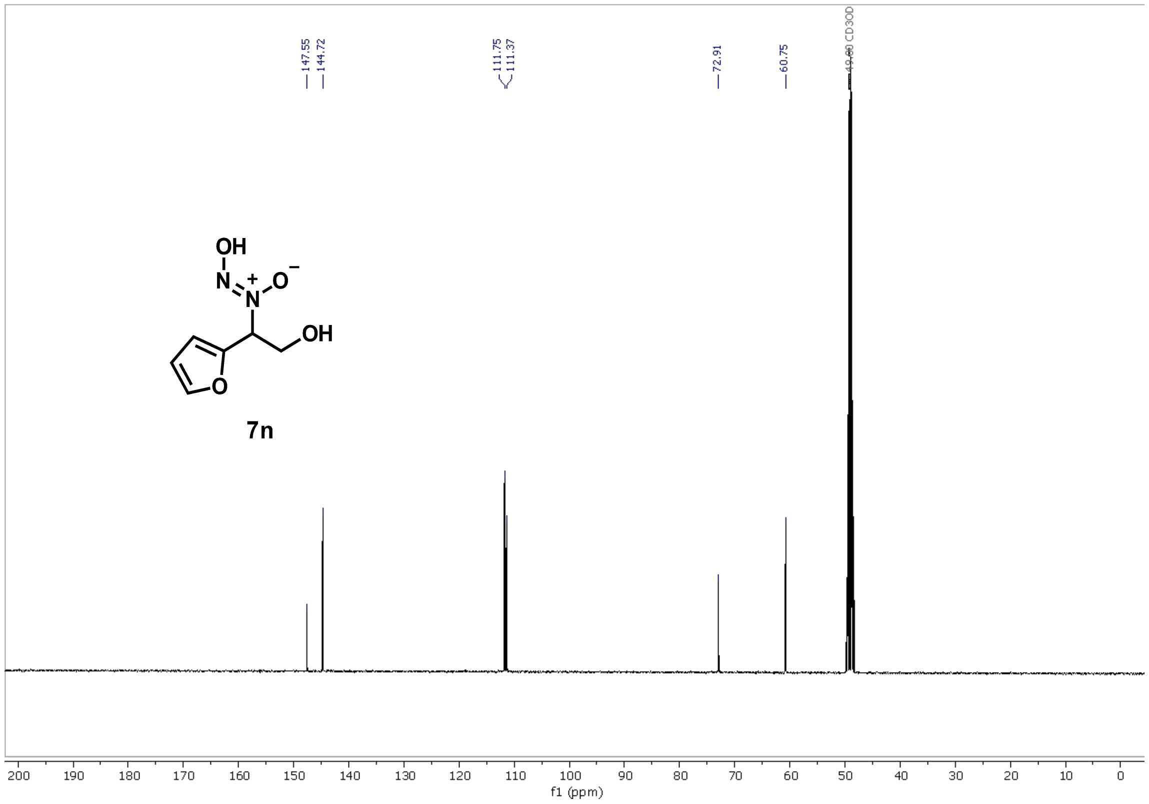

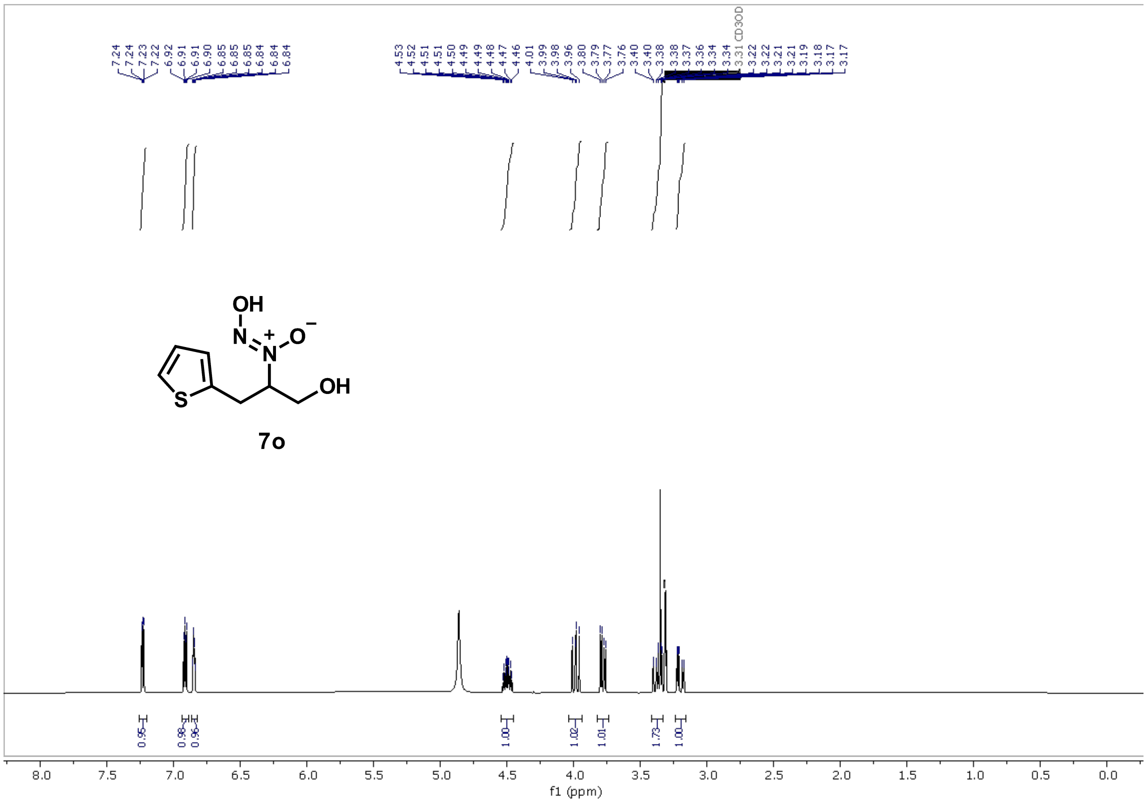
7o
f1 (ppm)

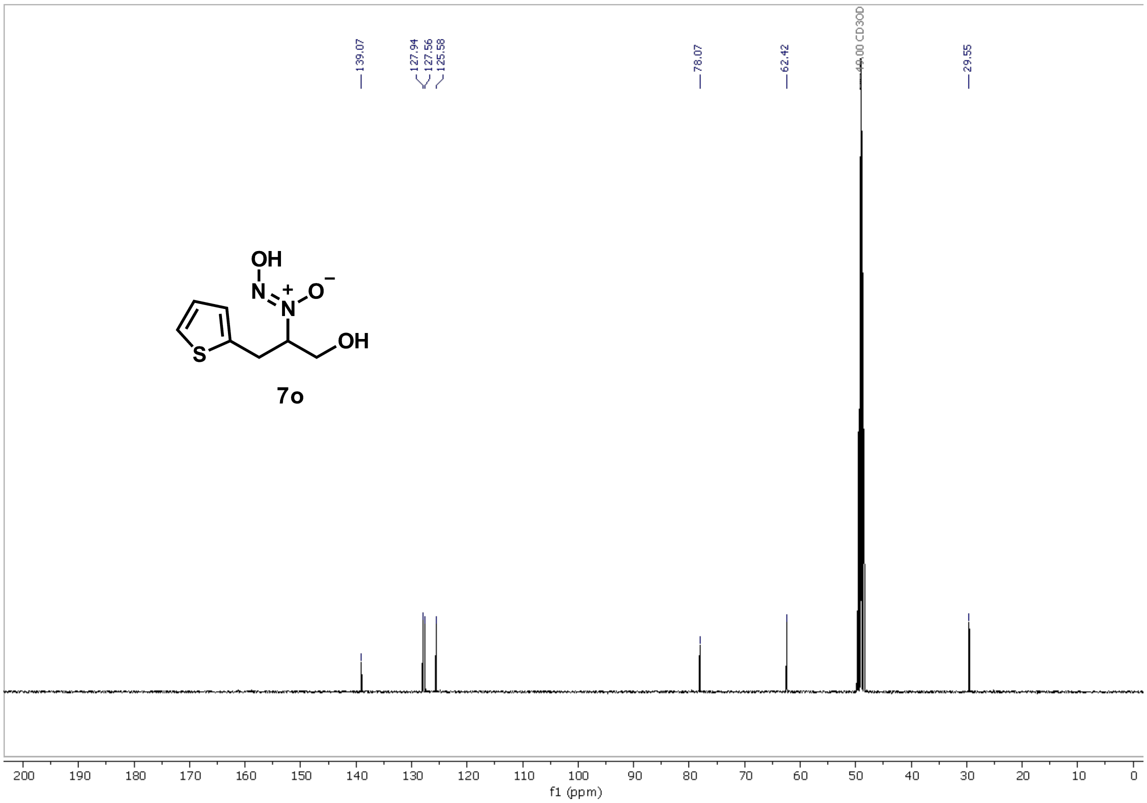
7o
f1 (ppm)

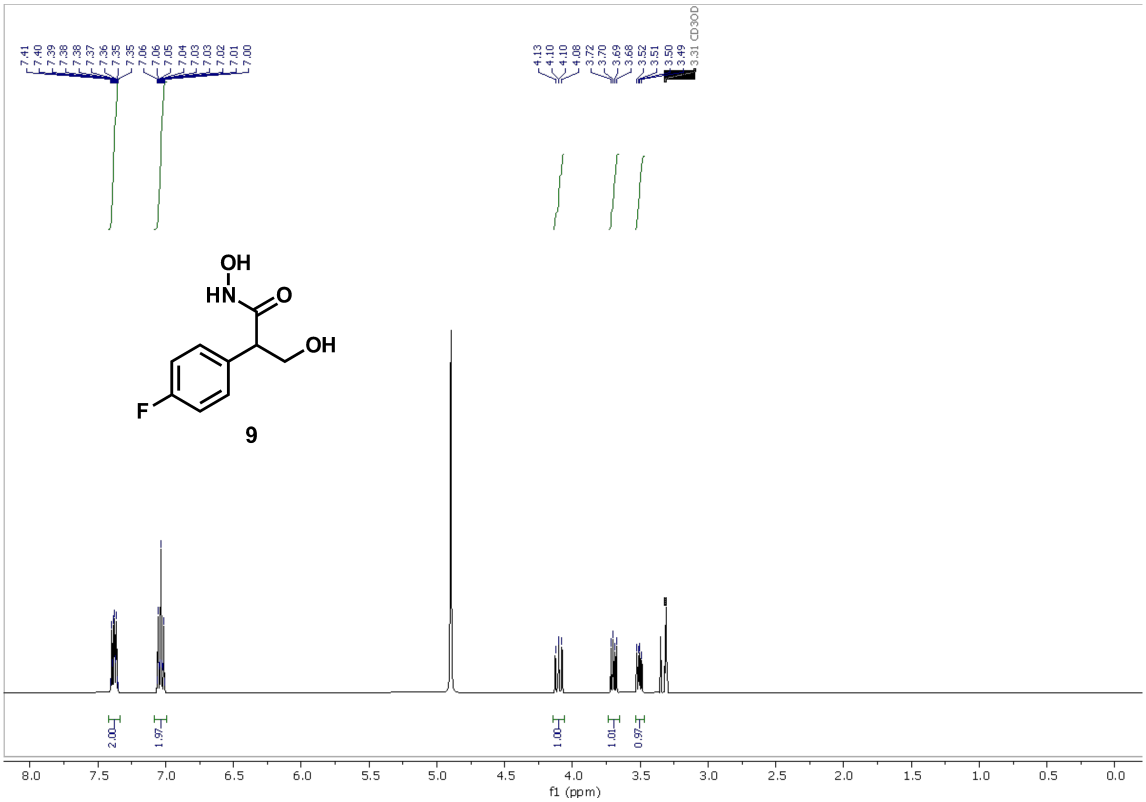
7.41
7.40
7.39
7.38
7.38
7.37
7.36
7.35
7.35
7.06
7.06
7.05
7.04
7.03
7.03
7.02
7.01
7.00
4.13
4.10
4.10
4.08
3.72
3.70
3.69
3.68
3.52
3.51
3.50
3.49
3.31 CD3OD
OH
HN
O
OH
F
9
2.00
1.97
1.00
1.01
0.97
f1 (ppm)

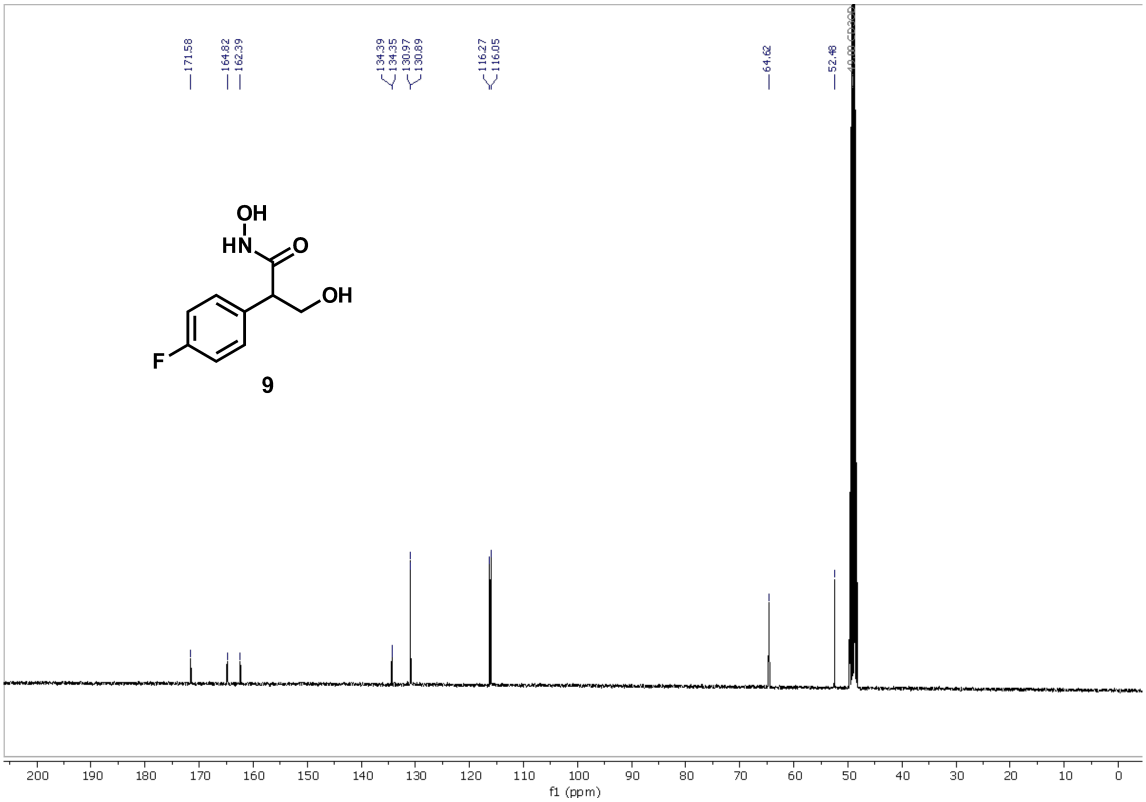
171.58
164.82
162.39
134.39
134.35
130.97
130.89
116.27
116.05
64.62
52.48
OH
HN
O
OH
F
9
f1 (ppm)

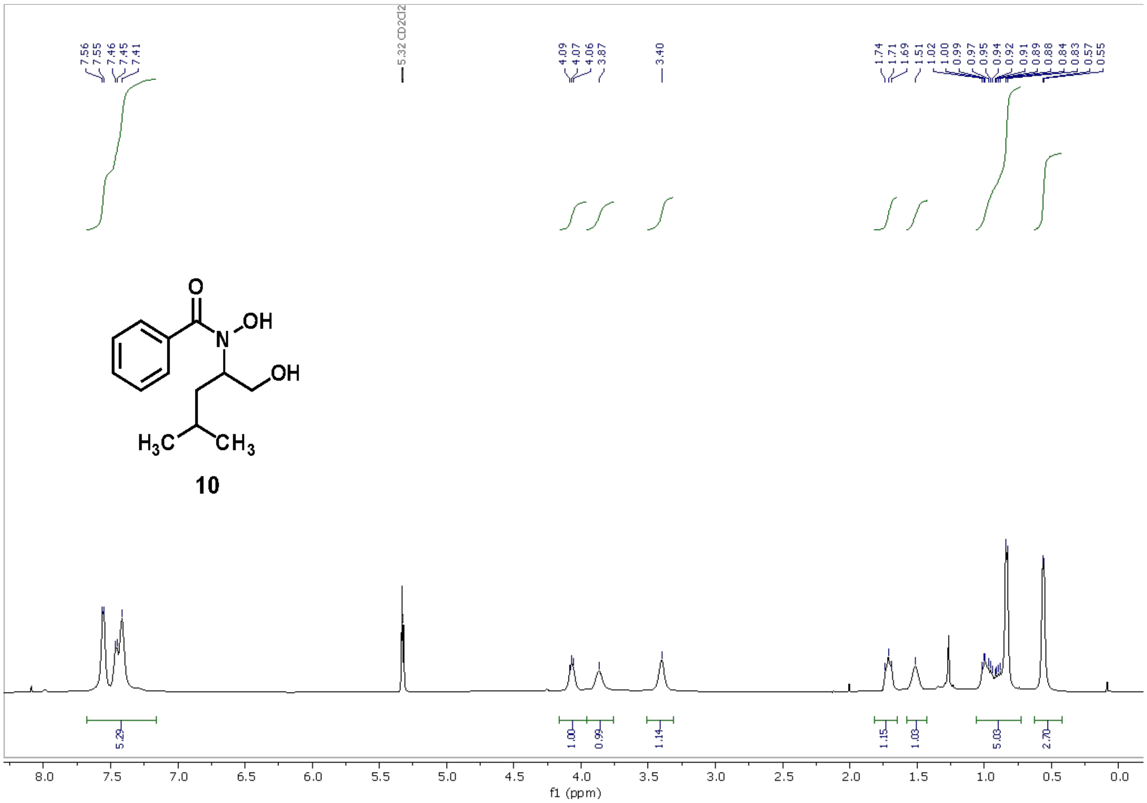
7.56
7.55
7.46
7.45
7.41
5.32 CD2Cl2
4.09
4.07
4.06
3.87
3.40
1.74
1.71
1.69
1.51
1.02
1.00
0.99
0.97
0.95
0.94
0.92
0.91
0.89
0.88
0.84
0.83
0.57
0.55
OH
N
O
OH
H3C
CH3
10
5.29
1.00
0.99
1.14
1.15
1.03
5.03
2.70
f1 (ppm)

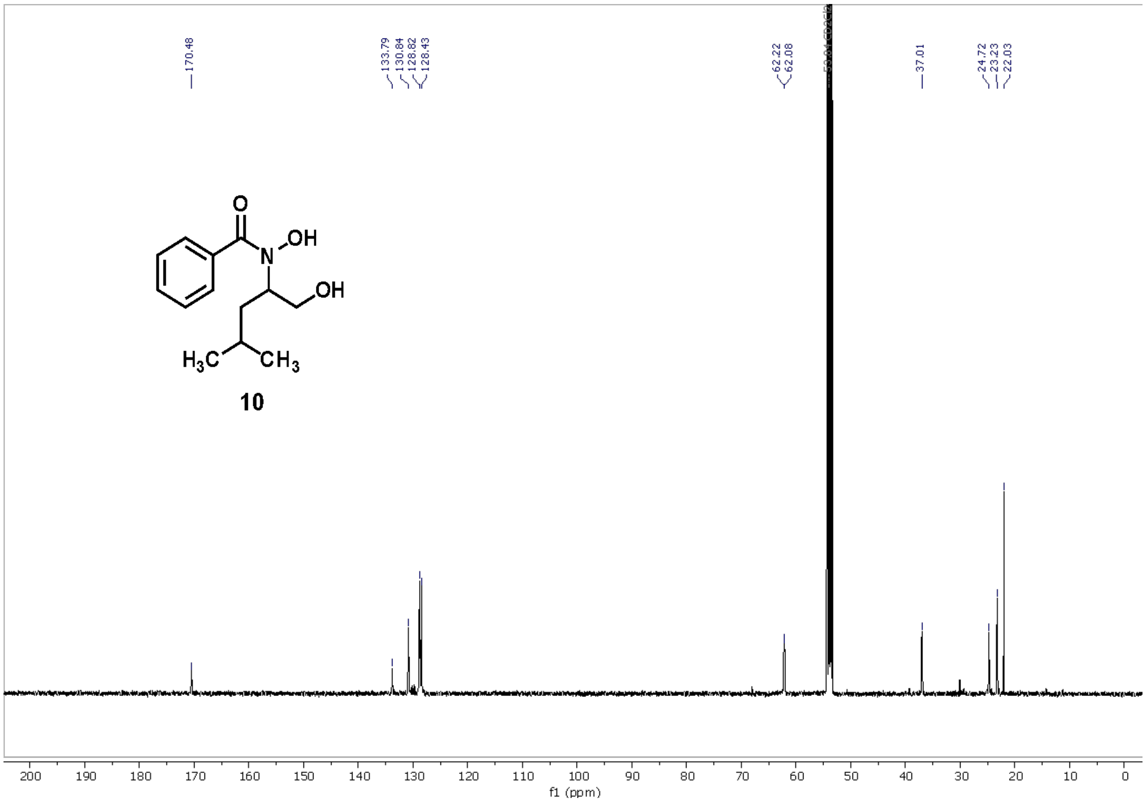
170.48
133.79
130.84
128.82
128.43
62.22
62.08
37.01
24.72
23.23
22.03
O
N
OH
OH
H3C
CH3
10
f1 (ppm)

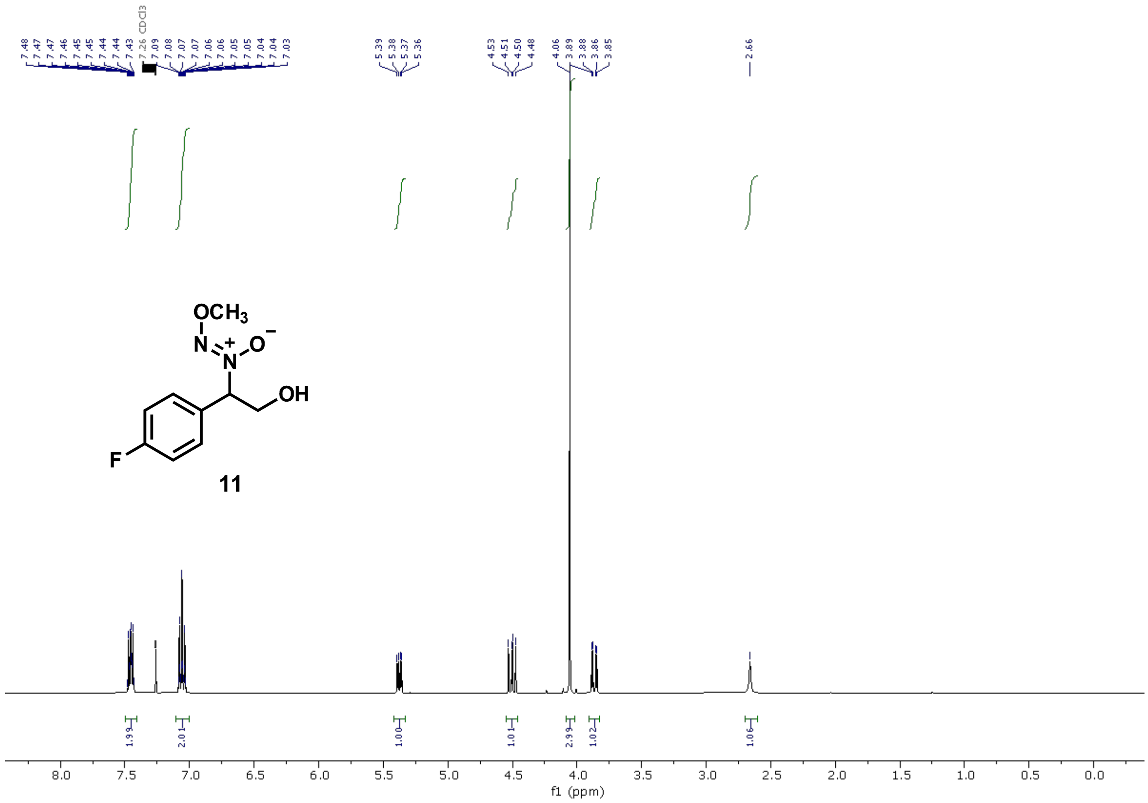
7.48 7.47 7.46 7.45 7.45 7.44 7.44 7.43 7.26 CDCl3 7.09 7.08 7.07 7.07 7.06 7.06 7.05 7.05 7.04 7.04 7.03
5.39 5.38 5.37 5.36
4.53 4.51 4.50 4.48
4.06 3.89 3.88 3.86 3.85
2.66
OCH3
N
N
O
OH
F
11
1.99 2.01 1.00 1.01 2.99 1.02 1.06
8.0 7.5 7.0 6.5 6.0 5.5 5.0 4.5 4.0 3.5 3.0 2.5 2.0 1.5 1.0 0.5 0.0
f1 (ppm)

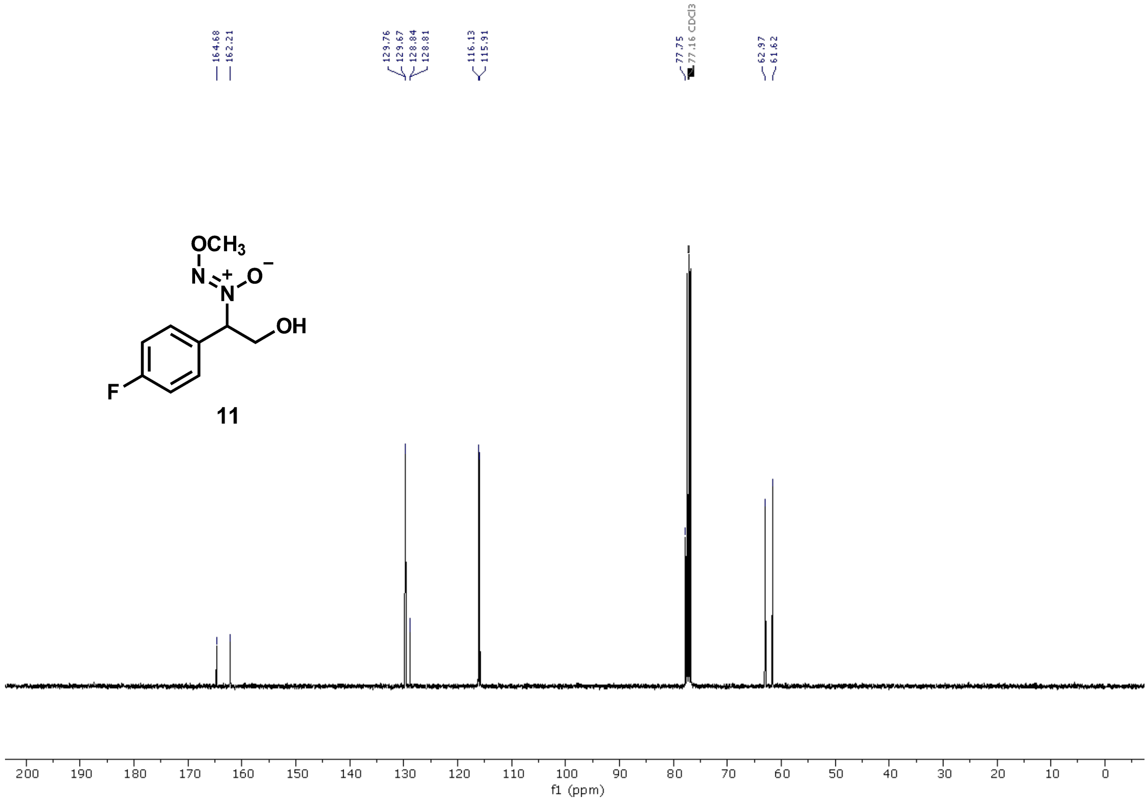
164.68 162.21
129.76 129.67 128.84 128.81
116.13 115.91
77.75 77.16 CDCl3
62.97 61.62
OCH3
N
N
O
OH
F
11
200 190 180 170 160 150 140 130 120 110 100 90 80 70 60 50 40 30 20 10 0
f1 (ppm)

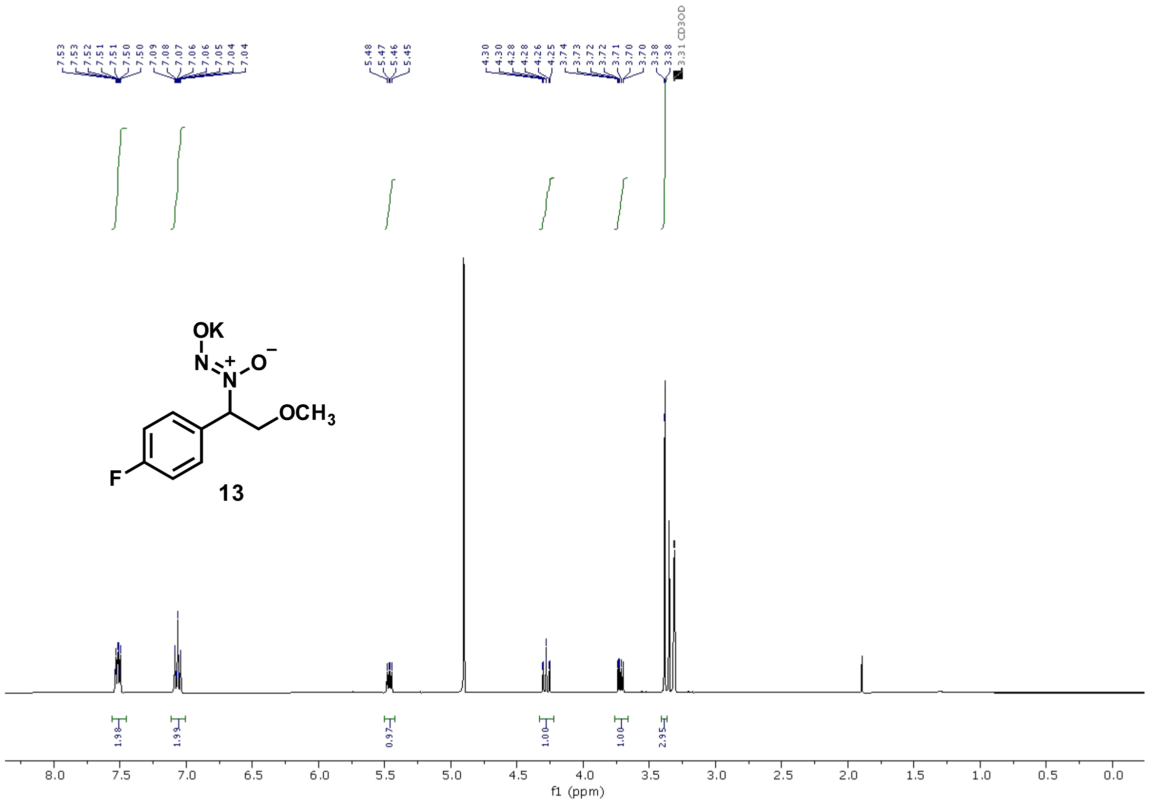

OK
N
O
OCH3
F
13
f1 (ppm)


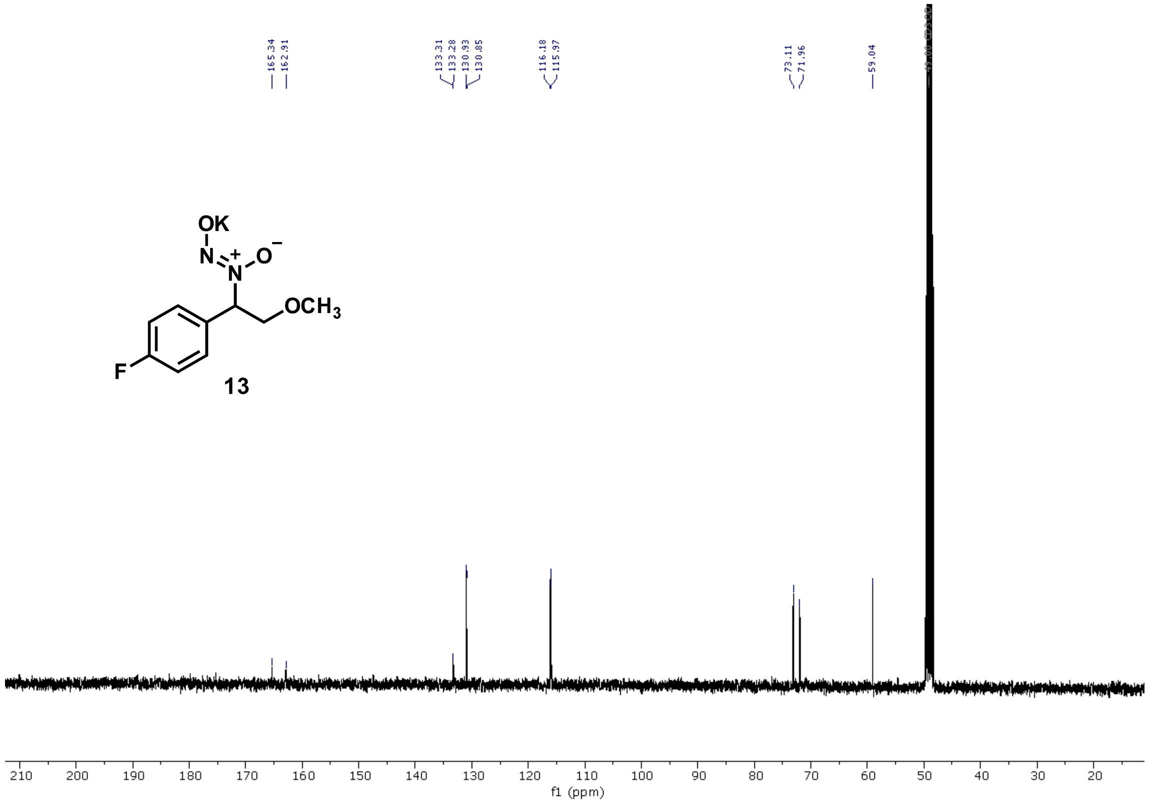

OK
N
O
OCH3
F
13
f1 (ppm)

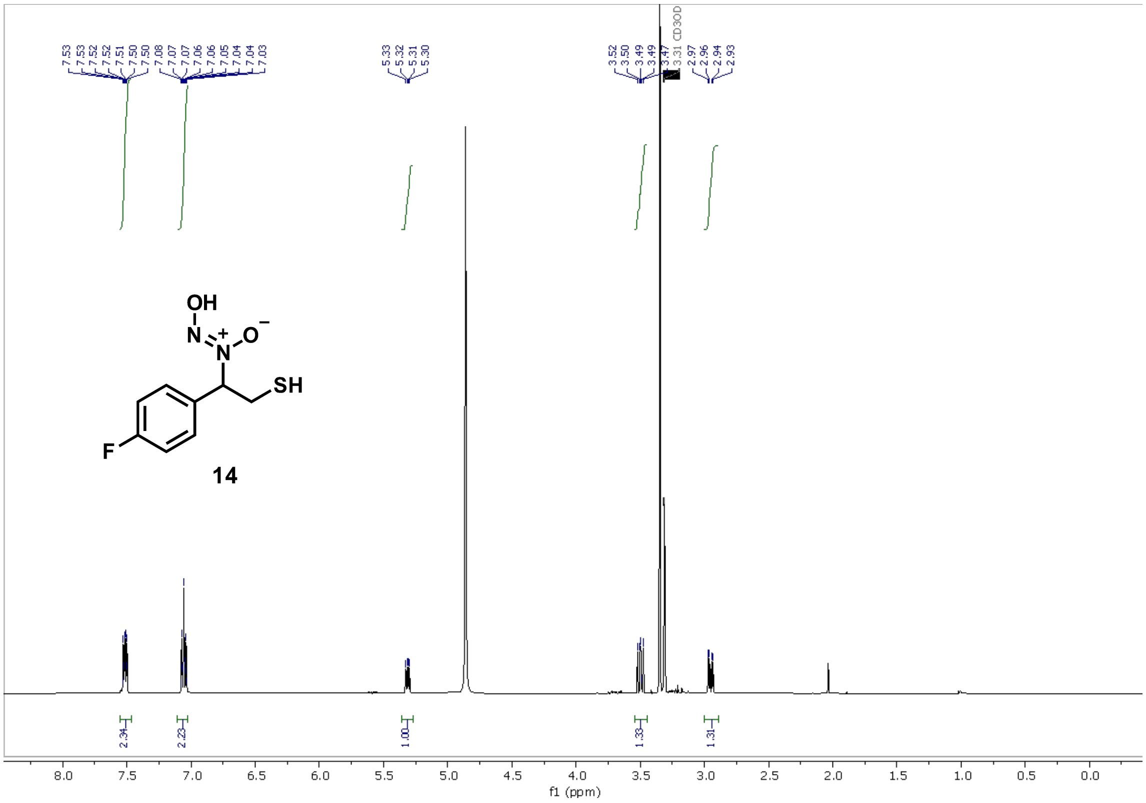
7.53
7.53
7.52
7.51
7.50
7.50
7.08
7.07
7.07
7.06
7.06
7.05
7.04
7.04
7.03
5.33
5.32
5.31
5.30
3.52
3.50
3.49
3.49
3.47
3.31 CD3OD
2.97
2.96
2.94
2.93
OH
N
O
SH
F
14
2.34
2.23
1.00
1.33
1.31
8.0
7.5
7.0
6.5
6.0
5.5
5.0
4.5
4.0
3.5
3.0
2.5
2.0
1.5
1.0
0.5
0.0
f1 (ppm)

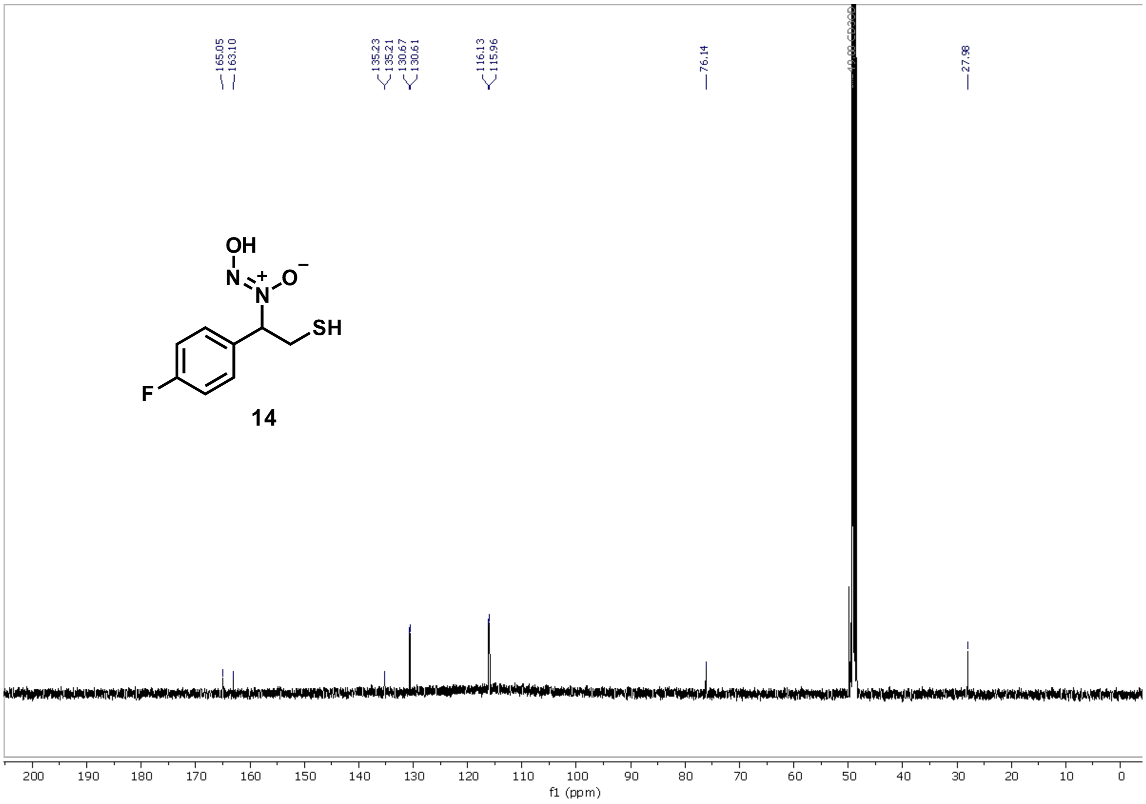
165.05
163.10
135.23
135.21
130.67
130.61
116.13
115.96
76.14
27.98
OH
N
O
SH
F
14
200
190
180
170
160
150
140
130
120
110
100
90
80
70
60
50
40
30
20
10
0
f1 (ppm)

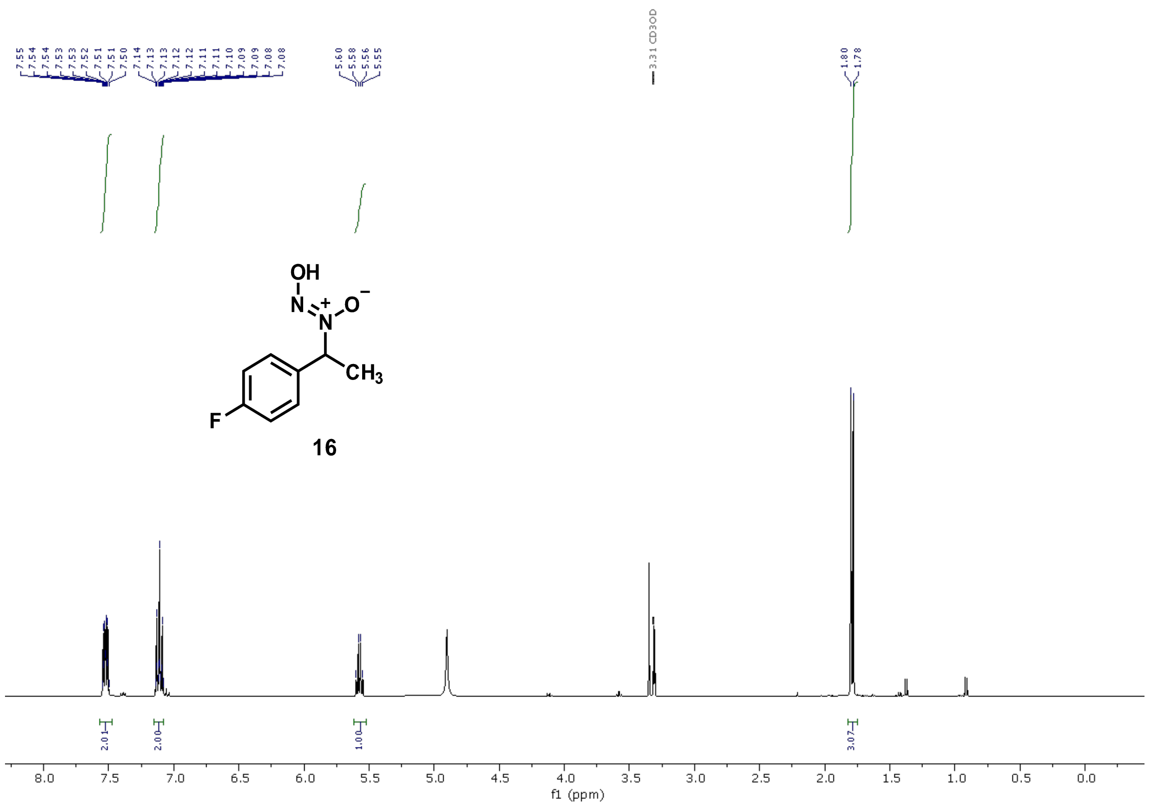

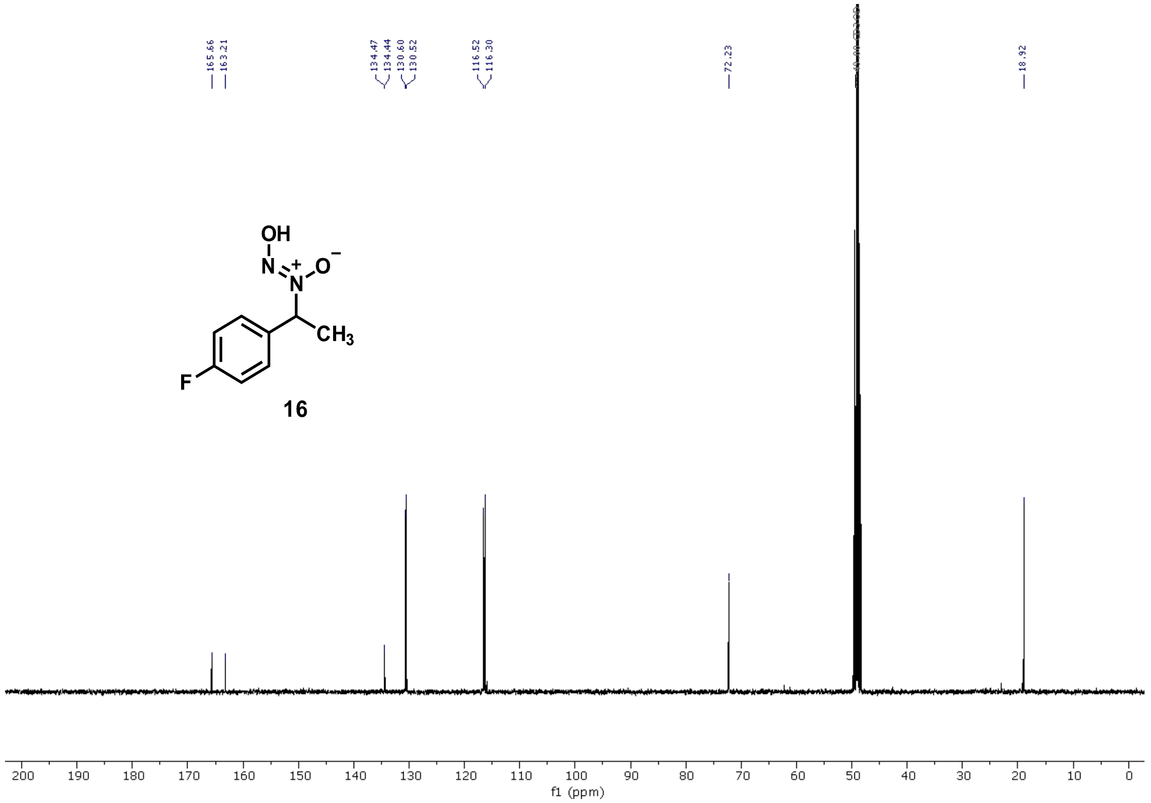

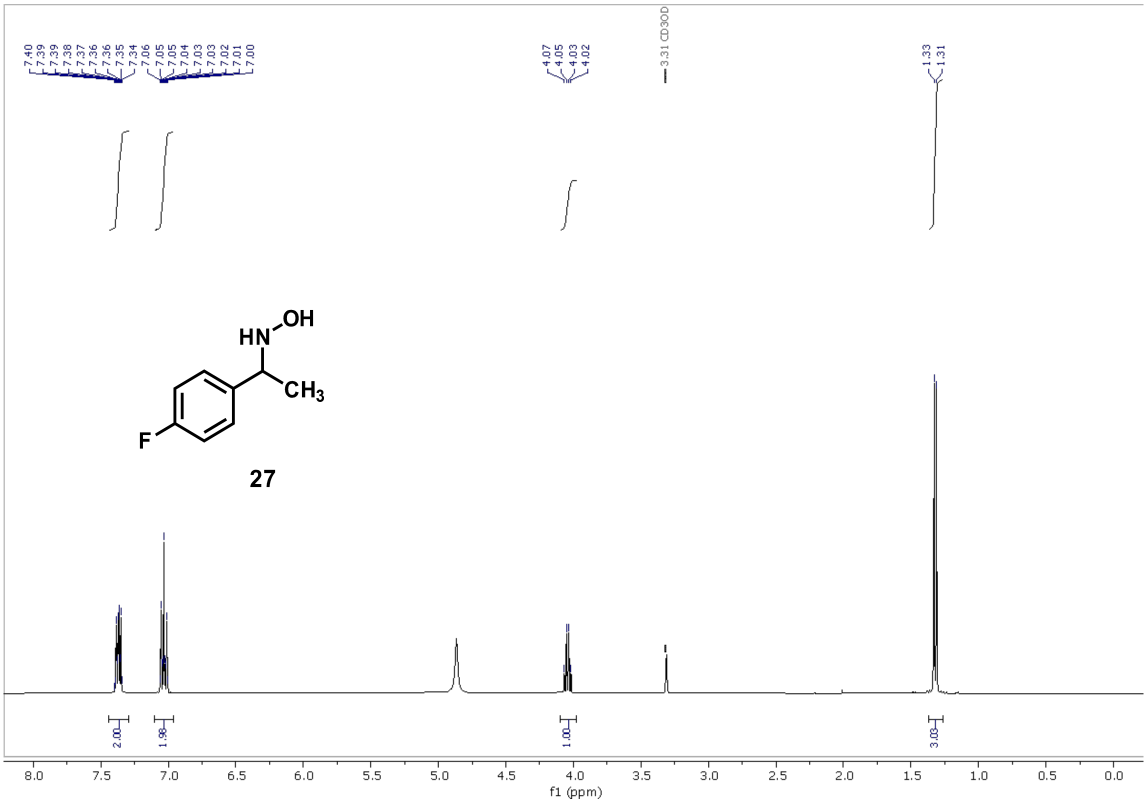
7.40 7.39 7.39 7.38 7.37 7.36 7.36 7.35 7.34 7.06 7.05 7.05 7.04 7.03 7.03 7.02 7.01 7.00
4.07 4.05 4.03 4.02
3.31 CD3OD
1.33 1.31
HN
OH
CH3
F
27
2.00
1.98
1.00
3.03
8.0 7.5 7.0 6.5 6.0 5.5 5.0 4.5 4.0 3.5 3.0 2.5 2.0 1.5 1.0 0.5 0.0
f1 (ppm)

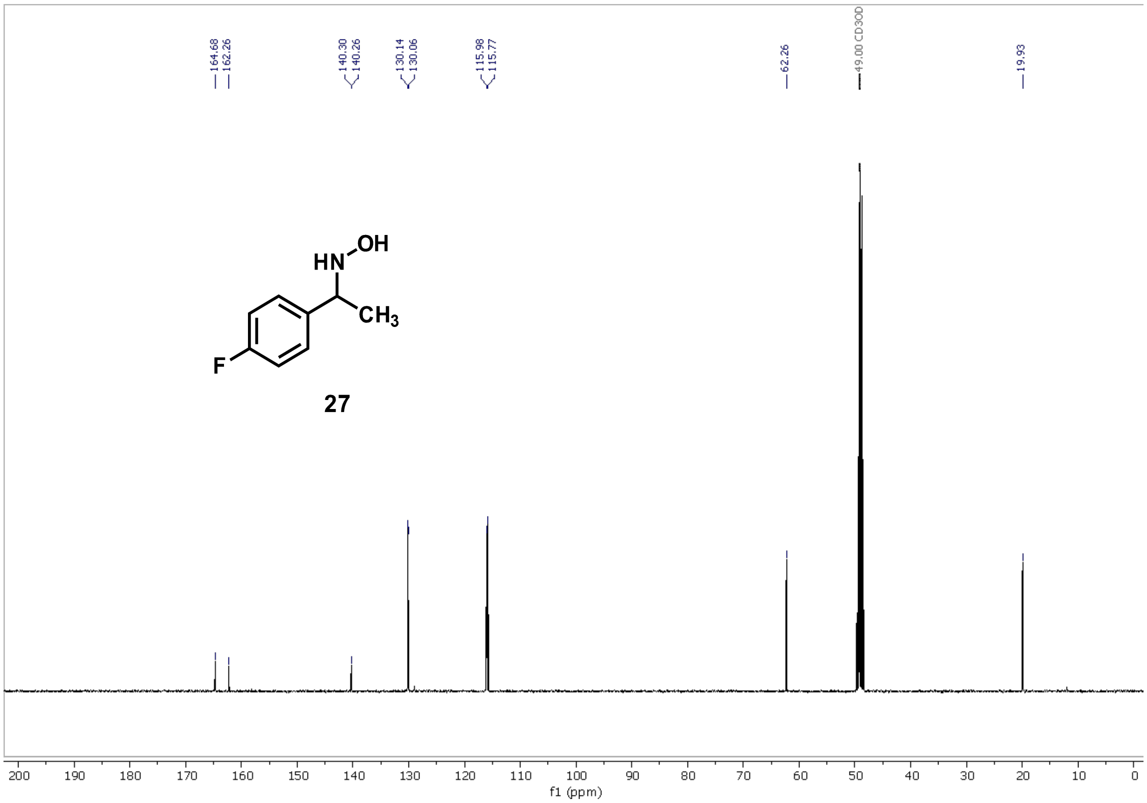
164.68 162.26
140.30 140.26
130.14 130.06
115.98 115.77
62.26
49.00 CD3OD
19.93
HN
OH
CH3
F
27
200 190 180 170 160 150 140 130 120 110 100 90 80 70 60 50 40 30 20 10 0
f1 (ppm)

**Evaluation of the antagonistic activity of the leudiazen derivatives by beta galactosidase assay**

A previous study showed that mangotoxin promoter is induced by leudiazen.[5] A transcriptional lacZ fusion of the mangotoxin promoter in the Pss strain was used as a sensor and the promoter activity of this locus was assessed by galactosidase assays[5] to identify the analogues competitively inhibiting leudiazen. Bacterial cells were grown overnight in PMS (Pseudomonas minimal medium) medium in the presence of 20 µM synthetic leudiazen or different leudiazen derivatives dissolved in methanol. Bacterial overnight cultures were centrifuged at 16,000 rpm for 5 minutes, resuspended in Z-buffer and $OD_{600}$ (optical density at 600 nm) values were recorded. To permeabilize the cell membrane, 25 µl of chloroform and 0.1 % SDS were added to the residual 1 ml of bacterial suspension, vortexed for 10 seconds and incubated at 28 °C for 10 minutes. 200 µl of o-nitrophenyl-β-D-galactoside (ONPG) solution (4 mg/ml in Z-buffer) were added to each sample to initiate the reaction, vortexed briefly and incubated at room temperature. The reaction was stopped by adding 500 µl 1 M $Na_2CO_3$ after the development of a satisfactory yellow color, or after 30 minutes in the absence of a reaction. The samples were centrifuged at 13,000 rpm for 10 minutes and 1 ml of cell-debris free supernatant was used to measure the absorbance at 420 nm and 550 nm. For every β-galactosidase assay, a sample containing only the growth medium was processed as described above and used as a blank.
To determine the promoter activity in the sample, Miller Units were calculated using the following formula:

1 Miller Unit = $1000 * [(OD_{420} - 1.75 * OD_{550})] / (t * v * OD_{600})$

t = reaction time in minutes; v = volume of assayed sample in mL

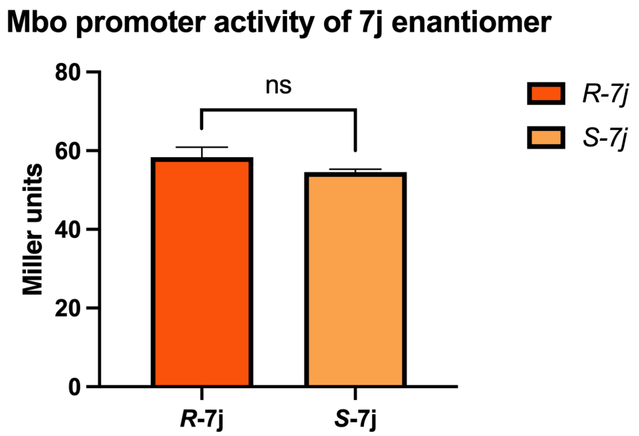


**Figure S1. Mbo promoter comparison activity of *R* and *S* 7j.** Activities of the mboA promoter in the presence of both enantiomers of **7j** (20 µM). The error bars represent the SEM and the analysis was performed using the t-test (the calculated p value was 0.22).

Virulence assays

**Mangotoxin production assay using *E. coli* as an indicator strain**

Mangotoxin production was determined using an *E. coli* strain as an indicator using a previously described procedure[5] with minor modifications. A layer of the indicator microorganism, an *E. coli* K12 strain was prepared, and after solidification, the Pss strains (WT and Δmgo mutant) were stab-inoculated on to the agar seeded with *E. coli*. In the antagonism assays, 20 μM leudiazen or their derivatives to be tested were added in to the medium. All the compounds were tested for any inhibitory effect on the growth of the *E. coli* strain before the assay. The plates were incubated at 30°C for 48 hours and the inhibition zones around the colony were measured.

**Virulence assay using tomato leaflets**

Virulence experiments were performed on detached tomato leaflets (*Solanum lycopersicum* Mill.) of Hellfrucht Frühstamm variety as previously described[5] with slight modifications. Six tomato leaflets were used per strain.

1. **Infection**. Each leaflet was disinfected with 0.1% sodium hypochlorite, washed and air-dried. Three 10 μl drops of the bacterial suspension (exponentially growing cultures of Pss in PMS adjusted to $10^8$ CFU $ml^{-1}$) were injected using a sterile entomological needle at different points on one side, while three 10 μl drops of aqueous $MgSO_4$ solution (10 mM) were injected on the other side of the same leaflet as a control.
2. **Treatment**. Some of the leaflets (6) infected with Pss were washed in aqueous soln. of compound **14** or leudiazen (20 μM) with manual shaking at RT, allowing a total contact time of around 20 seconds. The solution was decanted, and the leaves were then dipped in fresh sterile distilled water to remove any traces of compounds.

All infected and infected-then-treated leaflets were maintained at 22°C and a 16:8-hour light:dark photoperiod. The development of necrotic symptoms was determined over a period of 10 days. Digital pictures of the leaflets were used to determine the necrotic area using the ImageJ software. The total leaf area and the infected areas were defined manually on each leaflet and the relative virulence was calculated as (total lesion area/total leaf area) × 100. Bacterial strains were recovered from the leaflets after 10 days and the presence of Pss was confirmed by PCR using Pss specific primers.
